# Evaluating the practical aspects and performance of commercial single-cell RNA sequencing technologies

**DOI:** 10.1101/2025.05.19.654974

**Authors:** Anna E. Elz, Derrik Gratz, Annalyssa Long, David Sowerby, Azi Hadadianpour, Evan W. Newell

**Author notes:** Corresponding author: Anna Elz.

## Abstract

The rapid development of updated and new commercially available single-cell transcriptomics platforms provides users with a range of experimental options. Cost, sensitivity, throughput, flexibility, and ease of use, all influence the selection of an optimal workflow. We performed a comprehensive comparison of single-cell transcriptomic approaches using multiple standardized PBMCs from different donors. We report on standard single-cell metrics including cell recovery, sequencing efficiency, sensitivity, cell annotation, and differential gene expression for seven recently available kits that interrogate whole transcriptome mRNA, and two that include TCR profiling. We also discuss workflow throughput, sample requirements, timing, cost, and labor as critical factors to consider. In addition to variable experimental constraints imposed by each platform, our findings highlight differences in cell recovery and sensitivity, which we found to significantly influence the ability to resolve cell subtypes. This work provides a basis by which users can balance performance and practical considerations when selecting a single-cell RNA sequencing platform.

## Introduction

Single-cell sequencing technologies have revolutionized our understanding of cellular diversity and heterogeneity, providing unprecedented insights into biological processes at the individual cell level [1]. Over the past decade, as single-cell technologies have become increasingly mainstream for profiling the transcriptomes of cells from culture, blood, tissues, and organisms, additional options for single-cell RNA gene expression analysis have emerged. Studies focused on evaluating the performance of different platforms continue to guide choices in research projects as platforms evolve [2], [3], [4], [5]. In turn, we have performed an extensive comparative study to assess the performance and logistical characteristics of current market technologies.

To perform single-cell analysis, individual cells must first be partitioned. In addition to various other approaches, this can be achieved by capturing cells in oil-based droplets, that act as individual chambers, or by multiple rounds of cell tagging where the cell serves as the reaction chamber. Commercial droplet-based systems encapsulate and barcode single cells in a partitioning oil either by microfluidics to create Gel Bead-in-Emulsions (GEMs, 10x Genomics) or by vortexed particle-templated instant partitions (PIPs, Fluent Biosciences / Illumina) [6] . Each encapsulated cell undergoes lysis, releasing mRNA that is reverse-transcribed with poly-dT barcoded primers. cDNA molecules in GEMs and v4 PIPs receive a cell barcode and unique molecular identifier (UMI), and cDNA products are used to generate sequencing libraries. Fluent V updated the approach to counting molecules by generating Intrinsic Molecular Identifiers from unique cut sites in the fragmented cDNA [7]. Plate-based approaches utilize a “split and pool” approach performed with ligation [8] or tagmentation [9], [10] beginning with individual sample tags on the first plate and a series of pooling and splitting samples across two additional plates to generating unique tag combinations to identify individual cells. This approach requires fixed and permeabilized cells which serve as the reaction compartments for lysis and reverse transcription (RT). The recovery of mRNA transcripts differs between the two commercially available platforms we tested. Parse Biosciences uses both a poly(T) primer and random hexamers to capture the full length of the transcript and ligates barcodes to tag cells. Scale Biosciences utilizes a poly(T) primer approach to capture the 3’ end of the transcript and employs longer indices to tag cells. At the library stage, samples are either sequenced in a set of sublibraries (Parse) or as a set of 96 pooled libraries (Scale).

Multiple comparative studies have examined the sensitivity and cost differences across various technologies [11], [12], [13], but with the recent addition of new platforms and updated chemistries, further comparative studies are necessary. The increasing diversity of available technologies means that a range of practical factors should be considered when selecting a kit tailored for specific experimental goals or user constraints. For instance, cost may hinder the breadth of experiment designs, labor requirements can impact the feasibility of some workflows, limited throughput can increase batch effects, and cell input requirements all contribute to the platform selection process. Other factors, such as the ability to fix samples soon after collection, scalability, and protocol flexibility, should also be examined before deciding on a platform for individual use or as a core service offering. Downstream analysis is also important to consider; data quality, quantity, and reproducibility dictate the utility of each platform. Lastly, while other comparative studies have normalized for cell recovery, reference genome, and QC thresholds (De Simone et al., 2025; Filippov et al., 2024; Xie et al., 2024), they lack a practical comparison of each kit’s full data recovery yielded from recommended pipeline settings.

To address this gap, we evaluated the latest offerings from widely used commercial single-cell sequencing platforms described above and compared their performance to earlier versions. Following recommended protocols for each platform, we generated consistent metric definitions across each platform to enable a fair and practical comparison of the key features and experimental considerations of each. This comparison consisted of two parallel studies. One study evaluated seven different single-cell platforms that generate 3’ or whole transcriptome (WT) data. In the other study, we compared two platforms that generate 5’ WT data in addition to linked single-cell TCR sequences from the same single cells. Based on these studies we comment on a wide range of factors to consider when planning single-cell sequencing experiments.

## Results

We performed two benchmarking experiments to evaluate recently released kits and provide comparisons to earlier chemistry. We tested several barcoding platforms with differing technological approaches including disposable microfluidic chips and a benchtop instrument (10x Genomics Chromium X), random emulsion with specialized equipment (Fluent Biosciences PipSeq Vortexer and Dry Bath Incubator), and split-pool synthesis plate-based approaches (Parse Biosciences and Scale Biosciences). Our goal was to utilize each kit’s maximum capacity to compare the practical considerations of each more fairly. We used frozen aliquots of healthy donor PBMCs in two separate experiments (See Methods); one tested reproducibility using two samples with replicates across seven scRNA platforms (Figure 1a), and the second profiled the TCR repertoire of four donor’s sorted T-cells with three assays (Supplemental Figure 1a). Each technology was assessed for workflow ease of use and flexibility, cell recovery, cost-effectiveness, sequencing efficiency, gene and transcript detection sensitivity, reproducibility, and the resolution of cellular subsets, including the sensitivity to detect differences in gene expression. These experiments also enabled us to comment comprehensively on the user experience as a major consideration in the choice of kits for single-cell experiments.

**Figure 1:**
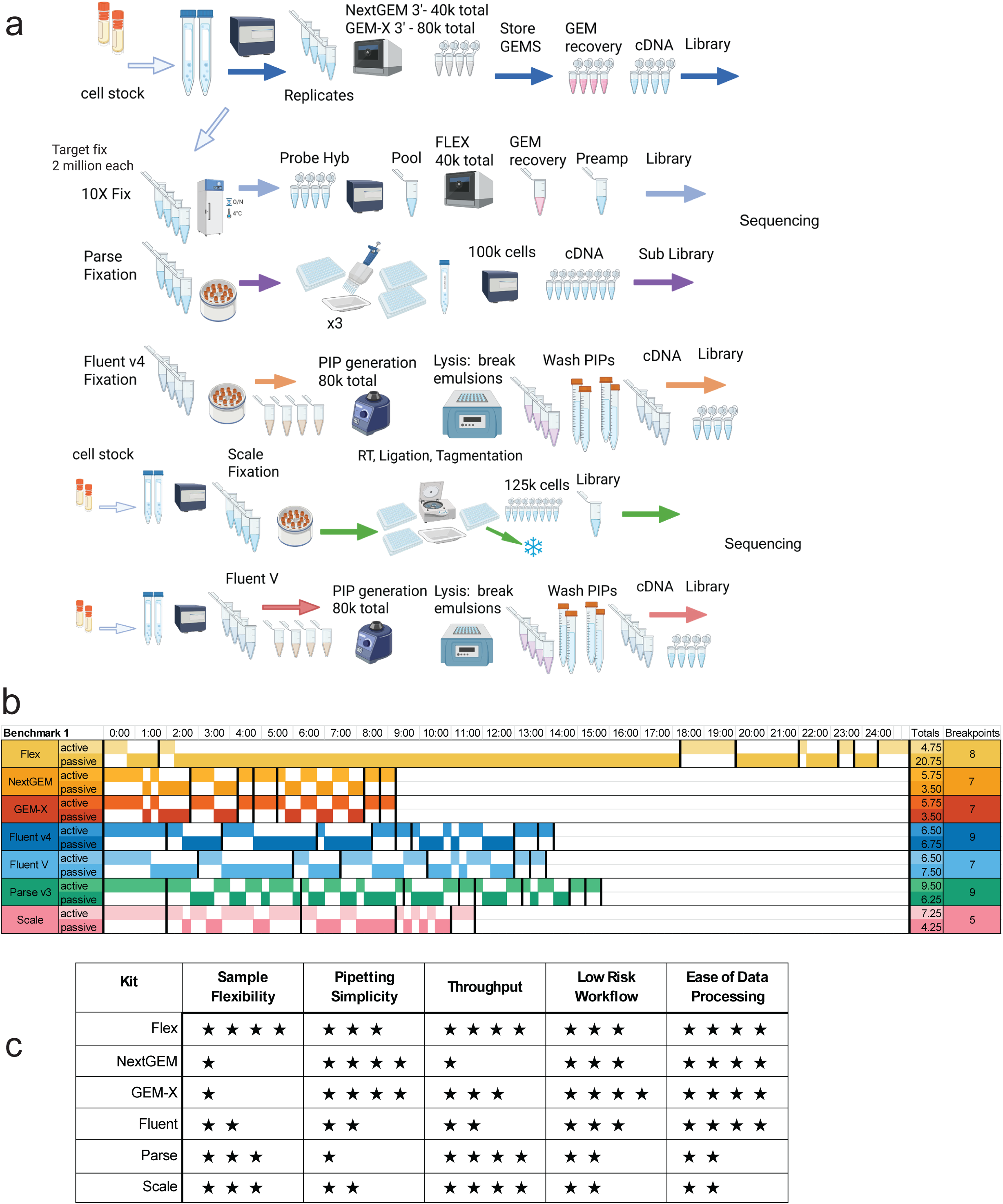
Description of the experimental workflow, processing steps, and experience a. Workflow diagram of different steps required for each platform used in the Benchmark 1 experiment from cell suspension to library sequencing. 10x NextGEM Flex, NextGEM 3’, GEM-X 3’, Fluent v4, and Parse v3 cells were from the sample aliquot. Scale and Fluent V processing occurred separately with additional aliquots of the same donor. b. Summary of time required to execute each workflow from sample preparation to library completion in terms of hours spent hands-on (active) and time spent waiting (passive). Stopping points are indicated with red lines, and the number of stopping points is summarized in the rightmost column. Gray indicates an overnight stopping point. c. Workflow processing experience summary table classifies each criterion from low (one star) to high (four stars). Sample flexibility refers to the option to fix and store samples prior to beginning the single-cell workflow. Pipetting refers to the number of pipetting steps. Throughput refers to the parallel processing of samples enabled by multiplexing. Low Risk refers to an ability to salvage an experiment from an intermediate stage or recover partial inputs. Data processing refers to the ease of use for each vendor’s software.

### Study Design

For our first experiment, Benchmark 1 (BM1), we used two healthy PBMC donor samples in replicate (F1A,B; F2A,B), to compare seven platforms: 10x Genomics’ Chromium NextGEM 3ʹ v3.1 (labeled “NextGEM 3’” in figures and results), GEM-X 3’ v4 (GEM-X 3’), and Flex NextGEM Fixed RNA Profiling (Flex), Parse’s Evercode WT v.3 (Parse v3), Scale Biosciences Single Cell RNA kit v1.1 (Scale), and Fluent Biosciences PIP Seq T20 3ʹ v4 (Fluent v4) and PIPseq V T20 3’ (Fluent V). All platforms were run in parallel on the same day except for Scale and Fluent PIPseq V assays, which were processed separately using aliquots from the same donors’ leukapheresis samples (Figure 1a).

For our second experiment, Benchmark 2 (BM2), we compared platforms that capture full-length TCR sequences from single cells (Supplemental Figure 1a). For this, we used flow cytometry to sort T-cells based on CD3 expression from 4 different healthy donor PBMCs (F1, F2, F4, F5) to run on 10x Genomics Chromium NextGEM 5ʹ v2 (NextGEM 5ʹ), GEM-X 5ʹ v3 (GEM-X 5’) and Parse EvercodeTCR (Parse v2, TCR). Two sets of samples were processed on separate days. A workflow error in the second NextGEM 5’ run was evident and produced poor-quality cDNA for two of the four samples (Supplemental Figure 1c). We removed these samples from the analysis.

In each study, we targeted the recommended maximum cell recovery of each kit (Table 1). Sample fixation for Parse and Fluent v4 was performed at the same time cells were loaded into the Chromium chips for the 10x Genomics platform. To reduce the influence of variable sequencing depth in our comparisons across technologies, we downsampled the FASTQ files before passing data through their respective pipelines with default settings (see “Sequencing Depth and Downsampling” and “Generating Count Matrices” in Methods).

**Table 1.** Platform metrics and descriptions.

| Assay details |  |  |  |  |  | Sequencing read length (nt) |  |  |  | Cell recovery |  | Cost |  |  |  |
| --- | --- | --- | --- | --- | --- | --- | --- | --- | --- | --- | --- | --- | --- | --- | --- |
| Protocol | Kit ID | Assay type | Sample type | Sample process | Detection method | Read 1 | Index 1 | Index 2 | Read 2 | Cell target per sample | Recommended cell loading | Instrument cost | Kit cost | Kit cost per sample (n= 4)<sup>1</sup> | Kit cost per cell<sup>1</sup> |
| Benchmark 1 |  |  |  |  |  |  |  |  |  |  |  |  |  |  |  |
| 10x Genomics Chromium Fixed RNA Profiling Kit, Multiplex 4 x 4 | Flex | Emulsion | PBMCs | Formalin Fixed | Probe Hybridization | 28 | 10 | 10 | 90 | 10,000 | 17,000 | \$75,000 | \$15,240 | \$952 | \$0.10 |
| 10x Genomics Chromium NextGEM 3' v3.1 4rxn | NextGEM 3' | Emulsion | PBMCs | Not fixed | Reverse Transcription 3' end | 28 | 10 | 10 | 90 | 10,000 | 17,000 | \$65,000 | \$7,840 | \$1,960 | \$0.20 |
| 10x Genomics Chromium GEM-X 3' v4 4rxn | GEMX 3' | Emulsion | PBMCs | Not fixed | Reverse Transcription 3' end | 28 | 10 | 10 | 90 | 20,000 | 29,000 | \$75,000 | \$6,811 | \$1,703 | \$0.10 |
| Fluent Biosciences PIPSeq T-20 v4 | Fluent v4 | Emulsion | PBMCs | DSP/Methanol Fix | Reverse Transcription 3' end | 54 | 8 | 8 | 67 | 20,000 | 40,000 | \$3,000 | \$3,600 | \$900 | \$0.05 |
| Fluent Biosciences PIPSeq T-20 V | Fluent V | Emulsion | PBMCs | Not fixed | Reverse Transcription 3' end | 45 | 10 | 10 | 72 | 20,000 | 40,000 | \$3,000 | \$3,600 | \$900 | \$0.05 |
| Parse Evercode v3 | Parse v3 | Combinatorial Indexing | PBMCs | Fixed Permeabilized | Reverse Transcription 3' end | 64 | 8 | 8 | 58 | 25,000 | 350,000 | NA | \$10,780 | \$2,695 | \$0.11 |
| Scale Biosciences | Scale | Combinatorial Indexing | PBMCs | DSP/Methanol Fix | Reverse Transcription full | 34 | 10 | 10 | 76 | 31,250 | 960,000 | NA | \$8,500 | \$2,125 | \$0.07 |
| Benchmark 2 |  |  |  |  |  |  |  |  |  |  |  |  |  |  |  |
| 10x Genomics Chromium NextGEM 5' v2 | NextGEM 5' | Emulsion | T-cells | Not fixed | Reverse Transcription 5' end | 26 | 10 | 10 | 90 | 10,000 | 17,000 | \$75,000 | \$7,840 | \$1,960 | \$0.20 |
| 10x Genomics Chromium GEM-X 5' v3 | GEMX 5' | Emulsion | T-cells | Not fixed | Reverse Transcription 5' end | 28 | 10 | 10 | 90 | 20,000 | 29,000 | \$65,000 | \$6,811 | \$1,703 | \$0.10 |
| Parse Evercode TCR | Parse TCR | Combinatorial Indexing | T-cells | Fixed Permeabilized | Reverse Transcription full | WT: 66;<br>TCR: 216 | 8 | 8 | WT: 86;<br>TCR: 86 | 25,000 | 350,000 | NA | \$12,500 | \$3,125 | \$0.13 |

<sup>1</sup> Does not included indices

### Sample Processing

The amount of hands-on vs. hands-off time varied across platforms and is reported based on published user guides aligned with our experience. Protocol duration began with standard cell preparation steps to the first stop point (fixation or emulsion) (Figure 1b). Flex had the shortest hands-on time, but the overnight hybridization increased time to completion while also increasing sample preparation flexibility since the samples can be frozen after this step. The new GEM-X 3’ and NextGEM 3’ kits had nearly identical protocols and provided the fastest processing time with shorter intervals between stop points. The two combinatorial barcoding platforms’ pooling approaches differed, with Scale’s use of a centrifugal plate funnel, rather than pipette transfers, resulting in less hands-on time, but longer no-stop intervals. The Parse protocols required more hands-on time due to multiple bead-clean steps after sublibrary generation, while Scale utilized stop enzymes and a single bead clean-up step for the final indexed library. Fluent had more hands-on time in comparison to the other emulsion assays and a longer overall protocol due to the incubation steps. With the addition of the TCR profiling, each platform in the BM2 experiment had expectedly longer workflows than their WT counterparts due to the additional TCR library preparation and pipette-based mixing for Parse v2 chemistry (Supplemental Figure 1b).

Based on these experiments, we subjectively assessed the processing experience from least favorable (*) to most favorable (****) based on sample flexibility (ability to bank samples), the quantity of pipetting required, throughput potential, the risk for experiment or sample failure, and the ease of the initial data processing (Figure 1c, see criteria in “Processing Comparison” in Methods). Sample flexibility considered the fixation process, storage time, and cell range requirements. Flex rated highest with formalin fixation for up to 10 million cells per sample and multiple stages to bank samples. The Parse and Scale‘s fixation process involved more steps and a smaller input range per sample. Fluent’s fixation was optional, and cells could only be stored for a week. Parse required the most pipetting followed by Scale, which reduced liquid transfer steps with centrifugal pooling and split cells into a 384-well plate. While we processed only 4 samples per kit, throughput refers to the total possible number of samples or cells from the kits used in these experiments. With multiplexing, Flex can deliver the most cells per sample and up to 160k cells total. Both Parse and Scale allowed for increased sample throughput, 24 and 48 respectively, which decreases cell recovery for each additional sample. GEM-X doubled the cell throughput of NextGEM and matched Fluent’s but ranked higher in sample throughput due to the ease of processing. We found a lower risk potential in the ability to process individual samples, as seen in our NextGEM 5’ loss. (Supplemental Figure 1c). Improved chip fluidics lowered the risk for GEM-X. The “all-in” approach required by combinatorial plate assays was deemed a higher risk since experimental loss was possible with an error at almost any step. Recognizing that the experience and risk level of any platform will vary across users, our assessment is provided as a general guideline.

For data processing, 10x’s Cellranger and Fluent’s PIPseeker stood out as standalone pipelines that were installed easily, executed without issue, provided intermediate files for deeper introspection, and had comprehensive, easily interpretable output summaries. Parse’s ‘TrailMaker’ is a cloud-based pipeline accessed through a web browser. Trailmaker does not provide sample-level FASTQs or BAMs, limiting the addition of accessory workflows. Parse also offers an offline, HPC-compatible version, which we struggled to run due to software bugs unrelated to user inputs or FASTQ quality. Scale’s ScaleRna pipeline relies on additional installations of Nextflow and docker/singularity. Scale’s pipeline operates directly on BCL files and does not automatically export sample-level FASTQs. Users can download and manually run a component of the Nextflow pipeline to extract sample-level FASTQs, though it is not a documented procedure. Scale’s summary did not include saturation curve outputs. These factors are all considerations that must be weighed when choosing kit or offering a service for both small and large projects.

### Cellular Data Recovery and Cost

We calculated the percent cell recovery using the number of cells loaded and recovered from the analysis pipelines using default settings and consistent filtering thresholds (Figure 2a-b). Low-quality cells and multiplets were removed from further analysis (see “Filtering count matrices” in Methods). Average high-quality single-cell recovery in BM1 ranged from 21.5 ± 2.5%, SE (Parse v3) to 71.1 ± 1.5% SE (GEM-X 3’) (Figure 2a). The updated GEM-X chemistry increased recovery by almost 20%. Fluent V’s cell recovery (43.1 ± 1.0%, SE) improved over the previous version (37.6 ± 2.0%, SE). Cell recovery varied more across sample replicates for fixed samples in Fluent v4, Parse v3, and Scale (SE = 2.0-4.1%) compared to other kits (SE = 0.3%-1.4%) since each replicate was fixed and counted separately. Both plate-based assays delivered a lower percent recovery, as calculated and expected for loading the first plate, which accounts for loss through the barcoding process. Fluent V had the lowest multiplet rate of recovered cells (3.6 ± 0.2%, SE) compared to Scale (8.8% ± 2.9%) and Parse v3 (11.6% ± 0.5%). Both 10x 3’ platforms had higher than expected multiplet rates (10%-11.6%) likely due to chip overloading since we recovered more than 20,000 cells per sample. 10x 3’ kits also observed more low-quality cells (2.3-7.6% of recovered cells) than other kits (0.2-2.5%) due to looser cell-calling criteria in their pipeline. In BM2, the cell recoveries followed a similar trend with GEM-X 5’ outperforming NextGEM 5’ by 15% on average. NextGEM 5’ had almost 2X higher percent recovery and GEM-X had 2.5X high-quality singlets when compared to Parse v2 (Figure 2b).

**Figure 2:**
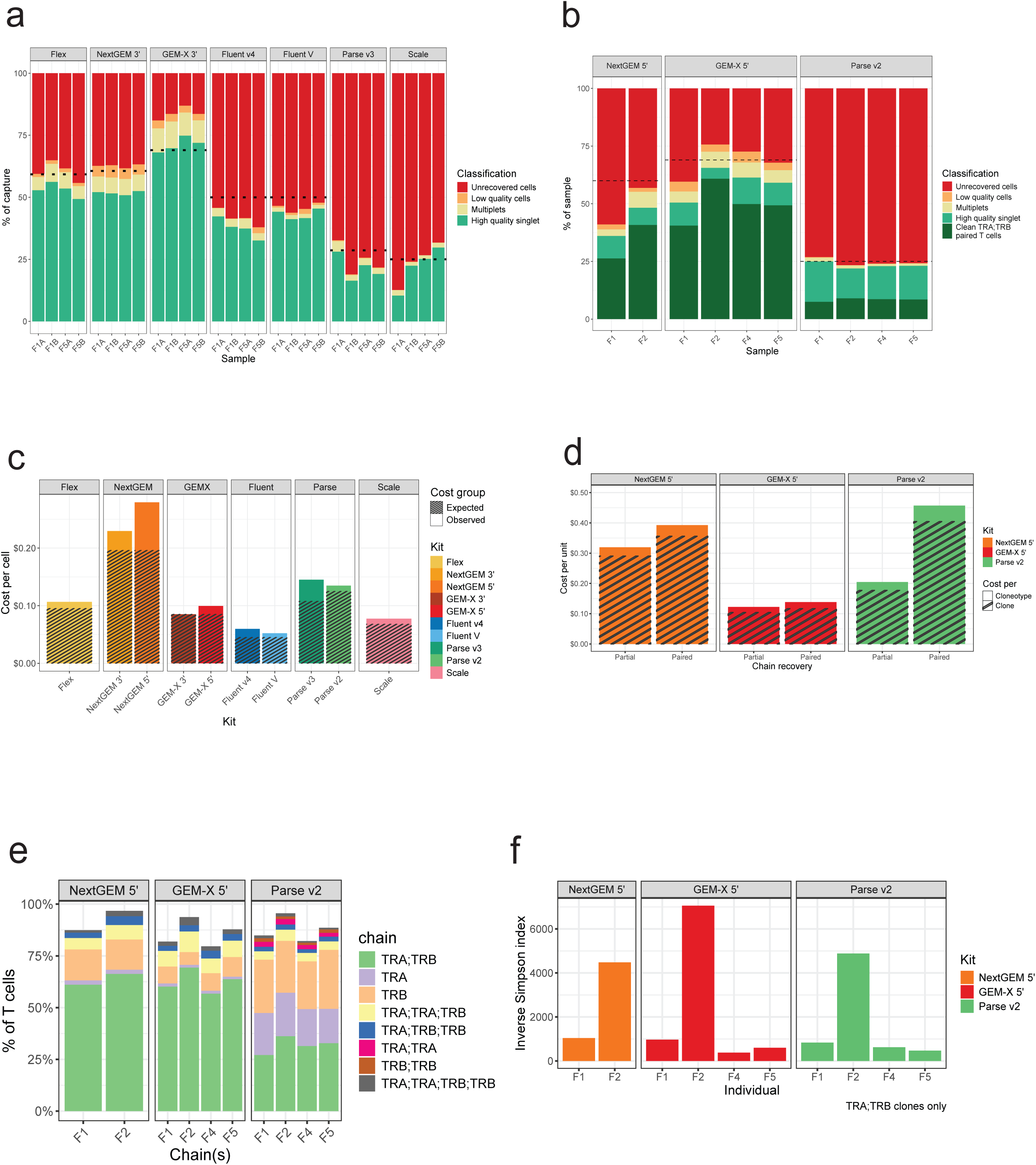
Cellular data recovery a. Cell recovery of Benchmark 1 transcriptome assays from whole PBMCs. Recovery is reported as the percentage of the total cells loaded. Dotted lines indicate the expected cell recovery calculated for each kit from the respective user manuals for each input. Unrecovered cells (red) are the difference between loaded cells and barcodes reported in the filtered matrix. Low-quality cells (orange) and multiplets (yellow) are barcodes filtered out by global quality thresholds and doublet detection during post-processing steps (see computational methods section for details). High-quality singlets (green) are cell barcodes used in downstream analyses. b. Cell recovery of Benchmark 2 transcriptome libraries from sorted T-cells as described in 2a. Dark green indicates T cells containing one paired αβ clone. c. Cost per cell based on kit cost and the observed cell recovery compared to the expected cell recovery projected by kit manuals. d. Recovery of TCR chains as a percent of total T cells per sample. Partial clones are missing either an α or β chain, while paired clones have both. The total population of T cells is calculated by the unique barcodes with reads in the TCR data, regardless of whether productive TCR chains could be assembled for that cell. e. Inverse Simpson index of complete and clean αβ clones per sample (no duplicate or missing chains). A higher value indicates a more diverse and even distribution of clonotypes. f. Cost per clone (productive T cell) and clonotype (unique chains) for VDJ assays. Partial recovery includes all clones with at least one alpha or beta chain. Paired recovery requires both an alpha and a beta chain.

Using our cell recoveries, we calculated the reagent cost per cell compared to the expected cost per cell based on each vendor’s projected recovery (Figure 2c). Fluent V provided the lowest expected cost per cell at $0.045/cell compared to our observed $0.059/cell cost. Scale had the next most cost-efficient recovery ($0.068/cell expected vs. $0.078/cell observed) followed by GEM-X ($0.083/cell) and 10x Flex’s observed $0.107/cell. Due to better cell recovery, Parse TCR kit’s observed cost per cell ($0.135) was less than Parse v3 ($0.145), even though the kit is more expensive. For context, in an experiment targeting 100,000 cells, a price difference of $0.01/cell costs $1,000.

For BM2, which was performed on sorted CD3+ T cells, we also prepared and sequenced TCRα and TCRβ which allowed for recovery of VDJ junctional sequences. TCR clone recovery was cheapest with GEM-X 5’ (Figure 2d). We calculated the frequencies of individual chains, paired chains, or instances with multiple versions of recovered TCRα and TCRβ for each sample (Figure 2e). This showed that NextGEM 5’ and GEM-X 5’ had higher rates of paired TCRα and TCRβ chain recoveries than for Parse v2 TCR. We also calculated the Inverse Simpson index of clonotype diversity. All kits showed good correspondence in that our F2 sample had the highest level of TCR sequence diversity (Figure 2f). Parse v2 was more cost-effective for partial chain recovery than NextGEM 5’, but less effective for paired chain recovery.

### Sequencing Efficiency and Cost

An additional expense of any scRNA experiment includes the cost of sequencing, which is often excluded from platform comparisons. We observed variation in the sequencing efficiency among the platforms which adds to overall experiment costs. For example, Flex performed very efficiently with 88.9% of reads aligned and from viable cells, whereas Fluent v4 and Parse v3 had less than 40% (Supplemental Figure 2a-b). Ambient RNA can also contribute to sequencing inefficiency. All kits had similarly low levels (< 5%) of estimated ambient RNA contamination, except for Fluent v4 with >10% (Supplemental Figure 2c-d). We opted to retain these reads to avoid transformations that could generate bias among kits.

The recommended read depth to achieve sequencing saturation ranged from 10,000-30,000 reads per cell. We downsampled the data to generate saturation curves consistently across kits (see “Sequencing saturation” in Methods, Figure 3a-d). Since the sequencing saturation output metrics varied across pipelines, we calculated the number of reads needed to recover half of the maximal expected value (rd50) as a measure of sequencing efficiency [13]. A lower rd50 suggests the kit recovered its projected yield with less sequencing, while a higher rd50 suggests the kit may have proportionally greater data recovery with deeper sequencing. The rd50 should be considered along with the projected yields (Supplemental Table 2) to calculate the value and utility of deeper sequencing.

**Figure 3:**
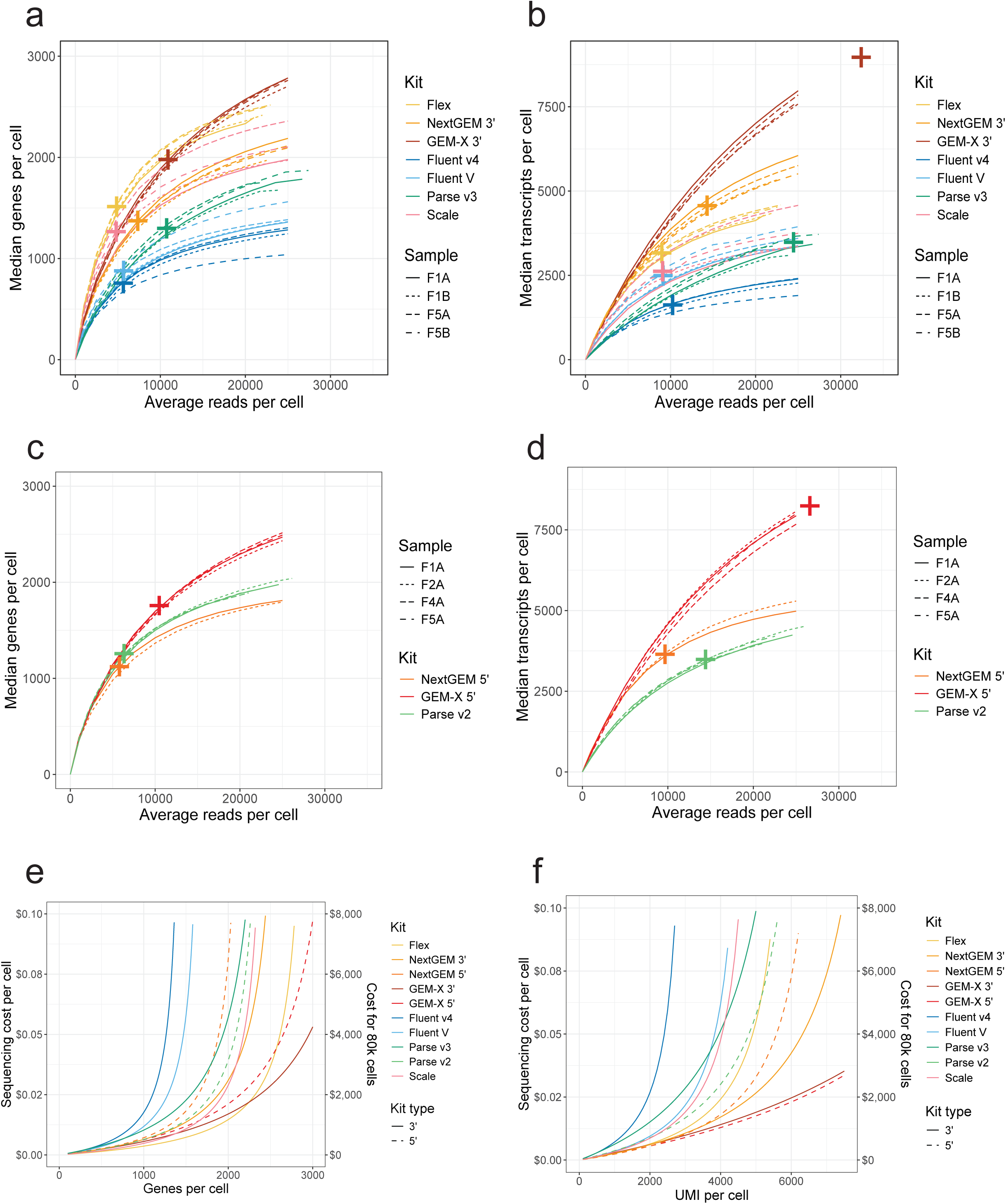
Sequencing saturation and cost Sequencing saturation curves showing recovery of genes (panels a and c) and UMI (b, d) per cell at different read depths for both 3’ (a,b) and 5’ (c, d) assays. Data is downsampled at the FASTQ level to a target number of reads per cell. Crosses indicate the half-maximal saturation value (see methods for rd50 calculation). e-f. Sequencing cost for different targets e) genes per cell and f) UMI per cell. Target read depth is estimated per-kit by calculating sequencing efficiency for read utilization (Figure 2D) and unique gene/UMI recovery (Figure 3A-D). The cost per read is based on a 1 billion read sequencing run for $1,500. The left y-axis shows the sequencing cost per cell based on the target read depth. The right y-axis shows an example sequencing cost for an 80k cell experiment.

In BM1, Flex’s gene recovery exceeded GEM-X at lower sequencing depths but was surpassed after ∼15,000 reads per cell (Figure 3a). GEM-X 3’ surpassed Flex’s UMI recovery regardless of sequencing depth (Figure 3b). GEM-X 3’ and Parse v3 had the highest rd50s for transcripts and genes, suggesting they would benefit the most (proportionally) from deeper sequencing. Flex, Scale, and Fluent all had similarly low rd50 values (rd50 for genes = 4,798-5,678, rd50 for UMI = 8,971-9,647). For Benchmark 2 WT libraries, GEM-X 5’ retrieved the most genes and UMIs at all depths (Figure 3c-d). The rd50s and sensitivity of Parse v2 and NextGEM 5’ were similar, though Parse detected more genes while NextGEM had better UMI recovery.

We modeled the per-cell sequencing requirements for each kit using the rd50 to estimate the cost per cell to reach UMI or gene per-cell targets (Figure 3e-f). The cost is based on a fixed sequencing price per read. We also calculated the cost of an example experiment requiring 2,000 UMI per cell for 80,000 cells (Supplemental Table 2). Sequencing cost was most efficient with Flex and GEM-X (Figure 3e) due to their steep saturation curves (low rd50) and high UMI and gene recovery. For 5’ assays, Parse v2 cost less than NextGEM for prioritizing gene recovery but more expensive for prioritizing UMI recovery (Figure 3f).

### Sensitivity and feature recovery

To analyze each kit’s ability to resolve biological signals, we considered downsampling the number of cells across kits to a fixed value. However, we observed that downsampling cells did not reduce the ability to resolve cell populations. Total cell count can affect cluster accuracy and the ability to resolve cell subsets, but it is not clear whether the effect is positive or negative [15]. We chose to analyze each dataset with all recovered high-quality cells as they would be used in a standard experiment.

In BM1, GEM-X detected more genes and UMIs per cell compared to the other kits, averaging 24% more genes than Scale and NextGEM and 30% more UMIs than Scale and NextGEM. (Figure 4a, Supplemental Figure 1c). Fluent’s kits had the lowest unique feature and transcript recovery of the 3’ assays, though the updated Fluent V chemistry recovered 23% more genes and 64% more UMIs per cell than v4. For 5’ assays, GEM-X 5’ had the highest feature and UMI recovery, while Parse v2 had more genes but fewer UMIs than NextGEM 5’ (Figure 3c-d, Supplemental Figure 1c). Metric distribution ranges were different across kits since each vendor pipeline’s cell calling algorithms had their thresholds for minimum data recovery. Fluent and Scale’s cell calling criteria allowed differing genes and UMIs per cell thresholds between samples. This inter-sample variation must be managed carefully when making biological comparisons with multiple samples.

**Figure 4:**
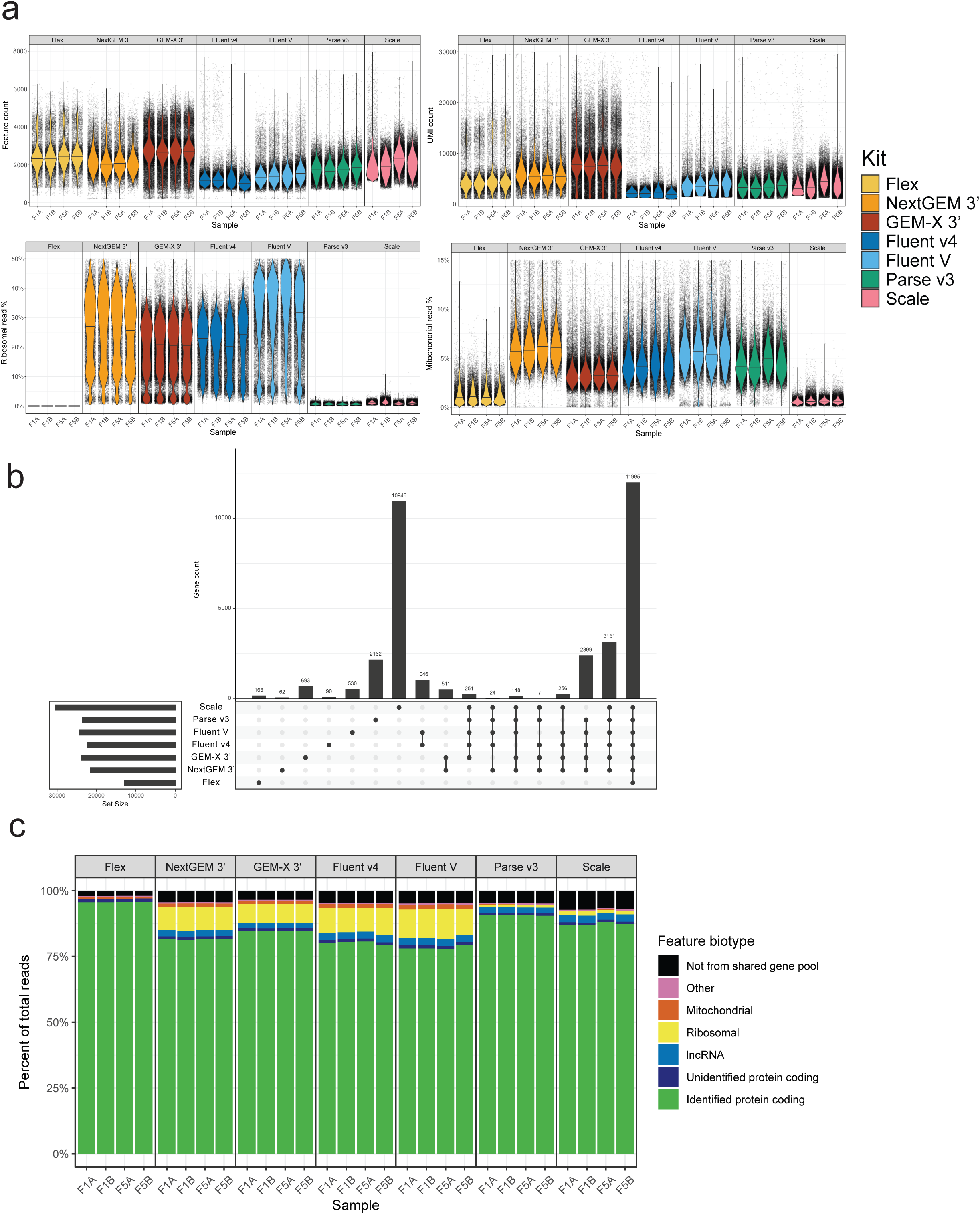
Gene and UMI sensitivity metrics a. Distribution of cell-level quality metrics per sample. Each point is a single cell. The violin overlays show the density distribution of cells in the sample with the median indicated by the line. From the top left going clockwise, the metrics are: feature (gene) count, UMI (transcript) count, percent of transcripts from mitochondrial genes, percent of transcripts from ribosomal genes. b. Genes that are uniquely detected in each kit and shared between kits. Each entry shows genes only found in that intersection. A gene was considered detected at the kit level if it was found in 10 or more cells and in 2 or more samples. Not all intersections and genes are accounted for in this plot; only the main intersections of interest with meaningful values are shown. c. Percent of reads assigned to feature biotype categories from Ensembl. Protein coding genes were binned as “identified” if they had a gene symbol or “unidentified” if they had location information or unique identifiers but no gene symbol. “Other” indicates a feature could not be binned as only one of the other biotypes. “Not shared” indicates the feature was not detected in all other kits. See supplemental figure 1A for a breakdown of the classifications for the non-shared features.

Ribosomal protein (RP) and mitochondrial DNA (MT) gene expression constitute a portion of sequencing reads that are generally used for QC purposes indicative of cell health in comparative studies rather than measuring biological outcomes. Plate-based assays (Parse and Scale) had a lower percentage of RP reads per cell (0.7-1.0%) than other kits, except Flex which has no ribosomal protein probes (Figure 4a). Scale also had the lowest percentage of MT reads (0.6% ± 0%), while Fluent V and NextGEM 3’ were highest (5.6-5.9%). Less MT and RP reads were detected in kits with cell fixation, except Fluent v4, which is consistent with prior studies [16], [17]. When we ran Fluent’s v4 workflow, their fixation protocol was in early development and may not have resulted in fully fixed cells, which could explain similar rates with other emulsion methods. MT read recovery was similar across 5’ kits. Fewer reads mapping to RP and MT genes allow greater data recovery for more informative features (Figure 4B).

The 3’ kits detected 11,995 shared features, and an additional 3,151 features were found by non-probe-based kits (Figure 4b). Over 90% of the reads for each kit were from shared genes (Figure 4c). Reads that originated from outside the shared gene pool primarily came from lncRNAs and other non-protein coding or ambiguous features (Supplemental Figure 3a, Supplemental Table 1).

Scale recovered the most biological features overall, including ∼11,000 features not found in any other kit (Figure 4b). Most of these genes were detected in trace amounts, and did not represent a major portion of the total reads per sample (Figure 4c). Additionally, most of the unique features were unannotated genomic regions or had unknown biological functions (Supplemental Figure 3a). Scale’s processing pipeline differs from the other kits analyzed in that it defaults to allow multimapping reads to contribute to the counts matrix, however, disabling the partial alignment of UMIs in favor of single-gene assignment via expectation maximization only decreased the median genes per cell by 21 (1%) and lowered the total detected genes by 62 (0.2%). Gene recovery patterns in 5’ data were similar, with most features being shared across kits and nearly all reads derived from these shared features (Supplemental Figure 3b-c).

### Cell annotation

Data were merged and clustered at the kit level and annotated using marker genes and reference projections (see “Cell Type Annotation” in Methods, Figure 5a, Supplemental Figure 4). We quantified the ease of cell type calling by summarizing the expression of canonical gene markers as a single module (Figure 5b). Populations with strong expression of a single cell type module and low expression of other modules were more confidently annotated, alternatively, populations with mixed expression of multiple cell type modules were difficult to annotate (Supplemental Figure 5). Flex observed the highest cell type-specific marker module expression, followed by GEM-X 3’ NextGEM 3’ and Fluent V. Parse and Scale exhibited weaker expression of canonical markers. A similar pattern was observed regarding the confusion with other cell type markers: Flex had less bleed-through of other cell type signatures, while Parse and Scale had higher expression of confounding marker genes.

**Figure 5:**
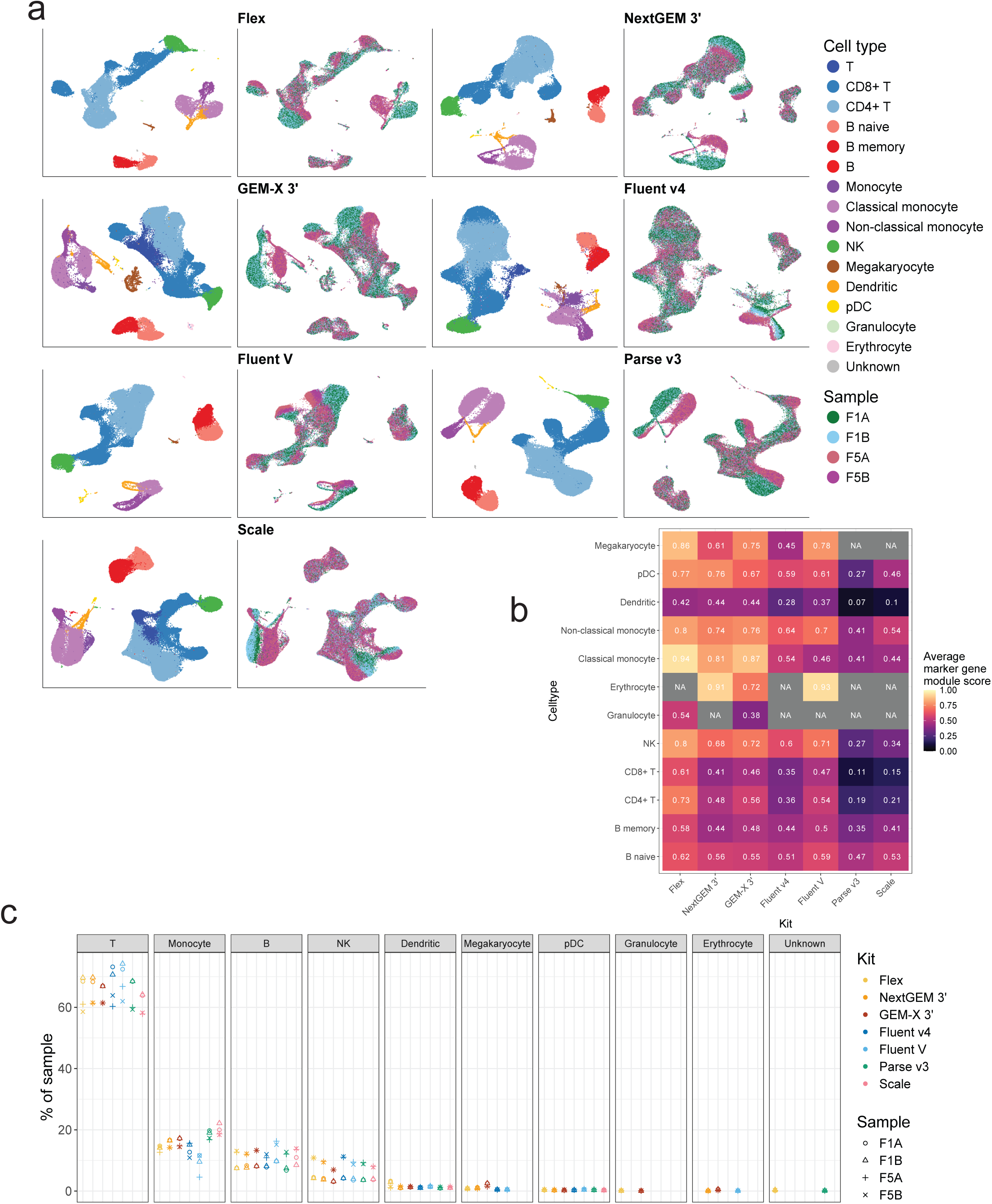
UMAPs of sample replicates and cell composition for each platform. a. UMAP projections of data merged at the kit level. Celltype labels from cluster-level annotation are shown on the left panels. Sample labels are shown on the right panels. b. Within each celltype, the expression of marker genes for that celltype is approximated by a module score, where a higher score indicates those genes are expressed more highly than would be expected by chance. The module score is averaged across cells and shown here as an approximation of how strongly resolved each celltype is in a kit. See supplemental figure 5 for confusion matrices of all celltypes and marker sets. c. Celltype composition of samples. T cells, b cells, and monocytes are grouped by their parent classification for simplicity (e.g., CD4+ and CD8+ T cells are grouped). No significant differences in compositions across kits were observed.

We used cell type annotations to profile the cell compositions from single-cell sequencing data. We aggregated cells in high-level classifications to minimize the effect of annotation ambiguity. The proportional compositions were not significantly different between kits (Figure 5c). Flex and GEM-X 3’ detected trace amounts of granulocytes and NextGEM 3’, GEM-X 3’, and Fluent V detected trace amounts of erythrocytes, but these niche populations represent <1% of the total cells captured.

### Sample replicability

Overall, the similarity of replicate samples from the same individual was quantified at the single-cell level using earth mover’s distance (EMD). Flex demonstrated the lowest EMD and lowest variance between samples, with Fluent v4 and Scale having the highest values and more inter-sample variance (Figure 6a). We considered if Flex’s low values may be due to the more minimal downsampling for this kit, as Flex yielded the fewest cells (mean per sample = 8,935) and thus set the threshold for cell downsampling in EMD calculations. We downsampled the other kits to 10,000 cells and performed the same EMD procedure, but it did not change the EMD values systematically. High intra-sample replicability will improve the ability to detect differentially expressed genes.

**Figure 6:**
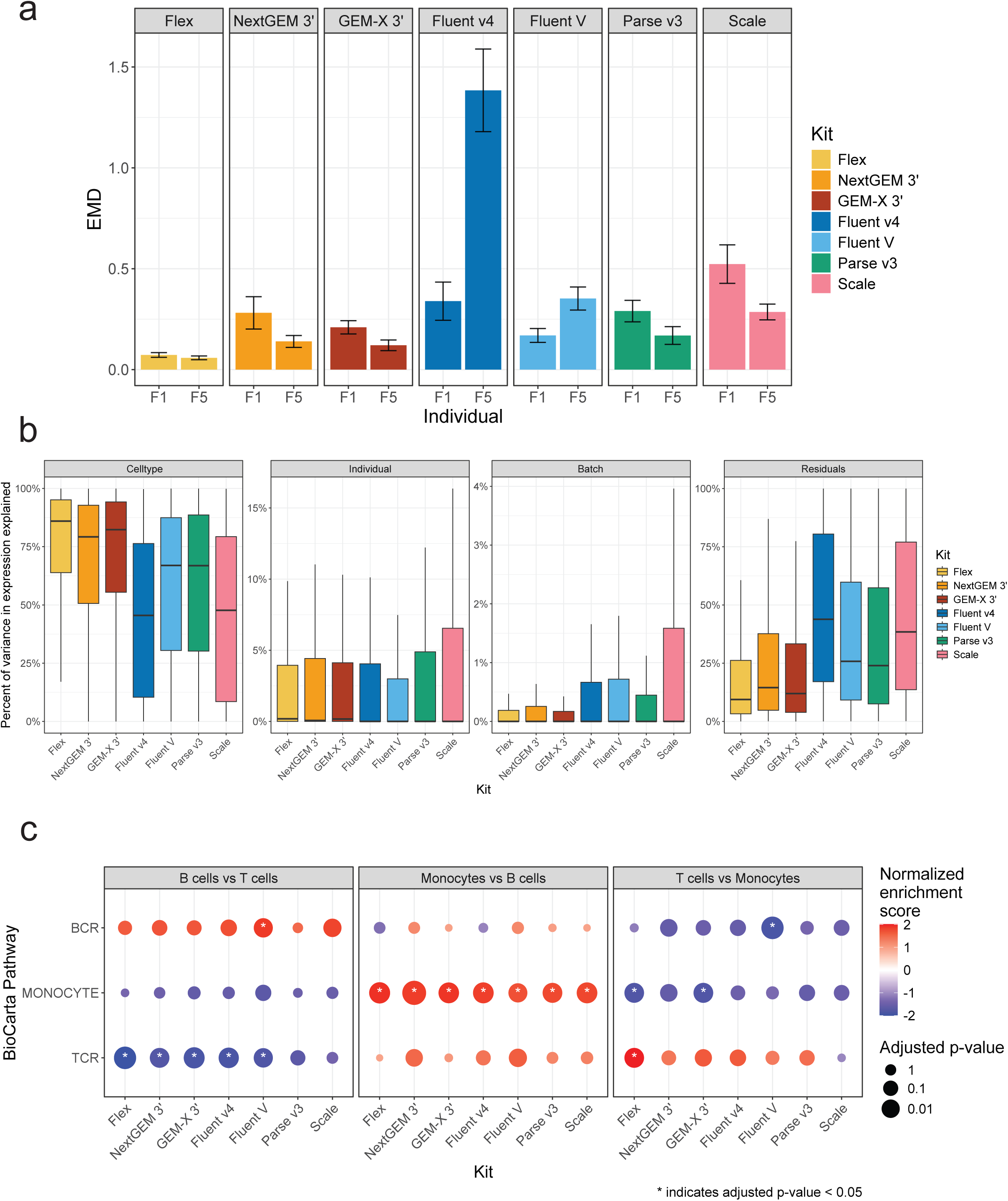
Detecting biological differences a. Earth mover’s distance (EMD) approximates the total difference between replicates, where a higher value indicates more difference. EMD was bootstrapped on data downsampled to 8k cells per sample. EMD provides a quantification of sample replicability including expression and composition as it is blind to discrete cell classifications. b. Percent of total variance in a gene’s expression attributable to explanatory variables. Gene expression was modelled on a per-kit basis with celltype, individual, and batch as fixed effects. The portion of expression variance not modelled by these terms are the residuals. Higher values for celltype and individual suggest good sensitivity in detecting true biological differences. Higher values for batch or residuals indicate more variability in results and less power to detect differential gene expression. c. Gene set enrichment analysis (GSEA) of BioCarta pathways using differential gene expression calculations between pseudobulk aggregated groups of cells within kits. Positive normalized enrichment scores (NES) indicate higher expression of pathway genes in the first group, while negative NES indicates higher expression in the second group. For example, a positive NES in the first sub-panel indicates enrichment of those genes in B cells relative to T cells, while a negative NES indicates enrichment in T cells relative to B cells. Asterisks indicate pathways with an adjusted P value below 0.05.

### Differential gene expression

We aggregated the expression of single cells into pseudo-bulk cell populations for each sample as pseudobulking provides a more precise and accurate calculation of differential gene expression (DGE) (Squair et al., 2021).We observed that the magnitude of downsampling cells for pseudobulking had a strong effect on the overlap in DEGs: when pseudobulk samples were created from more cells, DEGs were more similar across kits. This indicates the importance of high cell recovery for reliable estimates of biological differences. We downsampled to 2,500 cells per cell type for the final analysis to approximate the power in a DGE analysis of a moderately abundant cell population. Each kit found thousands of significantly differentially expressed genes (DEGs) for each contrast. Many of the DEGs were shared between kits, though each kit found several hundred to several thousand significant genes not detected in other kits.

Gene expression was modeled at the kit level as a function of cell type, individual, and batch. Variance in each gene’s expression was broken down by these explanatory terms, with residuals being the remaining unexplained variance (Figure 6b). Flex, GEM-X 3’, and NextGEM 3’ had the highest variance attributable to cell type (86 - 79%) and the lowest residuals (9-15%). This implies a strong ability to capture biological differences in a reproducible manner. Scale and Fluent v4 showed lower variance attributable to cell type (48%, 46% respectively) and higher residuals (38%, 44%), suggesting weaker ability to detect biological differences. Scale also had more variance attributable to batch (upper quartile = 1.6%, other kits = 0.2-0.7%) which may confound analyses in underpowered studies. Fluent V and Parse v3 performed similarly and fell between the other kits.

We performed gene set enrichment analysis (GSEA) on each comparison to test each kit’s ability to resolve biologically relevant signals from noise. We focused on easily annotated cell types (B cells, T cells, and monocytes) and a handful of pathways relating to these cell types to act as positive controls for what we predicted to resolve (Figure 6c). For example, all kits showed the expected significant enrichment of the monocyte pathway when comparing monocytes to B cells. Similarly, all kits showed significant enrichment of the TCR pathway when comparing T and B cells except Parse v3 and Scale. Only GEM-X 3’ and Flex were capable of correctly identifying the enrichment of the monocyte pathway when comparing monocytes with T cells, and Fluent V was the only kit to observe significant enrichment of the BCR pathway when contrasting T and B cells. These findings were consistent with the hypotheses that high sample replicability and low residual variance lead to superior resolution of the biological signal.

## Discussion

We expanded upon recent evaluations of commercially available single-cell RNA sequencing (scRNA-seq) platforms [12], [13] by benchmarking the latest chemistries and analyzing performance across technologies. Our goal was to give researchers and service providers a clear roadmap for selecting the most suitable scRNA-seq solution for their unique needs. We conducted two in-depth comparisons: Benchmark 1 assessed sensitivity and reproducibility using PBMCs from two donors in replicate across seven platforms and Benchmark 2 focused on T cells sorted from four donors, evaluating gene expression and TCR profiling across three kits. We targeted each platform’s maximal recommended cell capture limits to ensure a practical comparison in our appraisal of the user experience and kit performance. A high-level summary of key metrics from our evaluation is provided in Table 2 and Table 3.

**Table 2.** Summary metrics for benchmark 1. Cells are colored based on absolute deviation from the median value for the column. Stronger color values indicate greater deviance. Green indicates better performance than the median, while red indicates worse performance. Text color values have no meaning, they only differ to provide contrast from the background.

|  | Cell recovery |  | Sequencing efficiency |  |  |  | Feature recovery |  |  | Replicability |  |  |
| --- | --- | --- | --- | --- | --- | --- | --- | --- | --- | --- | --- | --- |
|  | High quality cell recovery | Cost per high quality cell | Reads mapped to transcriptome | Reads contributing to counts matrix | Estimated maximum gene recovery | Estimated maximum transcript recovery | Median genes per cell | Median transcripts per cell | Transcripts mapping to informative genes | Earth mover distance | Residual variance in gene expression | Successfully detected enriched pathways |
| Flex | 53% | \$0.11 | 94% | 89% | $3.0 \times 10^3$ | $6.2 \times 10^3$ | $2.4 \times 10^3$ | $4.3 \times 10^3$ | 96% | 0.06–0.07 | 9% | 4/6 |
| NextGEM 3' | 52% | \$0.23 | 72% | 68% | $2.7 \times 10^3$ | $9.0 \times 10^3$ | $2.1 \times 10^3$ | $5.6 \times 10^3$ | 82% | 0.14–0.28 | 15% | 2/6 |
| GEM-X 3' | 71% | \$0.08 | 65% | 60% | $3.9 \times 10^3$ | $17.8 \times 10^3$ | $2.7 \times 10^3$ | $7.6 \times 10^3$ | 85% | 0.12–0.21 | 12% | 3/6 |
| Fluent v4 | 38% | \$0.06 | 77% | 37% | $1.5 \times 10^3$ | $3.1 \times 10^3$ | $1.2 \times 10^3$ | $2.1 \times 10^3$ | 80% | 0.34–1.38 | 44% | 2/6 |
| Fluent V | 43% | \$0.05 | 73% | 58% | $1.7 \times 10^3$ | $4.9 \times 10^3$ | $1.4 \times 10^3$ | $3.6 \times 10^3$ | 78% | 0.17–0.35 | 26% | 3/6 |
| Parse v3 | 22% | \$0.15 | 38% | 34% | $2.6 \times 10^3$ | $6.9 \times 10^3$ | $1.8 \times 10^3$ | $3.4 \times 10^3$ | 91% | 0.17–0.29 | 24% | 1/6 |
| Scale | 22% | \$0.08 | 63% | 57% | $2.5 \times 10^3$ | $5.1 \times 10^3$ | $2.0 \times 10^3$ | $3.6 \times 10^3$ | 87% | 0.29–0.52 | 39% | 1/6 |

**Table 3.** Summary metrics for benchmark 2. Cells are colored based on absolute deviation from the median value for the column. Stronger color values indicate greater deviance. Green indicates better performance than the median, while red indicates worse performance. Text color values have no meaning, they only differ to provide contrast from the background.

|  | Cell recovery |  | Sequencing efficiency |  |  |  | Feature recovery |  |  | TCR recovery |  |
| --- | --- | --- | --- | --- | --- | --- | --- | --- | --- | --- | --- |
|  | High quality cell recovery | Cost per high quality cell | Reads mapped to transcriptome | Reads contributing to counts matrix | Estimated maximum gene recovery | Estimated maximum transcript recovery | Median genes per cell | Median transcripts per cell | Transcripts mapping to informative genes | Cost per recovered clone pair | Cells with paired clones |
| NextGEM 5' | 42% | \$0.28 | 71% | 69% | $2.2 \times 10^3$ | $7.2 \times 10^3$ | $1.8 \times 10^3$ | $5.1 \times 10^3$ | 81% | \$0.36 | 34% |
| GEM-X 5' | 59% | \$0.10 | 63% | 60% | $3.5 \times 10^3$ | $16.4 \times 10^3$ | $2.5 \times 10^3$ | $7.9 \times 10^3$ | 83% | \$0.12 | 50% |
| Parse v2 | 23% | \$0.13 | 54% | 49% | $2.5 \times 10^3$ | $6.9 \times 10^3$ | $1.9 \times 10^3$ | $4.2 \times 10^3$ | 90% | \$0.40 | 8% |

Our assessment of sample flexibility, total runtime, built-in pause points, and overall throughput overlapped with the experimental features discussed by [13] and suggested that longer, labor-intensive assays performed by a service lab may offset a low kit cost for the user. Alternatively, required equipment costs, such as an upgrade to the Chromium X, required for Flex and GEM-X, or optional automation for combinatorial plate-based assays may be a barrier to start-up, however, the investment saves time, increases throughput, and reproducibility which is beneficial to service laboratories or high through-put labs. Fluent’s PIPseeker (now Illumina Single Cell 3’ RNA Prep) offered the best cost-per-cell and easy analysis, ideally for individual users on a tight budget who can afford or share the low equipment cost. Combinatorial indexing plate assays offer sample flexibility and throughput due to the ability to fix and freeze samples, aiding experiment planning and sample processing. Parse offered the only option for TCR profiling of fixed cells which is ideal for sorting and storing samples for later processing. We found, however, a greater risk with the “all-in” approach of these plate-based indexing assays, since all samples are processed on a single plate and any error affects the whole experiment. The new 10x GEM-X workflows increased cell throughput without adding time or effort to the protocol allowing users to process more cells than the NextGEM platform. The Chromium X run time is reduced and increases chip throughput; alternatively, individual samples can still be run when optimizing sample preparations. The microfluidics of the new GEM-X chips claim to be more robust to clogs and wetting failures, but if this does occur, it usually affects a portion of the samples. Sample fixation was not supported for GEM-X at the time of this experiment but since we performed our experiments, DSP methanol fixation, like Fluent and Scale’s approach, is supported adding flexibility to store samples for both the GEM-X 3’ and 5’ assays. 10x Flex fixation was the easiest workflow for quickly storing cells (or tissue) for either an hour or overnight, so long as all samples in a single experiment were processed in the same way. While Flex has the longest overall workflow time, it has the least hands-on time and many options to stop and store samples, increasing flexibility and throughput while decreasing risk.

All analysis pipelines ran on a high-performance computing environment with 16+ CPUs and 200+ Gb RAM. Parse and Scale pipelines were more challenging to execute due to installation requirements, a lack of intermediate files, and limited documentation. Parse and 10x offer cloud-based processing solutions that may appeal to researchers with limited computational resources. Pipelines have varying levels of default permissiveness for cell calling: 10x CellRanger defaults to returning more low-yield cells, while Fluent has more conservative default thresholds. Researchers must understand pipeline default behavior and weigh their priorities (data quality vs quantity) to adjust pipeline parameters to meet their study goals.

10x GEM-X assays led in cell recovery efficiency and had a higher proportion of low-quality cells and multiplets. Studies have shown droplet-based methods to have a positive correlation between the total cells loaded and the multiplet rate [19], [20] while split-pool combinatorial methods have an increased risk of doublets due to cell clumping [21]. Both overloaded 10x 3’ chips likely resulted in higher doublet rates due to overloading. However, the sensitivity of the doublet detection algorithm employed for this study is related to the strength of cell type-specific expression profiles; a moderately higher doublet rate can be a positive indicator of filtered data quality, not just a higher doublet incidence [22].

Recovery is even more important for cell-limited samples. The ability to include cell-limited samples in assays that require fixation is important due to the additional cell loss at this step. For example, 10x Flex, Parse, and Scale each provide guidance on lower limits for cell fixation and pooling. Combinatorial indexing methods required starting with over 350,000 total cells across samples due to the calculated loss of cells through plastics and pipetting transfers. Combining multiple low-input sample types, like cells and nuclei to utilize all the wells, is not recommended due to differences in cell quality and RNA expression levels which can create sample bias [21]. The utility of using fixed cells for TCR profiling with Parse was much anticipated to alleviate the need to process patient samples immediately and to limit bulk effects. While Parse TCR and 10x 5’ platforms recovered similar frequencies of unpaired α and β chains, the Parse kit recovered only half the number of cells with paired αβ chains. Since DSP fixation is now possible with the 10x platforms, users may choose their platform based on access to an instrument, sample input limitations, or cost.

Flex was highly cost-efficient and delivered the best value for shallow read depth due to its high read utilization and efficient gene recovery. GEM-X 3’ and 5’ produced high yields with moderate sequencing and efficiently generated more usable data with additional read depth. Fluent did not recover as many genes and UMIs but approached sequencing saturation quickly. Despite less overall data, Fluent V was still able to adequately resolve cell populations and expected biological activity. We observed slightly lower median gene and UMI recovery for Parse v3 compared to our Parse v2 data and the vendor-published web summary found on their website [23] and our Parse v2 TCR data. The lower recovery may be a product of the additional time the cells sat on ice while counting with a hemocytometer for accurate recovery metrics before adding them to the plate. Our sensitivity metrics comparing Parse v2 and 10x NextGEM were consistent with an analysis of thymocytes [14] and the opposite of a study using replicate PBMCs [12].

While Scale achieved similar median cell feature recovery to other kits, it identified over 10,000 features not detected by any other kit. Nonetheless, most of these features were either unannotated or non-protein-coding and constituted only a small fraction of the total recovered reads. These additional genes did not provide valuable biological insights and will weaken the power of differential expression testing when included in multiple hypothesis-testing correction methods. The default pipeline settings in Scale allow for reads to be assigned to multiple genes, but this accounts for just around 1% of the unique features found. In contrast, Parse v3 identified more than 2,000 unique features, likely due to its combination of oligo dT and random hexamer primers.

Our sequencing cost calculations did not factor in any variability in the sequencing run, e.g. the cluster percentage or the number of reads passing filter off the sequencers. We chose not to include these metrics since we did not have values for all libraries since the 10x libraries were pooled together, and some libraries were run on different models of Illumina instruments. Scale was the only library sequenced on two different sequencers and consistently produced the lowest observed reads passing filter percentage (55-66%) and less than the expected reads for the kit. Consequently, we had to sequence more to get the required number of reads which increased the cost per cell. The read configuration of the Parse v2 TCR kit still requires a 200-cycle sequencing cartridge for the WT library, making it more expensive to sequence, whereas all others can be processed with 100-cycle kits.

Accurately annotating cell populations is essential for downstream applications, and poorer resolution results in less power to detect biological differences. Although each technology was able to resolve cells into broad cell type classifications, distinguishing more niche subsets such as CD4+ and CD8+ T cells was more difficult with combinatorial plate-based assays. Consistent with other studies, Scale and Parse exhibited decreased expression or increased dropout of canonical marker genes, making them less capable of resolving some niche cell subsets [13], [24]. Intra-replicate variability is another important factor for statistical power: the more consistent replicates are, the more confidence there is in an observed difference. Scale and Fluent v4 showed the highest intra-sample variability, resulting in more residual variance in differential expression comparisons, decreasing their ability to resolve expected biological differences via pathway enrichment analysis. Flex had very low intra-sample variability and residual variance, resulting in the best performance at resolving expected biological differences. High-fidelity probe alignment produced consistent expression profiles that lend ample power to statistical comparisons [13].

Current technologies continue to evolve, and we expect new platforms to emerge. For example, multiple vendors now offer kits to process up to 5 million cells, proposing lower costs per cell. For larger projects, sample sequencing efficiency will be an important consideration. We focused this comparison on mRNA and TCR sequencing but each of these vendors are offering or developing a broad range of multi-modal applications that should be considered as options and for future comparative studies. No single scRNA-seq platform checks every box. The right choice depends on your experimental priorities be it sensitivity, sample flexibility, cost, or ease of use. We’ve outlined the key trade-offs to help researchers and service providers make informed decisions about commercially available platforms.

## Limitations of Study

Scale and Fluent V data were generated using separate aliquots of the same donor cells at later dates than the initial Benchmark 1 experiment. Any difference in cell viability due to processing separate aliquots could influence metrics.

We overloaded the 10x 3’ chips due to an automated counting error. The actual counts were recovered from hemocytometer images. It is possible that slightly more than 1 million cells were fixed but trypan blue imaging did not reveal any live cells.

We loaded cells into the first plate Parse v3 plate 15-20 minutes past the recommended time from thawing. The new protocol suggests counting an aliquot at the time of fixation before freezing to reduce the time counting cells to 30 minutes before loading the plate.

The NextGEM 5’ data was only presented for two samples because, F4 and F5, were removed from further analysis and comparison. We believe there was a processing error during the recovery step where RNA was lost. We believe this was a manual error and not a sample issue as the sample was run through the GEM-X and Parse TCR workflows and those data were not affected.

## Data availability

Processing and analysis code is available at https://github.com/Fred-Hutch-Innovation-Lab/BM02_analysis. Raw values for all figures are included in Supplementary Table 1. Raw sequencing data, processed counts matrices, and final annotated Seurat objects are available through GEO accession number GSE295527.

## Acknowledgments

This work was supported by the Immunotherapy Integrated Research Center of the Fred Hutchinson Cancer Center and by grants from the National Institute of Health (NIH, P30CA015704, P01CA225517, P01CA018029, U19AI128914, and R01CA264646). This work was partially supported in-kind by 10x Genomics and Scale Biosciences. We wish to thank all the vendors for their technical support, the Fred Hutch Flow Cytometry Shared Resources for their services, and Emily Park for her help with sample preparation. Workflow schematic figures were created with Biorender.

## Author Contributions

A.E.E. conceived the study, designed the experiments, interpreted the results, and wrote the manuscript. D.G. analyzed the data, interpreted the results, and wrote the manuscript. A.E.E., A.L., and D.S. performed the experiments. A.H. analyzed the data. E.W.N. interpreted the results and wrote the manuscript.

## Supplemental Materials

Supplementary Figure 1: Benchmark 2 experimental workflow, timing, and QC

a. Workflow diagram of different steps required for each platform used in the Benchmark 2 experiment from the same cell suspension from sorting T cells to library sequencing. Two samples were processed at a time on two separate days for each platform.

b. Summary of time taken to execute each kit from sample preparation to library completion in terms of hours spent hands-on (active) and time spent waiting (passive). Stopping points are indicated with redlines and the number of stopping points is summarized in the rightmost column. Gray indicates an overnight stopping point.

c. Cell-level QC metric distributions for 5’ assays. Each point is a single cell. The violin overlays show the density distribution of cells in the sample with the median indicated by the line. From the top left going clockwise, the metrics are: feature (gene) count, UMI (transcript) count, % of transcripts from mitochondrial genes, % of transcripts from ribosomal genes.

Supplementary Figure 2: Ambient RNA and read utilization

a-b. Percentage of demultiplexed reads from Benchmark 1 (a) and Benchmark 2 (b) WT gene expression libraries that contributed to the final counts matrix for each pipeline. Reads had to align to reference genome features and be from valid cells to contribute to the counts matrix. Aligned reads not in called cells could be included in the counts matrix with different cell calling criteria.

c-d. Estimated percentage of reads from ambient (i.e. extra-cellular) RNA. Error bars indicate 95% confidence interval for ambient contamination estimates. Ambient RNA was estimated but not removed from the data.

Supplementary Figure 3: Gene recovery

a. Read mapping feature classifications for the non-shared features from 3’ kits. These features were not identified across all kits but may have been found in more than one kit. Identified protein-coding features have gene symbols. Unidentified features have location information or unique identifiers, but no gene symbol. “Other” indicates a feature could not be binned as only one of the other biotypes.

b. Read mapping feature classification for all 5’ data. See classification explanations in (a).

c. Read mapping feature classifications for the non-shared features from 5’ kits. See classification explanations in (a).

Supplementary Figure 4: Celltype annotation

Reference mapping data and marker genes used to annotate clusters of cells. Each kit has one set of figures. The first table shows the top annotation call for each reference annotation used. “Cluster” indicates the unsupervised cluster being annotated. The score column indicates the the annotation method’s confidence in the annotated label (where closer to 1 is better), averaged over all cells with that label. The “%” column indicates the percentage of cells in that cluster with that label. The dotplot below the table shows the normalized expression of canonical marker genes (x axis) in each cluster (y axis). Bigger dots indicate the gene was detected in more cells, while more saturated color values indicate higher expression. The UMAPs below the dotplot show the 30 unsupervised clusters (left) and their labels (right). The table in the bottom right shows the final count and proportion of each celltype.

Supplementary Figure 5: Celltype marker module confusion matrices

The expression of celltype marker genes is approximated by a module score, where a higher score indicates those genes are expressed more highly than would be expected by chance. The module score is averaged across cells with the same label within a kit. For example, in the Flex data, the B naive cells (first column) had high expression of “B” and “B naïve” markers, with relatively lower expression of “B memory” and “Monocyte” markers, and little or no expression of “NK” and “megakaryocyte” markers.

Supplementary Table 1: Extended metrics

Results for all analyses on each sample with kit-level aggregations and error calculations where appropriate.

Supplementary Table 2: Sequencing cost modeling

Modeling the cost of an example experiment intending to recover the same quantity of transcripts per cell.

Supplementary Table 3: Marker genes

Canonical marker genes used to annotate cells and create celltype module scores.

## Methods

### Benchmark 1: 3’ Whole transcriptome comparison

Multiple donor leukopak frozen aliquots were purchased from Bloodworks Northwest. PBMCs from two healthy donors F1 and F5, stored in liquid nitrogen were thawed at 37 degrees C., transferred into 10ml prewarmed thawing media (Gibco RPMI 1640 + 10% FBS) and centrifuged at 500 x g for 5 minutes at 4 degrees. Cells were washed in 10ml cold 1XPBS -MgCl and -CaCl (Gibco), followed by subsequent centrifugation at 500 x g for 5 minutes and the pellet resuspended in 10ml 1X PBS. The CellacaMX (Revvity) live-dead assay was used to count AOPI stained cells. Viability was 83% and 88% respectively. Each cell suspension was aliquoted for use in the 10x Genomics GEM-X and NextGEM assays in which the cell suspensions were loaded in replicate, with a targeted recovery of 20,000 and 10,000 cells, respectively. The remaining cell suspensions were aliquoted out in technical replicates of 1 million cells and fixed using Parse’s Cell Fixation v3 User Manual (UM0027 v1.1), Fluent’s Demonstrated Protocol DSP-Methanol Fixation for Cells (FB0004708), and 10x Demonstrated Protocol Fixation of Cells and Nuclei for Chromium Fixed RNA Profiling (Flex) (CG000478 Rev C).

Separate aliquots of F1 and F5 donors were processed as above except with PBS 1% BSA for the final resuspension. One set was fixed in technical replicates of 1 million cells following the Scale’s standard fixation kit protocol Rev. B. The second set of aliquots was processed as above with PBS + 0.04% BSA and Promega RNase Inhibitor resuspension with no fixation for Fluent Biosystem/Illumina PIPseq V (see below).

#### 10x Genomics Flex Fixed RNA profiling

Per the 10x Genomics CG000527 Rev E protocol, fixed cells were incubated overnight with mRNA transcript probes targeting ∼18K genes, with the left handle containing a sample specific barcode. Following probe hybridization, samples were counted and pooled equally before a series of washes to remove unbound probes. After counting the pool, a volume targeting recovery of 40,000 cells was combined with a master mix that contains enzymes that ligate and extend the probes during GEM incubation. This mixture was then loaded into the NEXT GEM Chip Q with gel beads and partitioning oil. Following microfluidic GEM generation on the ChromiumX, the GEMs were recovered and cDNA was amplified for library preparation. A portion of the amplified material was re-amplified with library-specific P5 and P7 indices.

#### 10x Genomics GEM-X 3’v4 and NextGEM v3.1

Following the 10x Genomics GEM-X 3’v4 (CG000731) and NextGEM v3.1 (CG000315) protocols, PBMCs from donor F1 and donor F5 were combined with a reverse transcript mastermix and loaded onto a microfluidic chip (Chromium GEM-X 3’ chip and NextGEM Chip G) along with partitioning oil and barcoded gel beads. Cells were partitioned into droplets containing gel beads using the ChromiumX instrument and subsequently underwent reverse transcription. The cDNA was recovered, cleaned, and amplified to create enough material for library preparation. To generate gene expression libraries, a portion of cDNA was fragmented, adapter ligated, and an additional index with P5 and P7 handles were added during a final amplification.

#### Fluent Biosciences PIPseq v4

Fixed cells were partitioned, and libraries generated using the PIPseq T20 3’ Single Cell RNA Kit v4.0PLUS User Guide (FB0002130 Rev 8.3). To target a recovery of 20,000 cells per sample, 8uls of a cell suspension at a concentration of approximately 5000 cells/ul were captured in emulsion with Particle-templated Instant Partitions (PIPs), followed by cell lysis and barcoding unique to each PIP. The emulsions were then broken, and the mRNA reverse transcribed to generate cDNA. cDNA was then amplified, isolated from the PIPs, fragmented, end repaired and A-tailed, adapter-ligated, and indexed with unique P7 and P5 indices.

#### Fluent Biosciences PIPseq V

Sample replicates of live cells (40,000 each) were loaded into PIPs for capture and barcoding as described in PIPseq V T20 3’ Single Cell RNA kit User Guide (FB0005261). To target a recovery of 20,000 cells per sample, 8ul of 5000 cells/ul were captured in emulsion with Particle-templated Instant Partitions (PIPs), followed by cell lysis and barcoding unique to each PIP. The emulsions were then broken, and the mRNA reverse transcribed to generate cDNA. cDNA was then amplified, isolated from the PIPs, fragmented, end repaired and A-tailed, adapter ligated, and indexed with unique P7 and P5 indices.

#### Parse Evercode v3

Following the Parse protocol UMWT3300, fixed and permeabilized PBMCs were thawed, and counted using trypan blue and a hemocytometer, diluted and added to 12 wells per replicate of Plate 1 as directed by the sample loading table (48 wells total across all replicates). Following reverse transcription, cells were pooled and split an additional two times, the last of which added a barcode containing R2 and Biotin for magnetic bead enrichment Cells were pooled again, counted, and distributed into eight sublibraries each targeting under 12,500 cells where they were lysed and the biotin-labeled cDNA recovered with binder beads. cDNA bound to the binder beads was washed and incubated with a template switch oligo to add a 5’ adapter to the cDNA. The cDNA was amplified and bead-cleaned. A portion of this cleaned product was fragmented, adapter ligated, and p5 and p7 indices were added in a final PCR.

#### Scale Biosciences

Following the Scale 1020796 Rev A protocol, fixed cells at a concentration of approximately 2000 cells/ul were added to 96 wells of barcode plate 1 and well IDs documented to map back cells to their sample of origin. The cells underwent reverse transcription and were then pooled by centrifugation into a funnel. Cells were washed and then split across a 384 well ligation plate where cells were barcoded. Cells were again pooled by centrifugation, washed, and counted to assess for recovery. The cell stock was diluted to approximately 460 cells/ul and 4uls loaded into each well of the final distribution plate, with the remainder saved at -80 degrees. In the final distribution plate, the cell membrane is broken down, cDNA was tagmented with a transposase, and the library specific P5 and P7 indexes were added.

### Benchmark 2: 5’ WT, TCR single-cell comparison

Frozen PBMCs from four healthy donors (F1, F2, F4, and F5) of mixed sex (3 males, 1 female) and ethnicity (3 persons of Caucasian descent and 1 person of Asian descent) supplied by Bloodworks Northwest were processed in two batches one week apart (F1, F2 followed by F4, F5). Cells were thawed in a 37°C water bath before transfer to thawing media (Gibco RPMI 1640 + 10% FBS). Samples were then centrifuged at 400 x g for 10 minutes at 4°C. The supernatant was decanted, and the pellet resuspended in 10 mL of sorting media (cold Gibco 1X PBS + 1% BSA), followed by an additional centrifugation at 400 x g for 10 minutes.

Each sample was next resuspended in 200 µL of stain media. Cells were stained to enrich T-cells and to remove as many B-cells as possible. We stained the samples with human CD3, CD45, and CD19 (citation for antibodies). 7AAD was used to assess the in-flight viability of the cells during the sort. We sorted T cells using the Becton-Dickenson Symphony S6 cytometer. The final population was CD3+, CD45+, and CD19-.

An aliquot of the sorted T-cells was stained with AOPI and assessed for viability and cell counts using the Nexcelom CellacaMX (*Revvity*). Viabilities ranged from 88-90 % (F1 and F2) and 83-87% (F4 and F5). An aliquot of the of each donor T-cells was utilized for 10x Genomics GEM-X 5ʹ v3 (CG000733 RevA) and NextGEM 5ʹ v2 (CG000331 RevE) assays, in which 20,000 and 10,000 cells were targeted for recovery, respectively. The remaining cell suspension, ∼1.5 million cells, was fixed following Parse Evercode Cell Fixation v2 (UM0039 v2.3). Cells were frozen and later processed as directed by Evercode TCR Human WT + TCR v1 (UM0018).

#### 10x Genomics GEM-X 5’v3 and Next GEM 5’v2

Following the 10x Genomics GEM-X 5’v3 (CG000733) and NextGEM 5’v2 (CG000331) protocols, sorted T cells were added to a reverse transcription mastermix and loaded on to a microfluidic chip (Chromium GEM-X 5’ chip and NextGEM Chip K respectively) with partitioning oil and barcoded gel beads. Cells were partitioned into droplets containing gel beads and subsequently underwent reverse transcription. The cDNA was recovered and bead cleaned before being amplified to create enough material for library preparation for gene expression and TCR. To generate TCR libraries, 2ul of cDNA underwent 2 rounds of PCR with a set of TCR specific primers. A portion of the PCR product was fragmented, adapter ligated, and an additional index with P5 and P7 handles added during a final PCR. To generate gene expression libraries, 10ul of cDNA was fragmented, adapter ligated, and an additional index with P5 and P7 handles added during a final PCR.

#### Parse Evercode TCR

Fixed and permeabilized T cells were thawed, counted using trypan blue and a hemocytometer, following Parse protocol UMIT2300. Each sample was diluted according to the sample loading table and recounted before dispensing into 12 wells per sample of Plate 1 (48 wells total). Following reverse transcription, cells were pooled and split an additional two times to add barcodes, the last pool and split also adding the R2 and biotin. Cells were pooled, counted, and distributed into eight sub-libraries each targeting 12,500 cells. Cells were lysed and biotin labelled, and the cDNA recovered with binder beads. cDNA bound to the binder beads was washed and incubated with a template switch oligo to add a 5’ adapter to the cDNA. The cDNA was amplified and bead cleaned to be usable in TCR and whole transcriptome library preparation. Whole transcription libraries underwent library preparation as described in Evercode TCR Human WT + TCR v1 (UM0018), which utilized the Parse v2 WT chemistry. The TCR library was generated using 10ng of the WT cDNA in a PCR reaction targeting the V(D)J segments of the CDR3 region of the T cell.

#### Processing Comparison Metrics

The workflow processing experience was assessed based on factors that are often considered when executing an experiment. The experience was graded low to high based on how the criteria outlined were met by following each vendor’s protocol requirements and any limitations inherent in the kit.

Sample flexibility was defined as the ability and ease of fixing and storing samples, including the maximum storage time. Kits that have only recently added fixation are rated lowest. Pipetting simplicity was evaluated based on how many pipetting steps are required throughout the workflow, the overall amount of pipette plunges within each of those steps, and the time spent pipetting. We rated extensive pipetting lower than workflows with minimal pipetting. We defined throughput by considering the number of samples and/or the number of cells a kit could process. Kits capable of processing a higher number of cells and samples were rated favorably. We also considered how the workflow timing or handling impacted the potential for increased throughput. Workflow risk will vary based on the user, but we chose to define this category based on the inherent risk of errors within each platform that could impact either individual samples or the complete experiment. Platforms that allowed portions of the kit to be used to process individual samples were considered lower risk.

Data processing was evaluated on a per-vendor basis, as each vendor supplied a processing pipeline that works for the kits offered. We executed each pipeline in a high-performance computing environment, so hardware requirements were not an issue and were not considered for this evaluation. Data processing was evaluated on the following criteria:

- Ease of installation and installation requirements. Standalone pipelines were preferred over pipelines with additional software dependencies.
- Production of intermediate files. BAMs, sample-level FASTQs, molecule info files, and other intermediate outputs facilitate accessory workflows and data introspection. More transparency into the processing is better.
- Clarity and presence of key output metrics: particularly sequencing saturation curves and read utilization statistics. Understanding the sequencing efficiency and read utilization allows the user to quickly assess data quality and decide if additional sequencing would be useful.
- Documentation and ease of troubleshooting. Explanation of error messages and pipeline options should be readily available.

#### QC and Sequencing

All libraries except Scale’s were quantified with the Qubit fluorometer’s High sensitivity dsDNA reagents (Thermo Fisher Scientific) and sized on the Agilent 2200 Tapestation with high sensitivity tapes and reagents. Scale libraries required KAPA qPCR (Roche) quantification to achieve recommended loading concentration since Qubit measurements are not accurate for their libraries. All libraries were run on the Illumina NextSeq2000 except for Scale which was run on both the NextSeq2000 and the NovaSeq6000, and 10x libraries which were proportionally pooled in two separate runs (1. Flex, NextGEM3’ GEM-X 3’; 2. NextGEM5’ and GEM-X5’) and run on a NovaSeq 6000 .

## Computational Methods

### Sequencing Depth and Downsampling

Sample sequencing was conducted to achieve a depth exceeding 25,000 reads per cell for WT libraries except for Flex which targeted 20,000 reads per cell. TCR libraries were all sequenced to a targeted read depth of 5,000 reads per cell. When sequencing surpassed the targets, FASTQs were downsampled to 25,000 reads per cell for the WT libraries and 5,000 reads per cells for TCR libraries using Seqtk version 1.3 with random seed 100 [25].

### Generating count matrices

The software provided by each vendor was utilized to generate UMI count matrices. Pipelines were run with default parameters unless otherwise specified. For the 10x workflows, CellRanger version 8.0.0 with the multi subcommand was employed with the reference version refdata-gex-GRCh38-2024-A. Parse data was processed with splitpipe v1.2.1 using parse’s hg38_V1.1.3 reference. Fluent data was processed with PIPseeker version 3.3.0 with reference gex-reference-GRCh38-2022.04. For each sample, PIPseeker generates 5 outputs of varying sensitivity for calling cells from the background. We selected the sensitivity levels on a per-sample basis informed by the knee plots and vendor guidelines. Scale data was processed through scaleRNA version 1.5 with default parameters, library structure version 1.1 and the prebuilt GRCH38 reference genome provided by Scale. FASTQs were generated at the library level by passing raw BCL folders into the first scaleRNA run. We then ran Scale’s bc_parser (distributed with the Nextflow pipeline) with the “--demux” flag and options to output 2 read files and 2 index files for each sample. These were downsampled and ran with default parameters to produce the filtered expression matrix. To create an unfiltered expression matrix for ambient RNA inspection, downsampled data was processed with an additional scaleRNA run with “minCellRatio” set to 1000.

### Read utilization

Reads were divided into three categories:

1. reads not aligned to transcriptome or missing valid cell barcodes
2. reads aligned to transcriptome and with a cell barcode but not from a cell passing the pipeline’s filter
3. reads aligned to transcriptome and attributed to valid cells

The first category represents unusable data. The second category represents data that could be used if the cell calling criteria were adjusted. The final category represents the data contributing to the filtered reads matrix.

For all datasets, the first category was calculated as the difference between the total fastq reads and the reads in the second and third category. Calculation of the second and third categories differed by vendor.

For 10x, the second and third categories were extracted from the “sample_molecule_info.h5” datasets “feature_read_pairs” and “usable_read_pairs” respectively. Flex has samples mixed in the off-sequencer FASTQs, which must be further demultiplexed as part of the processing pipeline, but this can cause some minor read loss that is not reported in the pipeline reports.

The total original reads for each Flex sample were approximated by multiplying the “UMI per probe barcode” percentage by the total reads for the combined run.

For Fluent, the second category was reported as “num_mapped_reads” and the third category was “num_mapped_reads_in_cells” in the metrics_summary.csv.

For Parse, the values were calculated by summing the “mread_count” columns of the “cell_metadata.csv” for the unfiltered and filtered matrices to produce the second and third values respectively.

For Scale, the second category was calculated by summing the “uniquePassingReads” column of the “allcells.csv” for each sample. The third category was calculated by summing the “uniquePassingReads” column only for rows where the “pass” column was “TRUE”.

### Sequencing saturation

Not all pipelines provided saturation curves for gene and UMI recovery. For those that did, it was not always clear whether data was downsampled to a target read depth before or after any filtering. Additionally, all pipeline summaries included a sequencing saturation metric, but the calculation was not consistent across vendors. To ensure consistency across platforms, we generated our own saturation curves and metrics.

Saturation curves were generated by downsampling fastqs using Seqtk to target reads per cell based on the number of cells detected in the original run [25]. We downsampled our original 25,000 reads/cell data to a range of targets from 1,000 to 22,500 reads/cell. The number of cells was fixed in each pipeline’s parameters to enforce the target reads per cell throughout processing. We did not perform additional cell filtering of the pipeline outputs. Median genes and UMIs per cell were recorded from the pipeline summaries.

We adopted the approach of [26] to assess sequencing saturation, where saturation was quantified by fitting a model for the relationship of read depth to median gene gene/transcript count using the Michaelis-Menten equation. This model allowed us to predict the maximal values for gene and transcript recovery and calculate the number of reads needed to reach half the theoretical maximal yield. We report this metric as rd50.

### Filtering count matrices

Count matrices output by vendor pipelines were imported into R for additional processing. Ambient RNA contamination was estimated using SoupX [27], but count matrices were not adjusted to correct for ambient RNA to avoid potentially biased transformation of the data. The count matrices were imported into Seurat V5 [28]. Multiplet detection and removal was performed using scDblFinder with modified settings for unknown expected multiplet rates (dbr.sd=1) [22]. Unique molecule counts, feature counts, ribosomal read fraction, and mitochondrial read fraction data were calculated for each cell via Seurat.

The PercentageFeatureSet function was applied to flag mitochondrial and ribosomal genes, identified by gene symbols starting with “MT-” for mitochondrial genes and “RPS”/“RPL” for ribosomal proteins. Each sample was then filtered with loose quality metric thresholds reflecting standard experimental procedures intended to eliminate low quality cells. Cells with greater than 30,000 unique transcripts, greater than 10,000 unique genes, greater than 15% mitochondrial reads, or greater than 50% ribosomal reads were removed. Cells with less than 1,000 unique transcripts or less than 200 unique genes were removed. For each kit, genes detected in fewer than 10 cells in 3 or more samples were excluded for all samples in the kit.

### Categorizing features

Biological function and categorization for aligned biotypes was pulled from Ensembl via biomaRt [29]. Genes with “protein coding” biotypes were separated based on their name. Features with prefixes of “AC” (accession numbers), “LOC” (genomic location), “orf” (open reading frame), or “ENSG” (ensembl ID) were classified as “unidentified” features, while other gene symbols were considered “identified”. Protein coding features starting with “MT” or with a feature biotype of mitochondrial were labelled mitochondrial genes. Protein coding features starting with “RPL” or “RPS” or with a feature biotype of ribosomal were labelled ribosomal genes. Long non-coding RNA (lncRNA) were identified by ensembl biotype. Any remaining features, including uncommon RNA or features with multiple biotypes, were binned as “other”.

### Kit level cell clustering

Clustering was performed with merged sample-level matrices for each kit. Filtered count matrices were log-normalized and scaled via Seurat. A subset of 2000 highly variable genes identified by Seurat’s FindVariableFeatures function were selected as input for dimensional reduction. Principal component analysis (PCA) was calculated on these genes to generate 50 PCs. Elbow plots were reviewed for each sample to ensure that the majority of variance was captured in the first 20 components. The first 20 PCs were used to generate nearest neighbor graphs. These graphs were used as input to the uniform manifold approximation and projection (UMAP) algorithm for 2-dimensional visualization of cell variance. UMAP was performed by Seurat’s RunUMAP function with default arguments. The Leiden algorithm was implemented using iGraph [30] on the nearest neighbor graphs to identify cell clusters. Resolutions were selected to produce 30 clusters (excluding singletons).

### Cell Type Annotation

To accurately annotate cell types, we applied a combination of reference-based prediction and manual annotation approaches using canonical marker genes. SingleR [31] was used to align the expression profiles of our dataset with annotated bulk RNA expression profiles [32], [33]. An additional reference mapping approach was implemented via Seurat with a single-cell reference [34]. These reference-based methods annotated cell types based on comprehensive whole-transcriptome gene expression similarities. The resulting labels served as the foundational assignments for each cell type.

In addition to the reference-based assignments, we manually reviewed and annotated cells using canonical and empirical marker genes from cellxgene to further verify and refine cell type identities (see Supplemental Table 3 for marker genes) [35]. We implemented a consensus-based approach to resolve discrepancies between the reference-based and manual annotations. In cases where there was a discrepancy, we deferred to the identity suggested by canonical marker gene expression. For example, if a cell was annotated as a CD4+ T cell by SingleR but did not show strong expression of CD4 or CD8, we assigned it as an unresolved T cell. When annotation methods yielded conflicting results or when cells exhibited mixed marker expression (e.g., markers for both T and B cells), we marked these cells unidentified and excluded them from further analysis.

Cell type composition was modeled using scCODA [36].

### Cell type module scoring

Using the same set of canonical marker genes used for cell type annotation, we quantified the collective expression of these genes in their respective cell types using UCell’s gene set scoring algorithm [37]. Briefly, the method assesses if the expression of selected genes is higher than other genes using the Mann-Whitney U test. A higher score indicates the genes are collectively more highly expressed than would be expected by chance. Module scores are calculated on a per-cell basis, then averaged at the cluster level.

### Earth mover’s distance

Earth mover’s distance was calculated between replicates within kits on a 2-dimensional kernel density estimate of single-cells in shared UMAP space. Samples were downsampled to 8000 cells and reprocessed with the normalization and dimensional reduction workflow used previously (see “Kit level cell clustering” in Methods). The UMAP coordinates were scaled and passed to “MASS::kde2d()” [38]. The resulting density distributions for each replicate were passed to “emdist::emd2d()” [39]. The downsampling and calculation for each sample were bootstrapped 25 times.

### Pseudobulking and differential gene expression

Single-cell data for each sample was grouped by high-level cell type annotation (T cell, B cell, or monocyte, other populations not included), downsampled to 2500 cells per celltype, then aggregated using ‘Seurat::AggregateExpression()’ . Pseudobulk samples were grouped by kit and modeled using DESeq2 for differential expression testing with a design formula of ‘0 + celltype + Individual’ [40]. P values were adjusted using a false discovery rate of 0.05. Expression variance was quantified with variancePartion [41].

Gene set enrichment analysis (GSEA) was performed on DESeq2 outputs using fGSEA [42] with the BioCarta pathways collection from MSigDB [43], [44].

## References

[1] F. Ginhoux, A. Yalin, C. A. Dutertre, and I. Amit, “Single-cell immunology: Past, present, and future,” Mar. 08, 2022, Cell Press. doi: 10.1016/j.immuni.2022.02.006.

[2] C. Ziegenhain et al., “Comparative Analysis of Single-Cell RNA Sequencing Methods,” Mol Cell, vol. 65, no. 4, pp. 631–643.e4, Feb. 2017, doi: 10.1016/j.molcel.2017.01.023.

[3] J. Ding et al., “Systematic comparison of single-cell and single-nucleus RNA-sequencing methods,” Nat Biotechnol, vol. 38, no. 6, pp. 737–746, Jun. 2020, doi: 10.1038/s41587-020-0465-8.

[4] J. M. Ashton et al., “Comparative Analysis of Single-Cell RNA Sequencing Platforms and Methods,” Dec. 2021.

[5] K. Hatje et al., “Comparison of Fixed Single Cell RNA-seq Methods to Enable Transcriptome Profiling of Neutrophils in Clinical Samples,” Aug. 17, 2024. doi: 10.1101/2024.08.14.607767.

[6] I. C. Clark et al., “Microfluidics-free single-cell genomics with templated emulsification,” Nat Biotechnol, vol. 41, no. 11, pp. 1557–1566, Nov. 2023, doi: 10.1038/s41587-023-01685-z.

[7] K. M. Fontanez et al., “Intrinsic molecular identifiers enable robust molecular counting in single-cell sequencing,” Oct. 05, 2024. doi: 10.1101/2024.10.04.616561.

[8] A. B. Rosenberg et al., “Single-cell profiling of the developing mouse brain and spinal cord with split-pool barcoding.” [Online]. Available: https://www.science.org

[9] J. Cao et al., “Comprehensive single-cell transcriptional profiling of a multicellular organism,” Aug. 2017, [Online]. Available: https://www.science.org

[10] M. O’Huallachain et al., “Ultra-high throughput single-cell analysis of proteins and RNAs by split-pool synthesis,” Commun Biol, vol. 3, no. 1, Dec. 2020, doi: 10.1038/s42003-020-0896-2.

[11] X. Chen et al., “A multi-center cross-platform single-cell RNA sequencing reference dataset,” Sci Data, vol. 8, no. 1, Dec. 2021, doi: 10.1038/s41597-021-00809-x.

[12] Y. Xie et al., “Comparative Analysis of Single-Cell RNA Sequencing Methods with and without Sample Multiplexing,” Int J Mol Sci, vol. 25, no. 7, Apr. 2024, doi: 10.3390/ijms25073828.

[13] M. De Simone et al., “A comprehensive analysis framework for evaluating commercial single-cell RNA sequencing technologies,” Nucleic Acids Res, vol. 53, no. 2, Jan. 2025, doi: 10.1093/nar/gkae1186.

[14] I. Filippov, C. S. Philip, L. Schauser, and P. Peterson, “Comparative transcriptomic analyses of thymocytes using 10x Genomics and Parse scRNA-seq technologies,” BMC Genomics, vol. 25, no. 1, p. 1069, Dec. 2024, doi: 10.1186/s12864-024-10976-x.

[15] L. Yu, Y. Cao, J. Y. H. Yang, and P. Yang, “Benchmarking clustering algorithms on estimating the number of cell types from single-cell RNA-sequencing data,” Genome Biol, vol. 23, no. 1, Dec. 2022, doi: 10.1186/s13059-022-02622-0.

[16] M. Sánchez-Carbonell et al., “Effect of methanol fixation on single-cell RNA sequencing of the murine dentate gyrus,” Front Mol Neurosci, vol. 16, 2023, doi: 10.3389/fnmol.2023.1223798.

[17] H. Van Phan, M. van Gent, N. Drayman, A. Basu, M. U. Gack, and S. Tay, “High-throughput RNA sequencing of paraformaldehyde-fixed single cells,” Nat Commun, vol. 12, no. 1, Dec. 2021, doi: 10.1038/s41467-021-25871-2.

[18] J. W. Squair et al., “Confronting false discoveries in single-cell differential expression,” Nat Commun, vol. 12, no. 1, p. 5692, Sep. 2021, doi: 10.1038/s41467-021-25960-2.

[19] J. D. Bloom, “Estimating the frequency of multiplets in single-cell RNA sequencing from cell-mixing experiments,” PeerJ, vol. 2018, no. 9, 2018, doi: 10.7717/peerj.5578.

[20] H. M. Kang et al., “Multiplexed droplet single-cell RNA-sequencing using natural genetic variation,” Nat Biotechnol, vol. 36, no. 1, pp. 89–94, Jan. 2018, doi: 10.1038/nbt.4042.

[21] L. Kuijpers, B. Hornung, M. C. G. N. van den Hout - van Vroonhoven, W. F. J. van IJcken, F. Grosveld, and E. Mulugeta, “Split Pool Ligation-based Single-cell Transcriptome sequencing (SPLiT-seq) data processing pipeline comparison,” BMC Genomics, vol. 25, no. 1, Dec. 2024, doi: 10.1186/s12864-024-10285-3.

[22] P.-L. Germain, A. Lun, C. G. Meixide, W. Macnair, and M. D. Robinson, “Doublet identification in single-cell sequencing data using scDblFinder,” F1000Res, vol. 10, p. 979, Oct. 2022, doi: 10.12688/f1000research.73600.2.

[23] Parse Biosciences, “Comparison of Evercode WT v3 and Evercode WT v2 in Human Immune Cells (PBMCs).” Accessed: May 14, 2025. [Online]. Available: https://www.parsebiosciences.com/datasets/comparison-of-evercode-wt-v3-and-evercode-wt-v2-in-human-immune-cells-pbmcs/

[24] I. Filippov, C. S. Philip, L. Schauser, and P. Peterson, “Comparative transcriptomic analyses of thymocytes using 10x Genomics and Parse scRNA-seq technologies,” BMC Genomics, vol. 25, no. 1, p. 1069, Dec. 2024, doi: 10.1186/s12864-024-10976-x.

[25] H. Li, “seqtk,” 2012, 1.3. [Online]. Available: https://github.com/lh3/seqtk

[26] M. De Simone et al., “Comparative Analysis of Commercial Single-Cell RNA Sequencing Technologies,” Jun. 18, 2024. doi: 10.1101/2024.06.18.599579.

[27] M. D. Young and S. Behjati, “SoupX removes ambient RNA contamination from droplet-based single-cell RNA sequencing data,” Gigascience, vol. 9, no. 12, p. giaa151, Oct. 2020, doi: 10.1093/gigascience/giaa151.

[28] Y. Hao et al., “Dictionary learning for integrative, multimodal and scalable single-cell analysis,” Nat Biotechnol, vol. 42, no. 2, pp. 293–304, 2024, doi: 10.1038/s41587-023-01767-y.

[29] S. Durinck et al., “BioMart and Bioconductor: a powerful link between biological databases and microarray data analysis,” Bioinformatics, vol. 21, no. 16, pp. 3439–3440, Aug. 2005, doi: 10.1093/bioinformatics/bti525.

[30] G. Csárdi et al., “igraph: Network Analysis and Visualization in R,” 2024. doi: 10.5281/zenodo.7682609.

[31] D. Aran et al., “Reference-based analysis of lung single-cell sequencing reveals a transitional profibrotic macrophage,” Nat Immunol, vol. 20, no. 2, pp. 163–172, Feb. 2019, doi: 10.1038/s41590-018-0276-y.

[32] G. Monaco et al., “RNA-Seq Signatures Normalized by mRNA Abundance Allow Absolute Deconvolution of Human Immune Cell Types,” Cell Rep, vol. 26, no. 6, pp. 1627–1640.e7, Feb. 2019, doi: 10.1016/j.celrep.2019.01.041.

[33] N. A. Mabbott, J. Baillie, H. Brown, T. C. Freeman, and D. A. Hume, “An expression atlas of human primary cells: inference of gene function from coexpression networks,” BMC Genomics, vol. 14, no. 1, p. 632, 2013, doi: 10.1186/1471-2164-14-632.

[34] M. Stoeckius et al., “Simultaneous epitope and transcriptome measurement in single cells,” Nat Methods, vol. 14, no. 9, pp. 865–868, Sep. 2017, doi: 10.1038/nmeth.4380.

[35] S. Abdulla et al., “CZ CELL×GENE Discover: A single-cell data platform for scalable exploration, analysis and modeling of aggregated data,” Nov. 02, 2023. doi: 10.1101/2023.10.30.563174.

[36] M. Büttner, J. Ostner, C. L. Müller, F. J. Theis, and B. Schubert, “scCODA is a Bayesian model for compositional single-cell data analysis,” Nat Commun, vol. 12, no. 1, p. 6876, Nov. 2021, doi: 10.1038/s41467-021-27150-6.

[37] M. Andreatta and S. J. Carmona, “UCell: Robust and scalable single-cell gene signature scoring,” Comput Struct Biotechnol J, vol. 19, pp. 3796–3798, 2021, doi: 10.1016/j.csbj.2021.06.043.

[38] W. N. Venables and B. D. Ripley, Modern Applied Statistics with S, 4th ed. New York: Springer, 2002.

[39] S. Urbanek and Y. Rubner, “emdist: Earth Mover’s Distance,” 2023, 0.3-3.

[40] M. I. Love, W. Huber, and S. Anders, “Moderated estimation of fold change and dispersion for RNA-seq data with DESeq2,” Genome Biol, vol. 15, no. 12, p. 550, Dec. 2014, doi: 10.1186/s13059-014-0550-8.

[41] G. E. Hoffman and E. E. Schadt, “variancePartition: interpreting drivers of variation in complex gene expression studies,” BMC Bioinformatics, vol. 17, no. 1, p. 483, Nov. 2016, doi: 10.1186/s12859-016-1323-z.

[42] G. Korotkevich, V. Sukhov, N. Budin, B. Shpak, M. N. Artyomov, and A. Sergushichev, “Fast gene set enrichment analysis,” Jun. 20, 2016. doi: 10.1101/060012.

[43] A. Subramanian et al., “Gene set enrichment analysis: A knowledge-based approach for interpreting genome-wide expression profiles,” Proceedings of the National Academy of Sciences, vol. 102, no. 43, pp. 15545–15550, Oct. 2005, doi: 10.1073/pnas.0506580102.

[44] A. Liberzon, A. Subramanian, R. Pinchback, H. Thorvaldsdóttir, P. Tamayo, and J. P. Mesirov, “Molecular signatures database (MSigDB) 3.0,” Bioinformatics, vol. 27, no. 12, pp. 1739–1740, Jun. 2011, doi: 10.1093/bioinformatics/btr260.

