## Supplementary Materials for "Evaluating the practical aspects and performance of commercial single-cell RNA sequencing technologies"

### Supplemental figure 1

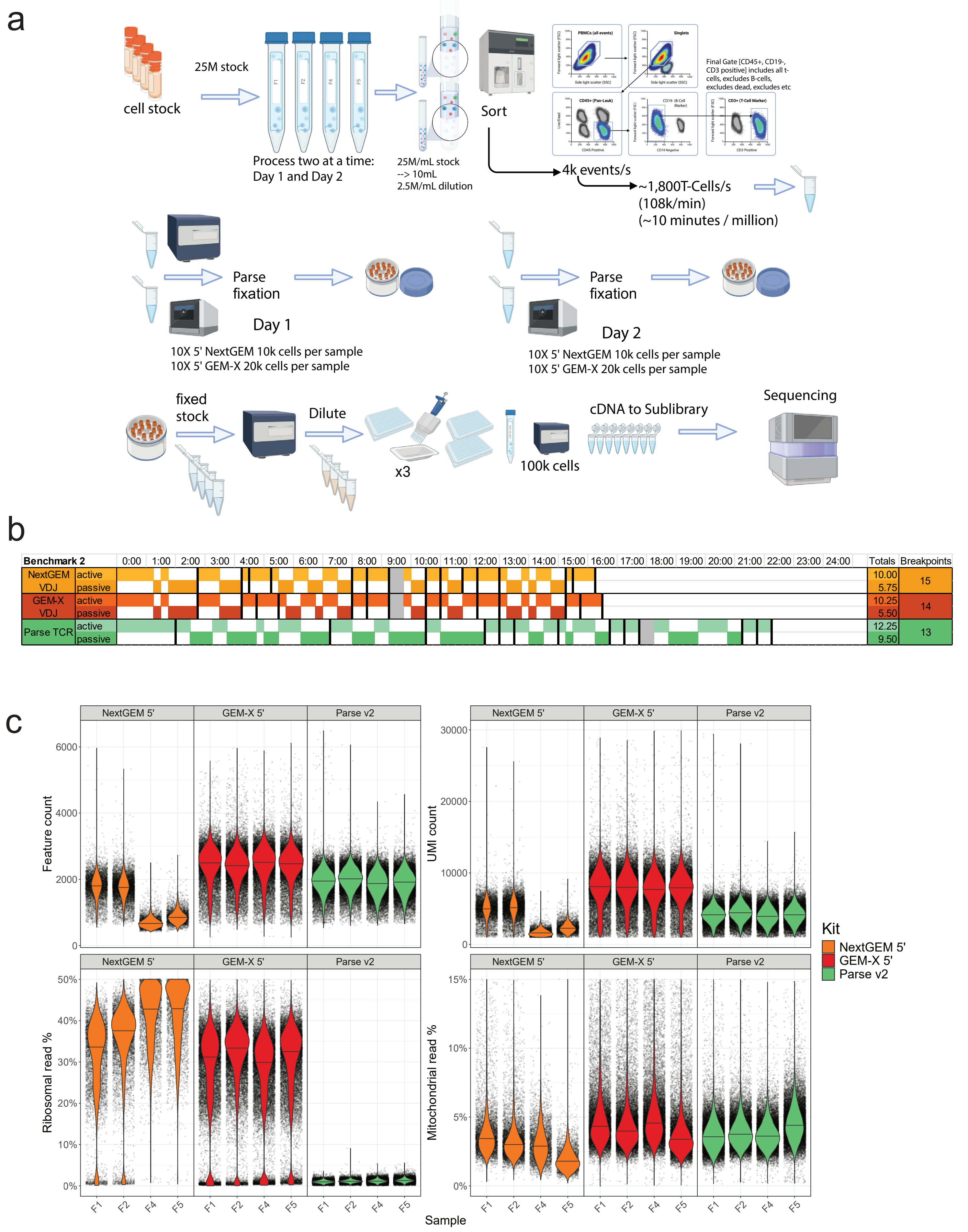

### Supplemental figure 2

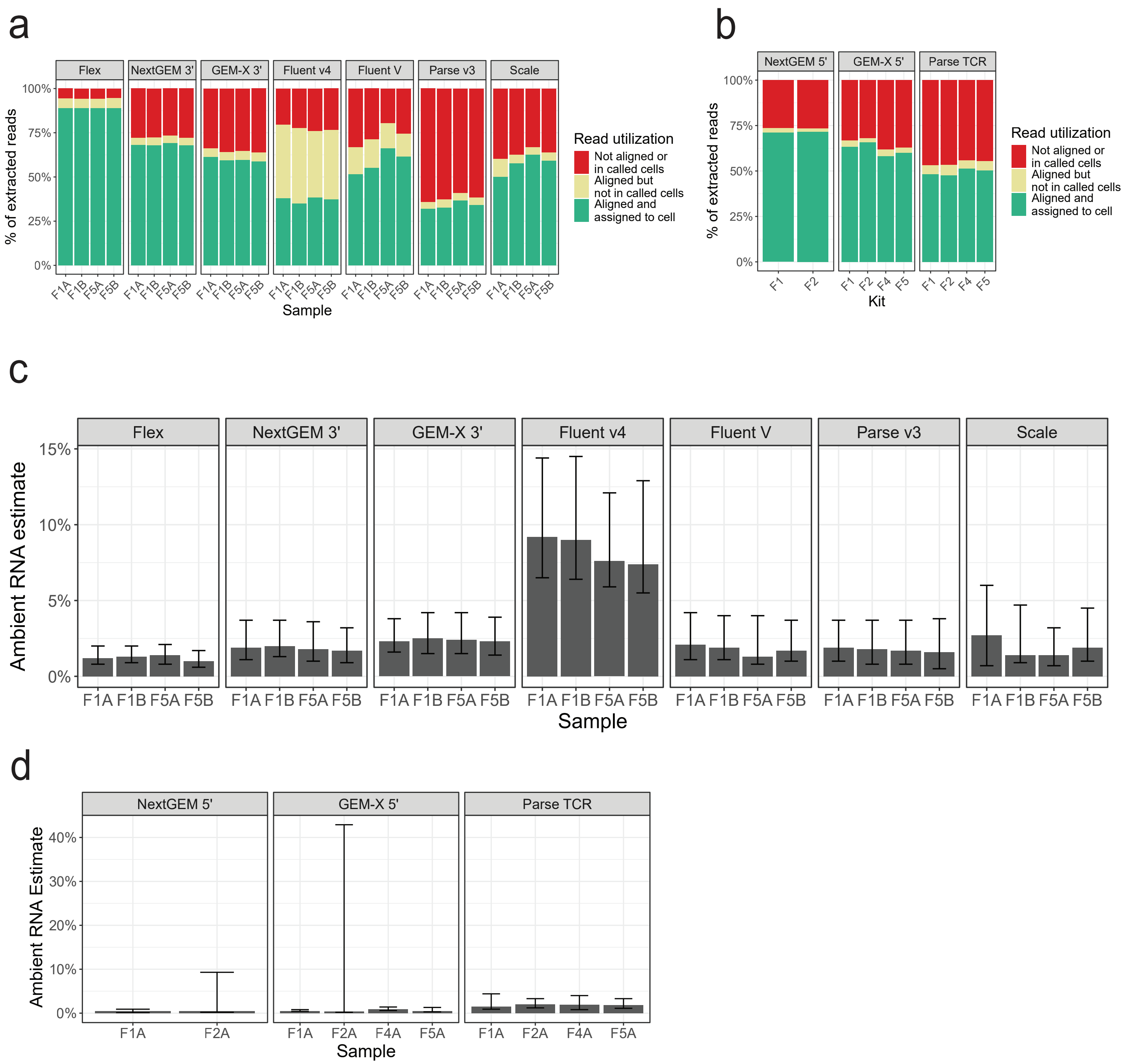

### Supplemental figure 3

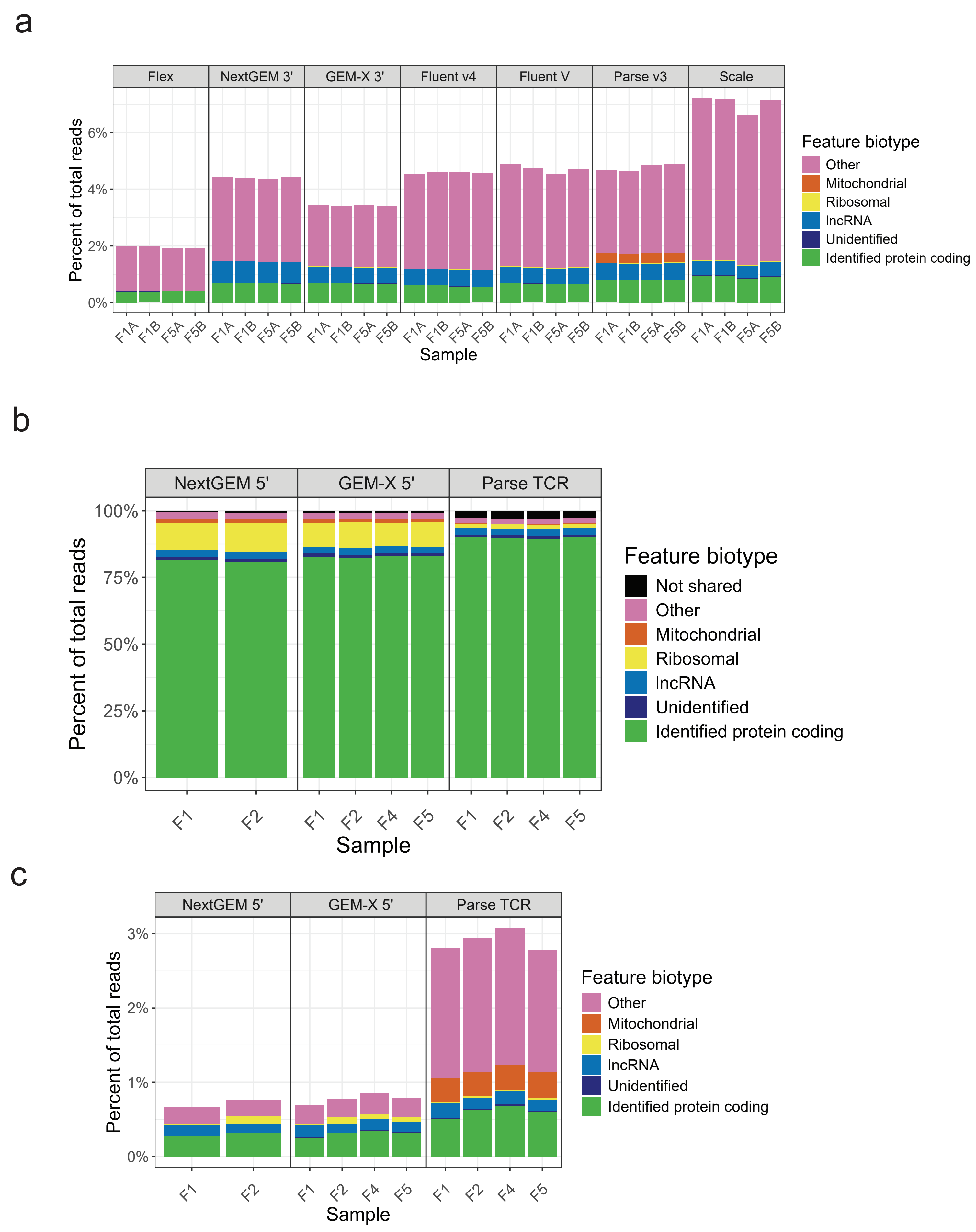

### Supplemental figure 4

#### Flex

| Cluster | Cell count | Seurat pbmcsca Label | Seurat pbmcsca Score | Seurat pbmcsca % called | Seurat pbmc3k Label | Seurat pbmc3k Score | Seurat pbmc3k % called | SingleR Mona Label | SingleR Mona Score | SingleR Mona % called | SingleR HPCA Label | SingleR HPCA Score | SingleR HPCA % called |
|---|---|---|---|---|---|---|---|---|---|---|---|---|---|
| 0 | 2602 | CD4+ T | 0.99 | 100.00 | CD4+ T | 0.98 | 100.00 | CD4+ T | 0.29 | 97.89 | T | 0.75 | 99.96 |
| 1 | 2028 | Classical monocyte | 1.00 | 99.16 | Classical monocyte | 0.99 | 96.65 | Monocyte | 0.07 | 87.28 | Monocyte | 0.07 | 90.38 |
| 2 | 2017 | NK | 0.91 | 96.93 | NK | 0.98 | 99.26 | CD8+ T | 0.10 | 69.96 | NK | 0.48 | 99.11 |
| 3 | 2002 | CD8+ T | 0.84 | 97.05 | CD8+ T | 0.77 | 92.76 | CD8+ T | 0.10 | 71.93 | T | 0.15 | 89.46 |
| 4 | 1981 | B | 1.00 | 99.04 | B | 1.00 | 98.94 | B | 0.09 | 93.99 | B | 0.07 | 97.32 |
| 5 | 1973 | CD4+ T | 0.96 | 100.00 | CD4+ T | 1.00 | 100.00 | CD4+ T | 0.25 | 94.98 | T | 0.69 | 99.44 |
| 6 | 1920 | CD8+ T | 0.73 | 74.01 | CD8+ T | 0.69 | 59.17 | CD8+ T | 0.10 | 98.02 | NK | 0.12 | 49.64 |
| 7 | 1872 | CD4+ T | 0.91 | 94.07 | CD4+ T | 0.97 | 95.19 | CD4+ T | 0.19 | 92.41 | T | 0.56 | 97.65 |
| 8 | 1681 | Classical monocyte | 1.00 | 99.82 | Classical monocyte | 0.98 | 99.52 | Monocyte | 0.07 | 93.16 | Monocyte | 0.08 | 93.63 |
| 9 | 1649 | CD4+ T | 1.00 | 100.00 | CD4+ T | 0.96 | 100.00 | CD4+ T | 0.25 | 96.06 | T | 0.70 | 99.76 |
| 10 | 1634 | B | 1.00 | 100.00 | B | 1.00 | 99.82 | B | 0.10 | 96.88 | B | 0.07 | 98.10 |
| 11 | 1576 | CD4+ T | 0.81 | 83.38 | CD4+ T | 0.96 | 91.88 | CD8+ T | 0.00 | 97.02 | T | 0.50 | 96.83 |
| 12 | 1519 | CD4+ T | 1.00 | 100.00 | CD4+ T | 0.91 | 100.00 | CD4+ T | 0.23 | 94.54 | T | 0.66 | 99.01 |
| 13 | 1446 | CD4+ T | 1.00 | 99.86 | CD4+ T | 0.94 | 100.00 | CD8+ T | 0.06 | 97.79 | T | 0.73 | 99.17 |
| 14 | 1391 | CD8+ T | 0.77 | 83.68 | CD8+ T | 0.83 | 91.59 | CD8+ T | 0.00 | 51.62 | T | 0.24 | 94.25 |
| 15 | 1254 | Non−classical monocyte | 0.97 | 96.97 | Non−classical monocyte | 1.00 | 98.56 | Monocyte | 0.08 | 85.09 | Monocyte | 0.08 | 94.42 |
| 16 | 1242 | CD8+ T | 0.75 | 84.30 | CD8+ T | 0.80 | 79.23 | CD8+ T | 0.10 | 90.02 | T | 0.20 | 89.86 |
| 17 | 1067 | CD4+ T | 1.00 | 100.00 | CD4+ T | 0.95 | 100.00 | CD8+ T | 0.05 | 99.44 | T | 0.69 | 99.91 |
| 18 | 998 | CD4+ T | 0.99 | 100.00 | CD4+ T | 0.84 | 100.00 | CD4+ T | 0.22 | 91.18 | T | 0.68 | 98.90 |
| 19 | 879 | CD4+ T | 0.93 | 99.54 | CD4+ T | 0.99 | 100.00 | CD4+ T | 0.19 | 92.72 | T | 0.38 | 97.84 |
| 20 | 805 | CD4+ T | 0.95 | 100.00 | CD4+ T | 1.00 | 100.00 | CD4+ T | 0.23 | 94.41 | T | 0.65 | 98.88 |
| 21 | 781 | Classical monocyte | 0.99 | 61.08 | Classical monocyte | 0.92 | 52.50 | Monocyte | 0.07 | 52.75 | Monocyte | 0.08 | 94.49 |
| 22 | 658 | NK | 0.71 | 74.16 | NK | 0.84 | 78.12 | CD8+ T | 0.10 | 62.16 | NK | 0.32 | 95.14 |
| 23 | 285 | Megakaryocyte | 1.00 | 74.39 | Megakaryocyte | 0.95 | 74.39 | Lymphocyte progenitor | 0.07 | 41.40 | Megakaryocyte | 0.11 | 65.96 |
| 24 | 144 | pDC | 0.54 | 77.78 | Dendritic | 0.90 | 100.00 | Dendritic | 0.15 | 99.31 | Myeloid | 0.00 | 57.64 |
| 25 | 114 | CD4+ T | 0.98 | 88.60 | CD4+ T | 0.99 | 91.23 | CD4+ T | 0.25 | 73.68 | T | 0.64 | 97.37 |
| 26 | 96 | CD4+ T | 0.73 | 40.62 | CD4+ T | 0.62 | 52.08 | CD4+ T | 0.06 | 20.83 | T | 0.12 | 30.21 |
| 27 | 61 | CD4+ T | 0.60 | 57.38 | CD4+ T | 0.62 | 95.08 | Granulocyte | 0.08 | 45.90 | Granulocyte | 0.19 | 32.79 |
| 28 | 35 | Classical monocyte | 0.89 | 94.29 | Classical monocyte | 0.66 | 42.86 | Monocyte | 0.26 | 68.57 | Monocyte | 0.11 | 88.57 |
| 29 | 29 | CD8+ T | 0.65 | 51.72 | CD4+ T | 0.26 | 62.07 | CD8+ T | 0.10 | 79.31 | NK | 0.62 | 62.07 |

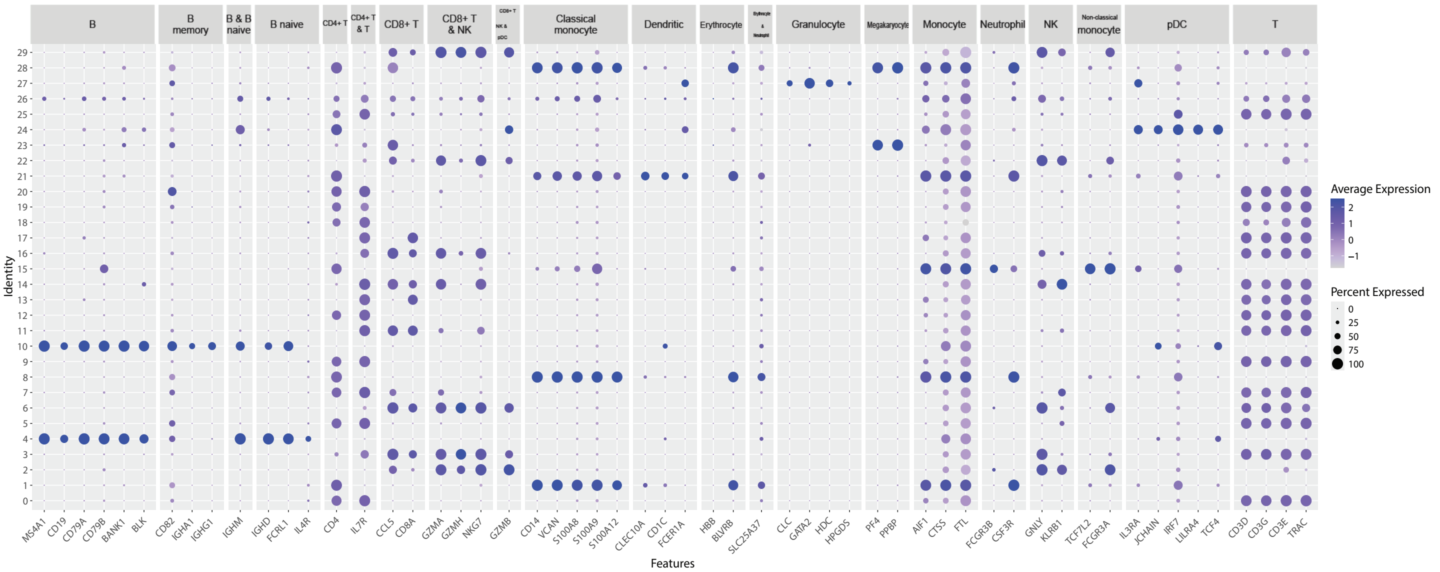

Leiden clustering

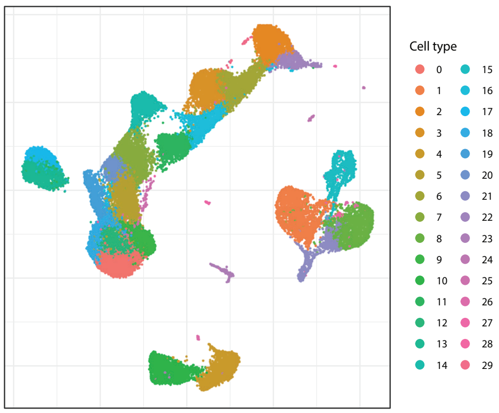

Final annotations

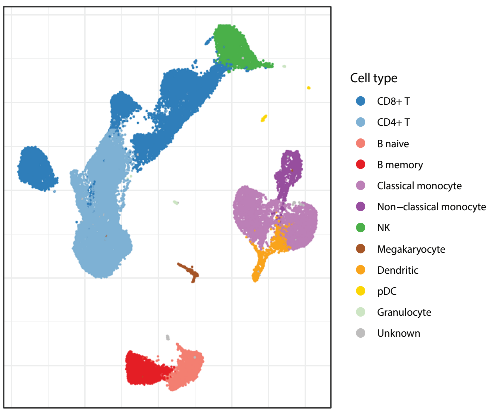

| Cell label | Count | Proportion |
|---|---|---|
| CD4+ T | 12411 | 0.35 |
| CD8+ T | 10673 | 0.30 |
| Classical monocyte | 3744 | 0.10 |
| NK | 2675 | 0.07 |
| B naive | 1981 | 0.06 |
| B memory | 1634 | 0.05 |
| Non−classical monocyte | 1254 | 0.04 |
| Dendritic | 781 | 0.02 |
| Megakaryocyte | 285 | 0.01 |
| pDC | 144 | 0.00 |
| Unknown | 96 | 0.00 |
| Granulocyte | 61 | 0.00 |

### NextGEM 3′

| | | Seurat | | | | | | SingleR | | | | | |
|---|---|---|---|---|---|---|---|---|---|---|---|---|---|
| | | pbmcsca | | | pbmc3k | | | Mona | | | HPCA | | |
| Cluster | Cell count | Label | Score | % called | Label | Score | % called | Label | Score | % called | Label | Score | % called |
| 0 | 4010 | CD4+ T | 1.00 | 99.83 | CD4+ T | 0.70 | 100.00 | CD4+ T | 0.17 | 83.42 | T | 0.21 | 98.33 |
| 1 | 3370 | CD4+ T | 1.00 | 99.97 | CD4+ T | 0.93 | 100.00 | CD4+ T | 0.13 | 72.64 | T | 0.16 | 96.14 |
| 2 | 2958 | NK | 0.79 | 71.60 | NK | 0.97 | 89.28 | NK | 0.00 | 72.24 | NK | 0.21 | 94.56 |
| 3 | 2817 | CD4+ T | 1.00 | 99.89 | CD4+ T | 0.86 | 100.00 | CD4+ T | 0.14 | 75.97 | T | 0.18 | 97.62 |
| 4 | 2791 | CD8+ T | 0.87 | 97.85 | CD8+ T | 0.92 | 96.74 | T | 0.08 | 34.65 | T | 0.10 | 73.84 |
| 5 | 2580 | Classical monocyte | 1.00 | 99.65 | Classical monocyte | 0.99 | 99.65 | Monocyte | 0.06 | 77.91 | Monocyte | 0.02 | 59.77 |
| 6 | 2547 | B | 1.00 | 99.92 | B | 1.00 | 99.92 | B | 0.08 | 80.64 | B | 0.07 | 90.58 |
| 7 | 2492 | CD8+ T | 0.81 | 87.04 | CD8+ T | 0.85 | 85.51 | NK | 0.00 | 41.97 | T | 0.09 | 51.93 |
| 8 | 2405 | CD4+ T | 0.97 | 99.42 | CD4+ T | 0.83 | 100.00 | CD4+ T | 0.16 | 82.62 | T | 0.19 | 97.21 |
| 9 | 2382 | CD4+ T | 0.91 | 96.73 | CD4+ T | 0.86 | 95.68 | CD4+ T | 0.10 | 64.69 | T | 0.15 | 94.21 |
| 10 | 2258 | B | 1.00 | 99.91 | B | 1.00 | 99.91 | B | 0.09 | 86.05 | B | 0.07 | 90.57 |
| 11 | 2239 | CD4+ T | 0.78 | 64.63 | CD4+ T | 0.81 | 66.95 | T | 0.08 | 41.76 | T | 0.14 | 92.27 |
| 12 | 1908 | CD4+ T | 1.00 | 99.84 | CD4+ T | 0.80 | 100.00 | CD8+ T | 0.07 | 55.66 | T | 0.16 | 97.12 |
| 13 | 1882 | Classical monocyte | 1.00 | 95.86 | Classical monocyte | 0.95 | 90.91 | Monocyte | 0.06 | 76.46 | Monocyte | 0.01 | 72.64 |
| 14 | 1625 | Classical monocyte | 1.00 | 99.94 | Classical monocyte | 1.00 | 99.75 | Monocyte | 0.00 | 70.09 | Monocyte | 0.02 | 45.91 |
| 15 | 1452 | CD4+ T | 1.00 | 100.00 | CD4+ T | 0.89 | 100.00 | CD8+ T | 0.07 | 56.13 | T | 0.15 | 96.76 |
| 16 | 1435 | CD8+ T | 0.70 | 56.79 | CD8+ T | 0.89 | 92.61 | T | 0.08 | 34.77 | T | 0.13 | 89.41 |
| 17 | 1398 | Non−classical monocyte | 0.88 | 94.92 | Non−classical monocyte | 1.00 | 94.71 | Monocyte | 0.06 | 80.26 | Monocyte | 0.00 | 65.67 |
| 18 | 1228 | CD4+ T | 0.96 | 98.29 | CD4+ T | 0.91 | 100.00 | CD4+ T | 0.10 | 67.67 | T | 0.14 | 94.87 |
| 19 | 1075 | CD4+ T | 0.91 | 94.98 | CD4+ T | 0.92 | 94.14 | CD4+ T | 0.10 | 66.70 | T | 0.16 | 94.88 |
| 20 | 983 | CD4+ T | 0.96 | 99.29 | CD4+ T | 0.96 | 99.69 | CD4+ T | 0.13 | 76.91 | T | 0.19 | 95.93 |
| 21 | 971 | CD4+ T | 0.89 | 93.92 | CD4+ T | 0.87 | 99.79 | CD4+ T | 0.11 | 68.59 | T | 0.12 | 88.36 |
| 22 | 618 | Dendritic | 0.72 | 78.16 | Dendritic | 0.95 | 91.42 | Dendritic | 0.07 | 49.19 | Monocyte | 0.05 | 59.55 |
| 23 | 474 | Megakaryocyte | 1.00 | 47.89 | Megakaryocyte | 0.94 | 48.73 | B | 0.05 | 39.24 | T | 0.12 | 50.84 |
| 24 | 395 | CD8+ T | 0.73 | 77.72 | CD8+ T | 0.89 | 93.92 | NK | 0.00 | 58.73 | T | 0.11 | 77.97 |
| 25 | 192 | CD4+ T | 0.51 | 38.54 | NK | 0.81 | 58.33 | NK | 0.00 | 50.00 | NK | 0.19 | 77.60 |
| 26 | 156 | pDC | 0.59 | 92.95 | Dendritic | 0.78 | 96.15 | Dendritic | 0.11 | 96.15 | Myeloid | 0.00 | 35.26 |
| 27 | 147 | CD4+ T | 0.92 | 95.92 | CD4+ T | 0.77 | 92.52 | T | 0.08 | 31.97 | T | 0.13 | 87.76 |
| 28 | 102 | Classical monocyte | 0.91 | 87.25 | Classical monocyte | 0.77 | 67.65 | Monocyte | 0.07 | 50.98 | Monocyte | 0.06 | 60.78 |
| 29 | 43 | Classical monocyte | 0.46 | 100.00 | CD4+ T | 0.59 | 100.00 | B | 0.00 | 83.72 | BM | 0.01 | 60.47 |

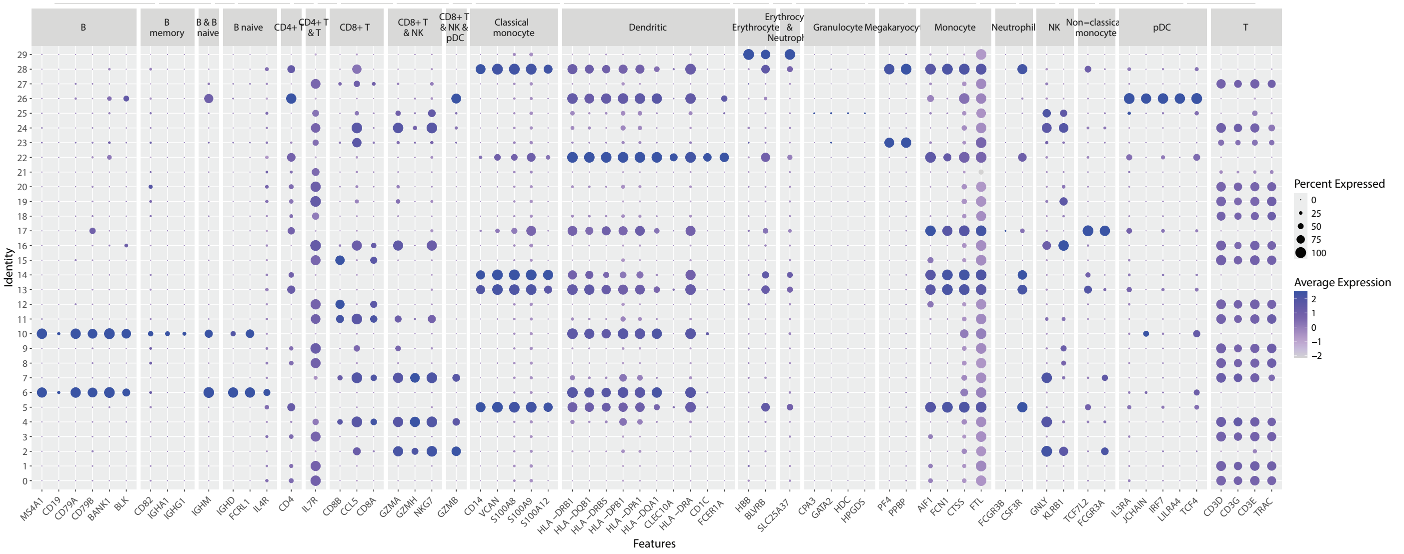

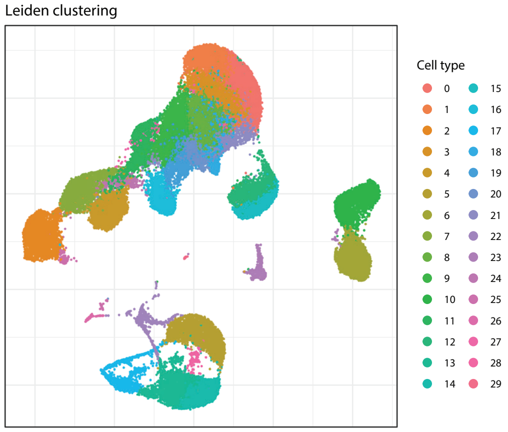

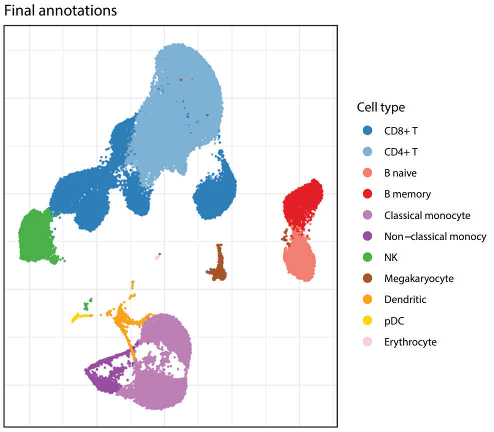

| Cell label | Count | Proportion |
|---|---|---|
| CD4+ T | 19241 | 0.39 |
| CD8+ T | 12859 | 0.26 |
| Classical monocyte | 6189 | 0.13 |
| NK | 3150 | 0.06 |
| B naive | 2547 | 0.05 |
| B memory | 2258 | 0.05 |
| Non−classical monocyte | 1398 | 0.03 |
| Dendritic | 618 | 0.01 |
| Megakaryocyte | 474 | 0.01 |
| pDC | 156 | 0.00 |
| Erythrocyte | 43 | 0.00 |

### GEM-X 3′

| | | Seurat | | | | | | SingleR | | | | | |
|---|---|---|---|---|---|---|---|---|---|---|---|---|---|
| | | pbmcsca | | | pbmc3k | | | Mona | | | HPCA | | |
| Cluster | Cell count | Label | Score | % called | Label | Score | % called | Label | Score | % called | Label | Score | % called |
| 0 | 14052 | CD4+ T | 1.00 | 100.00 | CD4+ T | 0.72 | 100.00 | CD4+ T | 0.15 | 77.33 | T | 0.24 | 99.57 |
| 1 | 8654 | CD4+ T | 0.92 | 95.86 | CD4+ T | 0.99 | 98.20 | CD4+ T | 0.12 | 70.29 | T | 0.20 | 98.65 |
| 2 | 7168 | CD4+ T | 0.91 | 93.96 | CD4+ T | 0.87 | 99.29 | CD4+ T | 0.07 | 46.00 | T | 0.12 | 90.33 |
| 3 | 7078 | Classical monocyte | 1.00 | 99.99 | Classical monocyte | 1.00 | 99.82 | Monocyte | 0.05 | 89.29 | Monocyte | 0.00 | 70.91 |
| 4 | 6520 | Classical monocyte | 1.00 | 99.57 | Classical monocyte | 0.94 | 98.01 | Monocyte | 0.05 | 89.46 | Monocyte | 0.00 | 71.58 |
| 5 | 5768 | NK | 0.85 | 78.59 | NK | 0.96 | 92.44 | NK | 0.00 | 83.44 | NK | 0.31 | 96.27 |
| 6 | 5724 | CD4+ T | 1.00 | 100.00 | CD4+ T | 0.74 | 100.00 | CD4+ T | 0.14 | 75.98 | T | 0.25 | 99.74 |
| 7 | 5674 | CD8+ T | 0.86 | 97.30 | CD8+ T | 0.86 | 93.14 | T | 0.01 | 53.03 | T | 0.11 | 81.14 |
| 8 | 5566 | B | 1.00 | 100.00 | B | 1.00 | 100.00 | B | 0.09 | 83.27 | B | 0.07 | 92.63 |
| 9 | 5477 | CD8+ T | 0.82 | 86.12 | CD8+ T | 0.78 | 78.46 | T | 0.01 | 54.06 | T | 0.10 | 56.95 |
| 10 | 5131 | CD4+ T | 0.83 | 70.80 | CD4+ T | 0.93 | 89.42 | T | 0.01 | 60.67 | T | 0.17 | 97.19 |
| 11 | 5011 | B | 1.00 | 100.00 | B | 1.00 | 99.94 | B | 0.09 | 87.13 | B | 0.07 | 90.46 |
| 12 | 4575 | CD4+ T | 0.98 | 99.93 | CD4+ T | 0.97 | 100.00 | CD4+ T | 0.14 | 79.48 | T | 0.23 | 99.23 |
| 13 | 4054 | CD8+ T | 0.78 | 86.95 | CD8+ T | 0.80 | 86.85 | NK | 0.00 | 45.58 | T | 0.15 | 93.49 |
| 14 | 3581 | CD4+ T | 1.00 | 99.94 | CD4+ T | 0.66 | 100.00 | T | 0.01 | 75.34 | T | 0.21 | 99.47 |
| 15 | 3273 | CD4+ T | 1.00 | 100.00 | CD4+ T | 0.66 | 100.00 | T | 0.01 | 77.05 | T | 0.20 | 99.08 |
| 16 | 3262 | CD4+ T | 0.99 | 99.91 | CD4+ T | 0.93 | 100.00 | CD4+ T | 0.15 | 81.85 | T | 0.24 | 99.69 |
| 17 | 2955 | Non−classical monocyte | 0.88 | 87.01 | Non−classical monocyte | 1.00 | 89.75 | Monocyte | 0.06 | 87.68 | Monocyte | 0.00 | 81.96 |
| 18 | 2808 | CD4+ T | 0.97 | 99.93 | CD4+ T | 0.94 | 100.00 | CD4+ T | 0.11 | 68.13 | T | 0.19 | 98.50 |
| 19 | 2434 | Megakaryocyte | 1.00 | 68.00 | Megakaryocyte | 0.97 | 68.00 | B | 0.07 | 45.81 | Megakaryocyte | 0.07 | 50.58 |
| 20 | 2081 | Classical monocyte | 0.93 | 64.30 | Classical monocyte | 0.94 | 26.86 | Monocyte | 0.02 | 47.62 | Granulocyte | 0.10 | 36.71 |
| 21 | 1962 | CD8+ T | 0.69 | 64.53 | CD4+ T | 0.69 | 41.54 | NK | 0.00 | 55.86 | NK | 0.16 | 53.01 |
| 22 | 1821 | B | 1.00 | 78.91 | B | 1.00 | 73.92 | B | 0.07 | 57.11 | B | 0.06 | 43.05 |
| 23 | 1600 | Dendritic | 0.66 | 59.81 | Dendritic | 0.96 | 81.00 | Monocyte | 0.00 | 44.44 | Monocyte | 0.00 | 58.13 |
| 24 | 501 | Classical monocyte | 0.48 | 74.05 | CD4+ T | 0.69 | 99.60 | B | 0.00 | 47.31 | BM | 0.01 | 44.31 |
| 25 | 440 | CD4+ T | 0.68 | 72.95 | CD4+ T | 0.69 | 79.77 | NK | 0.00 | 36.59 | T | 0.11 | 81.82 |
| 26 | 370 | pDC | 0.54 | 60.54 | Dendritic | 0.80 | 75.68 | Dendritic | 0.10 | 80.81 | Myeloid | 0.00 | 38.38 |
| 27 | 205 | Classical monocyte | 0.90 | 93.17 | Non−classical monocyte | 0.69 | 45.85 | Monocyte | 0.06 | 63.90 | Monocyte | 0.06 | 69.27 |
| 28 | 107 | Classical monocyte | 0.70 | 68.22 | CD4+ T | 0.60 | 74.77 | Granulocyte | 0.08 | 88.79 | Granulocyte | 0.06 | 51.40 |
| 29 | 62 | CD4+ T | 0.79 | 77.42 | CD4+ T | 0.93 | 95.16 | NK | 0.00 | 32.26 | NK | 0.11 | 59.68 |

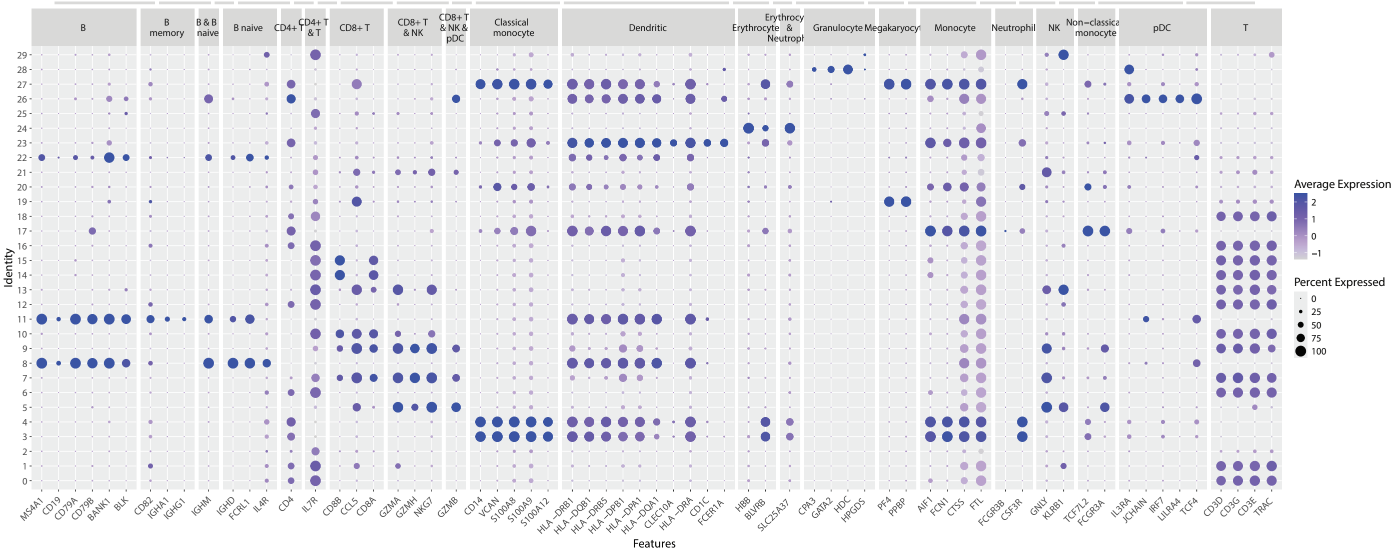

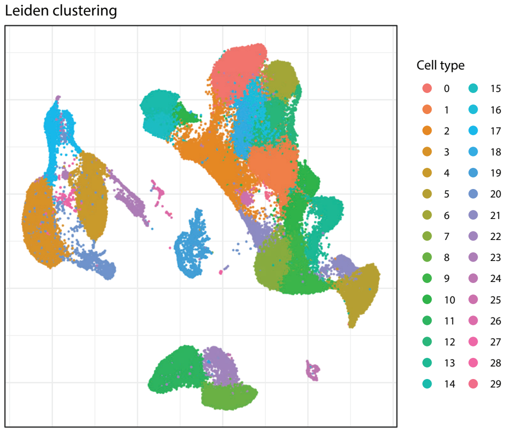

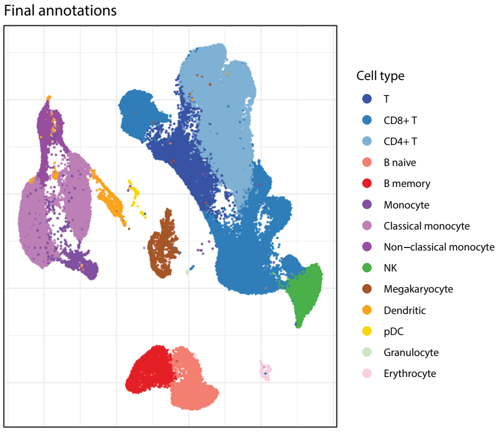

| Cell label | Count | Proportion |
|---|---|---|
| CD4+ T | 39075 | 0.33 |
| CD8+ T | 29152 | 0.25 |
| Classical monocyte | 13803 | 0.12 |
| T | 7670 | 0.07 |
| B naive | 7387 | 0.06 |
| NK | 5768 | 0.05 |
| B memory | 5011 | 0.04 |
| Non−classical monocyte | 2955 | 0.03 |
| Megakaryocyte | 2434 | 0.02 |
| Monocyte | 2081 | 0.02 |
| Dendritic | 1600 | 0.01 |
| Erythrocyte | 501 | 0.00 |
| pDC | 370 | 0.00 |
| Granulocyte | 107 | 0.00 |

### Fluent v4

| | | Seurat | | | | | | SingleR | | | | | |
|---|---|---|---|---|---|---|---|---|---|---|---|---|---|
| | | pbmcsca | | | pbmc3k | | | Mona | | | HPCA | | |
| Cluster | Cell count | Label | Score | % called | Label | Score | % called | Label | Score | % called | Label | Score | % called |
| 0 | 4493 | NK | 0.79 | 65.57 | NK | 0.89 | 72.71 | Granulocyte | 0.00 | 27.35 | NK | 0.11 | 79.10 |
| 1 | 4311 | CD8+ T | 0.90 | 96.10 | CD8+ T | 0.86 | 88.66 | Granulocyte | 0.02 | 26.68 | T | 0.08 | 54.49 |
| 2 | 3918 | CD4+ T | 0.99 | 99.72 | CD4+ T | 0.94 | 100.00 | CD4+ T | 0.01 | 39.77 | T | 0.10 | 73.05 |
| 3 | 3481 | CD4+ T | 0.99 | 99.57 | CD4+ T | 0.98 | 99.97 | CD4+ T | 0.01 | 36.68 | T | 0.09 | 71.07 |
| 4 | 3007 | CD8+ T | 0.84 | 82.21 | CD8+ T | 0.85 | 84.84 | CD8+ T | 0.00 | 23.11 | T | 0.08 | 45.73 |
| 5 | 2995 | B | 1.00 | 99.97 | B | 1.00 | 99.87 | B | 0.00 | 27.88 | B | 0.06 | 67.41 |
| 6 | 2884 | CD4+ T | 0.75 | 58.95 | CD4+ T | 0.66 | 79.02 | Granulocyte | 0.02 | 27.98 | T | 0.09 | 62.14 |
| 7 | 2810 | B | 1.00 | 99.75 | B | 1.00 | 99.68 | B | 0.00 | 26.37 | B | 0.07 | 68.01 |
| 8 | 2776 | CD4+ T | 0.99 | 99.14 | CD4+ T | 0.96 | 99.96 | T | 0.01 | 27.92 | T | 0.09 | 68.48 |
| 9 | 2738 | CD8+ T | 0.73 | 63.11 | CD8+ T | 0.72 | 70.38 | Granulocyte | 0.04 | 27.17 | T | 0.08 | 56.61 |
| 10 | 2561 | CD4+ T | 0.84 | 89.65 | CD4+ T | 0.73 | 94.38 | Granulocyte | 0.02 | 27.33 | T | 0.09 | 65.52 |
| 11 | 2516 | Classical monocyte | 1.00 | 98.81 | Classical monocyte | 0.96 | 92.13 | Monocyte | 0.00 | 32.11 | Monocyte | 0.01 | 40.46 |
| 12 | 2447 | CD4+ T | 0.90 | 94.97 | CD4+ T | 0.74 | 98.86 | CD4+ T | 0.01 | 36.13 | T | 0.09 | 70.17 |
| 13 | 2269 | CD4+ T | 0.96 | 97.93 | CD4+ T | 0.92 | 100.00 | CD4+ T | 0.01 | 35.04 | T | 0.08 | 64.74 |
| 14 | 2267 | CD4+ T | 0.90 | 96.12 | CD4+ T | 0.75 | 98.68 | CD4+ T | 0.01 | 38.20 | T | 0.10 | 72.25 |
| 15 | 2198 | CD4+ T | 0.99 | 99.68 | CD4+ T | 0.96 | 100.00 | CD4+ T | 0.01 | 40.67 | T | 0.09 | 74.39 |
| 16 | 1999 | CD4+ T | 0.69 | 58.93 | CD4+ T | 0.75 | 76.14 | Granulocyte | 0.06 | 30.67 | HSPC | 0.01 | 42.17 |
| 17 | 1741 | Non−classical monocyte | 0.88 | 82.37 | Non−classical monocyte | 1.00 | 82.65 | Monocyte | 0.05 | 37.22 | Monocyte | 0.00 | 43.48 |
| 18 | 1445 | CD4+ T | 0.98 | 78.75 | CD4+ T | 0.91 | 81.59 | Lymphocyte progenitor | 0.11 | 40.00 | T | 0.09 | 67.82 |
| 19 | 1403 | CD4+ T | 0.94 | 96.72 | CD4+ T | 0.88 | 99.57 | Granulocyte | 0.01 | 30.15 | T | 0.09 | 69.71 |
| 20 | 1325 | CD8+ T | 0.78 | 78.94 | CD8+ T | 0.78 | 79.77 | Granulocyte | 0.02 | 24.91 | T | 0.08 | 48.75 |
| 21 | 1246 | CD4+ T | 1.00 | 99.68 | CD4+ T | 0.97 | 100.00 | T | 0.01 | 33.39 | T | 0.09 | 72.79 |
| 22 | 1236 | Classical monocyte | 1.00 | 99.76 | Classical monocyte | 0.94 | 97.09 | Monocyte | 0.00 | 29.05 | Monocyte | 0.00 | 41.26 |
| 23 | 798 | Classical monocyte | 0.96 | 85.34 | Classical monocyte | 0.88 | 77.44 | Lymphocyte progenitor | 0.10 | 29.07 | HSPC | 0.00 | 44.61 |
| 24 | 720 | Dendritic | 0.76 | 84.44 | Dendritic | 0.96 | 80.42 | Dendritic | 0.00 | 48.61 | Monocyte | 0.00 | 53.06 |
| 25 | 594 | CD4+ T | 0.97 | 95.62 | CD4+ T | 0.92 | 99.33 | Granulocyte | 0.05 | 34.51 | T | 0.08 | 60.44 |
| 26 | 292 | Megakaryocyte | 0.78 | 39.04 | Megakaryocyte | 0.85 | 41.10 | Granulocyte | 0.07 | 25.34 | T | 0.08 | 32.53 |
| 27 | 213 | pDC | 0.68 | 94.84 | Dendritic | 0.75 | 78.87 | Dendritic | 0.08 | 80.75 | Monocyte | 0.01 | 23.94 |
| 28 | 33 | CD4+ T | 0.97 | 100.00 | CD4+ T | 0.94 | 100.00 | Granulocyte | 0.01 | 42.42 | T | 0.10 | 45.45 |
| 29 | 27 | CD4+ T | 0.96 | 92.59 | CD4+ T | 0.93 | 100.00 | Granulocyte | 0.02 | 37.04 | T | 0.10 | 66.67 |

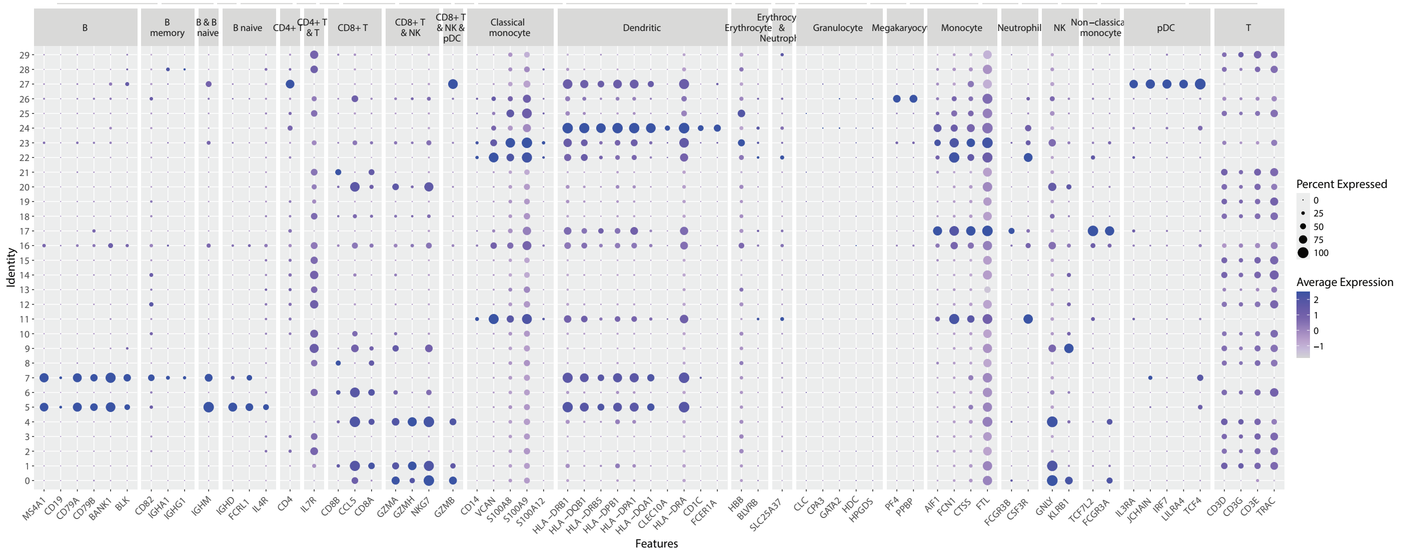

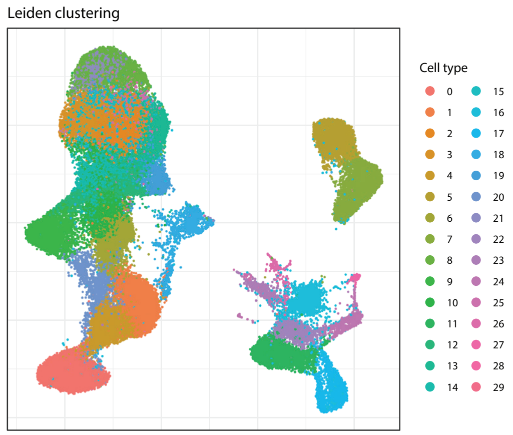

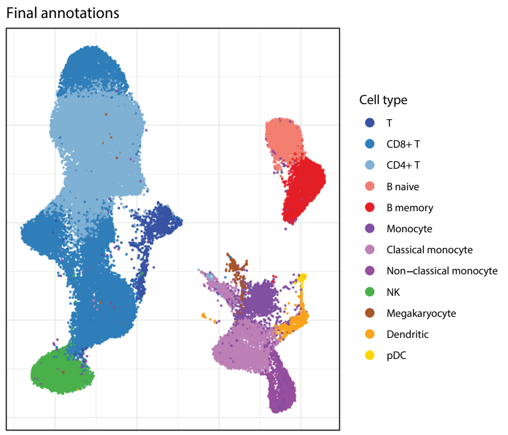

| Cell label | Count | Proportion |
|---|---|---|
| CD4+ T | 21198 | 0.35 |
| CD8+ T | 18287 | 0.30 |
| Classical monocyte | 4550 | 0.07 |
| NK | 4493 | 0.07 |
| B naive | 2995 | 0.05 |
| B memory | 2810 | 0.05 |
| Monocyte | 1999 | 0.03 |
| Non−classical monocyte | 1741 | 0.03 |
| T | 1445 | 0.02 |
| Dendritic | 720 | 0.01 |
| Megakaryocyte | 292 | 0.00 |
| pDC | 213 | 0.00 |

### Fluent V

| | | Seurat | | | | | | SingleR | | | | | |
|---|---|---|---|---|---|---|---|---|---|---|---|---|---|
| | | pbmcsca | | | pbmc3k | | | Mona | | | HPCA | | |
| Cluster | Cell count | Label | Score | % called | Label | Score | % called | Label | Score | % called | Label | Score | % called |
| 0 | 4818 | CD4+ T | 1.00 | 99.96 | CD4+ T | 1.00 | 100.00 | CD4+ T | 0.01 | 46.08 | T | 0.10 | 81.32 |
| 1 | 4688 | CD4+ T | 0.96 | 98.76 | CD4+ T | 0.88 | 99.85 | CD4+ T | 0.06 | 51.49 | T | 0.12 | 90.87 |
| 2 | 4426 | NK | 0.86 | 82.94 | NK | 0.99 | 91.08 | Lymphocyte progenitor | 0.06 | 23.90 | NK | 0.15 | 90.99 |
| 3 | 4387 | CD8+ T | 0.76 | 51.79 | CD4+ T | 0.76 | 53.32 | T | 0.01 | 46.61 | T | 0.10 | 82.95 |
| 4 | 4177 | B | 1.00 | 99.88 | B | 1.00 | 99.76 | B | 0.06 | 41.18 | B | 0.06 | 87.93 |
| 5 | 3597 | CD8+ T | 0.94 | 97.22 | CD8+ T | 0.93 | 97.22 | T | 0.01 | 30.41 | T | 0.10 | 77.20 |
| 6 | 3233 | CD8+ T | 0.86 | 71.48 | CD8+ T | 0.85 | 77.27 | T | 0.01 | 33.44 | T | 0.09 | 67.34 |
| 7 | 3209 | CD4+ T | 1.00 | 99.91 | CD4+ T | 0.93 | 100.00 | CD4+ T | 0.06 | 46.77 | T | 0.12 | 90.06 |
| 8 | 3179 | CD4+ T | 0.89 | 86.06 | CD4+ T | 0.84 | 82.45 | CD4+ T | 0.01 | 41.84 | T | 0.11 | 88.83 |
| 9 | 3124 | B | 1.00 | 99.90 | B | 1.00 | 99.94 | B | 0.06 | 51.09 | B | 0.06 | 90.69 |
| 10 | 2857 | CD4+ T | 1.00 | 99.93 | CD4+ T | 0.94 | 100.00 | CD4+ T | 0.02 | 48.02 | T | 0.11 | 88.20 |
| 11 | 2810 | Non−classical monocyte | 0.90 | 85.02 | Non−classical monocyte | 1.00 | 87.08 | Monocyte | 0.06 | 51.60 | Monocyte | 0.00 | 60.00 |
| 12 | 2762 | CD4+ T | 1.00 | 99.89 | CD4+ T | 0.97 | 100.00 | CD4+ T | 0.02 | 46.20 | T | 0.11 | 87.55 |
| 13 | 2477 | CD4+ T | 1.00 | 99.84 | CD4+ T | 0.97 | 99.96 | T | 0.01 | 57.77 | T | 0.10 | 87.28 |
| 14 | 2458 | CD4+ T | 0.97 | 99.63 | CD4+ T | 0.88 | 99.92 | CD4+ T | 0.06 | 50.28 | T | 0.12 | 89.63 |
| 15 | 2416 | CD4+ T | 1.00 | 99.96 | CD4+ T | 0.96 | 100.00 | CD4+ T | 0.02 | 45.45 | T | 0.12 | 91.06 |
| 16 | 2404 | CD4+ T | 1.00 | 99.83 | CD4+ T | 0.97 | 100.00 | T | 0.01 | 63.81 | T | 0.11 | 90.35 |
| 17 | 2274 | CD8+ T | 0.74 | 54.44 | CD8+ T | 0.92 | 96.13 | NK | 0.00 | 21.68 | T | 0.10 | 79.73 |
| 18 | 2027 | Classical monocyte | 0.99 | 98.22 | Classical monocyte | 0.96 | 98.13 | Monocyte | 0.00 | 27.18 | Monocyte | 0.00 | 39.42 |
| 19 | 1654 | Classical monocyte | 1.00 | 97.88 | Classical monocyte | 0.87 | 88.45 | Granulocyte | 0.10 | 29.02 | Monocyte | 0.00 | 39.18 |
| 20 | 1471 | B | 1.00 | 100.00 | B | 1.00 | 100.00 | B | 0.08 | 52.41 | B | 0.06 | 89.19 |
| 21 | 1399 | CD4+ T | 0.97 | 94.07 | CD4+ T | 0.86 | 95.35 | CD4+ T | 0.06 | 38.24 | T | 0.13 | 83.85 |
| 22 | 1195 | CD4+ T | 0.93 | 96.49 | CD4+ T | 0.94 | 99.67 | CD4+ T | 0.06 | 46.69 | T | 0.11 | 88.45 |
| 23 | 973 | Dendritic | 0.66 | 72.76 | Dendritic | 0.97 | 86.33 | Monocyte | 0.00 | 45.02 | Monocyte | 0.00 | 55.19 |
| 24 | 392 | pDC | 0.59 | 77.04 | Dendritic | 0.85 | 77.30 | Dendritic | 0.11 | 80.36 | Myeloid | 0.00 | 34.44 |
| 25 | 321 | Megakaryocyte | 1.00 | 79.44 | Megakaryocyte | 0.98 | 74.14 | B | 0.23 | 30.53 | Megakaryocyte | 0.08 | 62.93 |
| 26 | 73 | Classical monocyte | 0.53 | 98.63 | CD4+ T | 0.60 | 98.63 | Granulocyte | 0.00 | 36.99 | BM | 0.00 | 56.16 |
| 27 | 72 | Dendritic | 0.60 | 83.33 | Dendritic | 0.95 | 97.22 | Dendritic | 0.07 | 72.22 | Myeloid | 0.00 | 50.00 |
| 28 | 15 | CD4+ T | 0.99 | 86.67 | CD4+ T | 0.70 | 100.00 | B | 0.01 | 26.67 | T | 0.09 | 86.67 |
| 29 | 11 | CD4+ T | 1.00 | 100.00 | CD4+ T | 0.94 | 100.00 | CD4+ T | 0.09 | 36.36 | T | 0.10 | 81.82 |

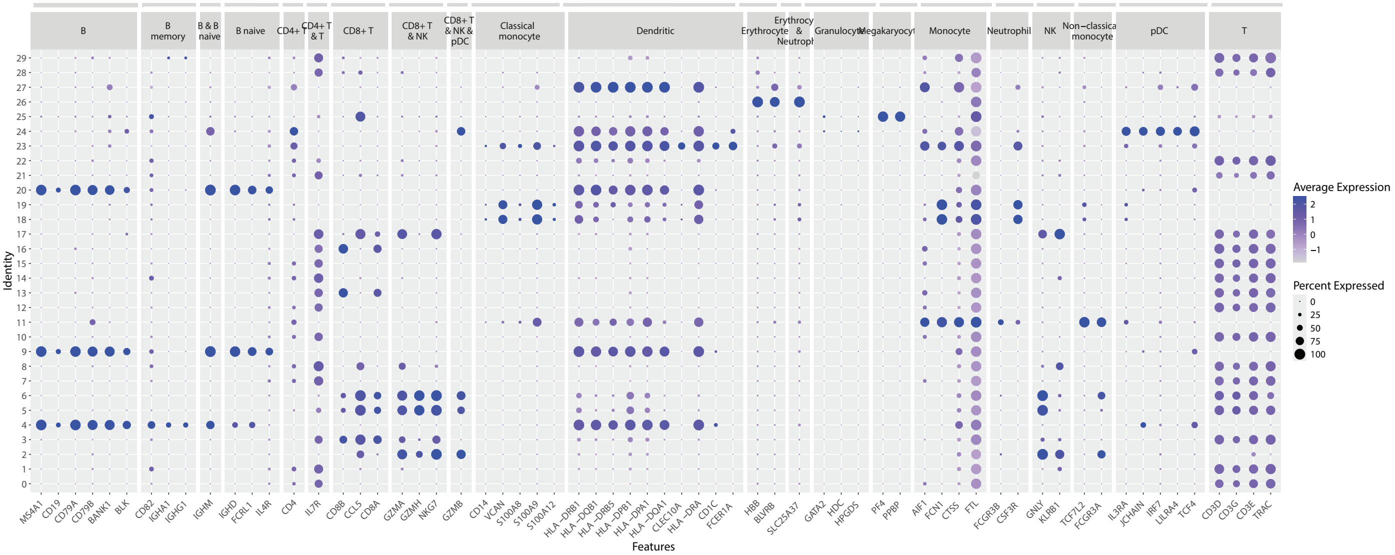

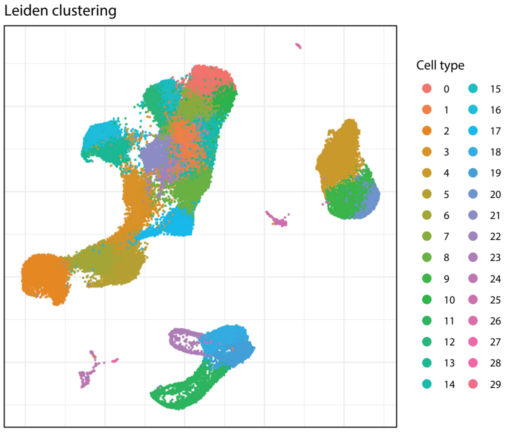

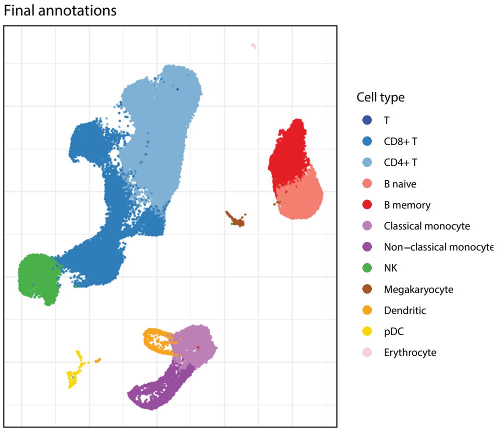

| Cell label | Count | Proportion |
|---|---|---|
| CD4+ T | 28981 | 0.42 |
| CD8+ T | 18387 | 0.27 |
| B naive | 4595 | 0.07 |
| NK | 4426 | 0.06 |
| B memory | 4177 | 0.06 |
| Classical monocyte | 3681 | 0.05 |
| Non−classical monocyte | 2810 | 0.04 |
| Dendritic | 1045 | 0.02 |
| pDC | 392 | 0.01 |
| Megakaryocyte | 321 | 0.00 |
| Erythrocyte | 73 | 0.00 |
| T | 11 | 0.00 |

### Parse v3

| | | Seurat | | | | | | SingleR | | | | | |
|---|---|---|---|---|---|---|---|---|---|---|---|---|---|
| | | pbmcsca | | | pbmc3k | | | Mona | | | HPCA | | |
| Cluster | Cell count | Label | Score | % called | Label | Score | % called | Label | Score | % called | Label | Score | % called |
| 0 | 5059 | CD8+ T | 0.65 | 49.16 | NK | 0.73 | 63.04 | NK | 0.00 | 68.16 | NK | 0.33 | 94.70 |
| 1 | 4794 | CD4+ T | 0.99 | 99.87 | CD4+ T | 0.78 | 100.00 | CD4+ T | 0.07 | 51.44 | T | 0.28 | 96.60 |
| 2 | 4541 | CD4+ T | 0.96 | 98.22 | CD4+ T | 0.75 | 100.00 | CD4+ T | 0.08 | 62.54 | T | 0.26 | 95.49 |
| 3 | 4007 | CD8+ T | 0.73 | 87.87 | CD8+ T | 0.69 | 65.96 | NK | 0.00 | 56.48 | T | 0.10 | 52.98 |
| 4 | 3932 | B | 1.00 | 100.00 | B | 1.00 | 99.97 | B | 0.08 | 83.21 | B | 0.12 | 78.15 |
| 5 | 3910 | CD4+ T | 0.99 | 99.90 | CD4+ T | 0.71 | 100.00 | CD4+ T | 0.08 | 64.02 | T | 0.27 | 96.50 |
| 6 | 3865 | Classical monocyte | 1.00 | 100.00 | Classical monocyte | 0.82 | 95.39 | Monocyte | 0.02 | 54.54 | Granulocyte | 0.13 | 63.78 |
| 7 | 3863 | CD4+ T | 0.96 | 99.77 | CD4+ T | 0.87 | 100.00 | CD4+ T | 0.06 | 52.78 | T | 0.18 | 95.29 |
| 8 | 3766 | B | 1.00 | 99.89 | B | 1.00 | 99.71 | B | 0.08 | 80.72 | B | 0.13 | 73.31 |
| 9 | 3667 | CD8+ T | 0.76 | 87.13 | CD8+ T | 0.66 | 69.40 | NK | 0.00 | 75.24 | NK | 0.21 | 72.46 |
| 10 | 3071 | Classical monocyte | 1.00 | 98.73 | Classical monocyte | 0.75 | 80.56 | Monocyte | 0.02 | 61.77 | Granulocyte | 0.13 | 46.34 |
| 11 | 2885 | CD4+ T | 1.00 | 99.83 | CD4+ T | 0.74 | 100.00 | NK | 0.00 | 40.31 | T | 0.27 | 93.97 |
| 12 | 2808 | CD4+ T | 1.00 | 100.00 | CD4+ T | 0.71 | 100.00 | CD4+ T | 0.07 | 55.63 | T | 0.27 | 97.01 |
| 13 | 2799 | CD4+ T | 0.99 | 99.86 | CD4+ T | 0.78 | 100.00 | NK | 0.00 | 38.01 | T | 0.27 | 95.89 |
| 14 | 2706 | Classical monocyte | 1.00 | 99.93 | Classical monocyte | 0.87 | 94.12 | Monocyte | 0.00 | 45.90 | Granulocyte | 0.13 | 65.00 |
| 15 | 2668 | Non−classical monocyte | 0.71 | 84.07 | Non−classical monocyte | 0.98 | 95.61 | Monocyte | 0.06 | 61.77 | Granulocyte | 0.10 | 45.88 |
| 16 | 2635 | CD4+ T | 0.94 | 97.91 | CD4+ T | 0.78 | 99.58 | CD4+ T | 0.07 | 58.14 | T | 0.17 | 91.65 |
| 17 | 2590 | CD4+ T | 0.71 | 68.57 | CD4+ T | 0.59 | 62.20 | NK | 0.00 | 47.10 | T | 0.13 | 72.86 |
| 18 | 2491 | CD4+ T | 0.86 | 95.78 | CD4+ T | 0.76 | 99.12 | CD4+ T | 0.06 | 48.78 | T | 0.17 | 88.72 |
| 19 | 2454 | CD4+ T | 0.73 | 77.34 | CD4+ T | 0.73 | 89.12 | NK | 0.00 | 45.60 | T | 0.14 | 79.01 |
| 20 | 2137 | CD4+ T | 0.87 | 93.12 | CD4+ T | 0.74 | 98.46 | CD4+ T | 0.07 | 56.76 | T | 0.16 | 89.75 |
| 21 | 2136 | Classical monocyte | 0.94 | 98.08 | Classical monocyte | 0.70 | 75.84 | Monocyte | 0.05 | 66.25 | Monocyte | 0.06 | 50.51 |
| 22 | 1990 | CD4+ T | 0.93 | 98.49 | CD4+ T | 0.76 | 99.65 | CD4+ T | 0.06 | 51.66 | T | 0.17 | 92.46 |
| 23 | 1773 | CD4+ T | 0.86 | 97.69 | CD4+ T | 0.74 | 99.77 | NK | 0.00 | 43.71 | T | 0.16 | 86.35 |
| 24 | 818 | Classical monocyte | 0.52 | 77.87 | Dendritic | 0.86 | 95.11 | Monocyte | 0.00 | 41.93 | Monocyte | 0.07 | 53.79 |
| 25 | 805 | CD4+ T | 0.92 | 97.52 | CD4+ T | 0.72 | 99.75 | NK | 0.00 | 43.60 | T | 0.16 | 83.60 |
| 26 | 615 | CD4+ T | 0.68 | 55.77 | CD8+ T | 0.64 | 51.38 | NK | 0.00 | 69.11 | T | 0.12 | 50.89 |
| 27 | 326 | pDC | 0.69 | 66.87 | Dendritic | 0.86 | 70.55 | Dendritic | 0.09 | 65.95 | NK | 0.01 | 35.58 |
| 28 | 102 | CD4+ T | 0.72 | 82.35 | CD4+ T | 0.71 | 79.41 | B | 0.12 | 48.04 | T | 0.12 | 60.78 |
| 29 | 68 | Classical monocyte | 0.46 | 80.88 | Dendritic | 0.81 | 95.59 | Dendritic | 0.07 | 61.76 | NK | 0.01 | 30.88 |

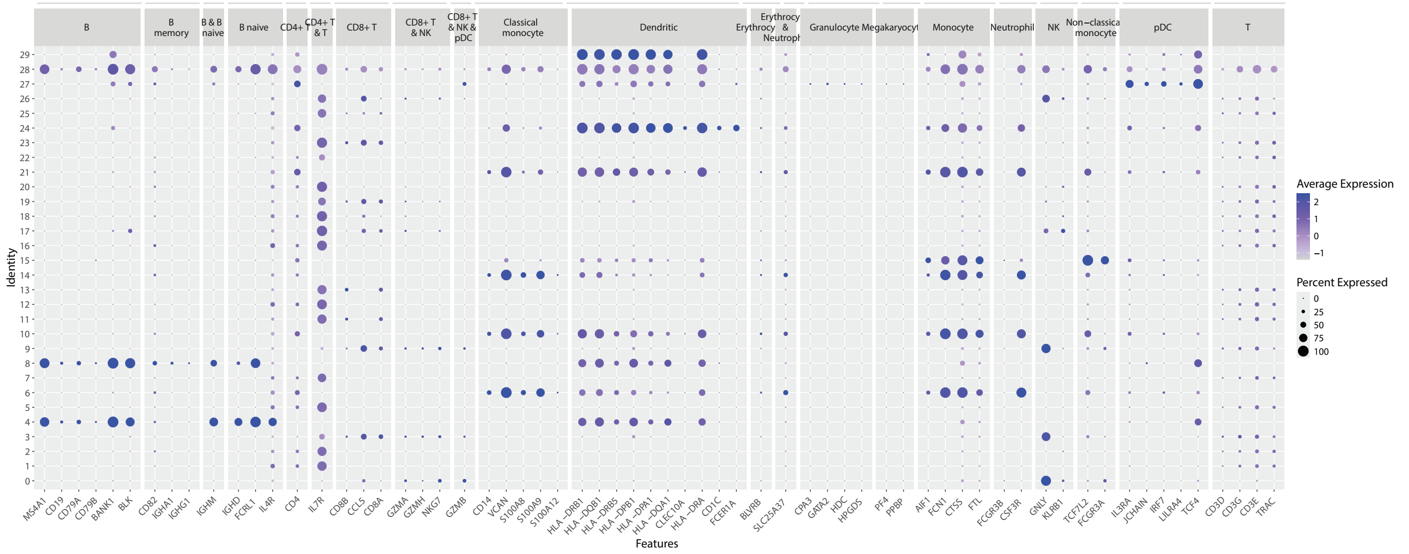

Leiden clustering

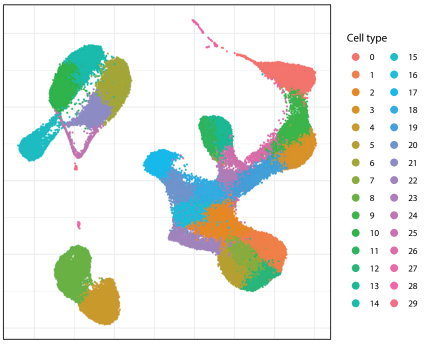

Final annotations

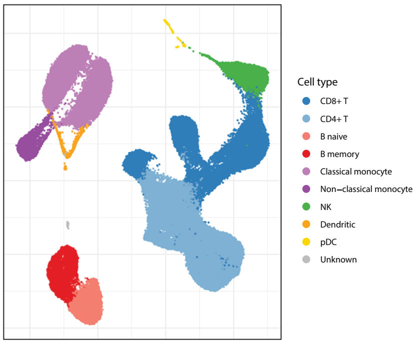

| Cell label | Count | Proportion |
|---|---|---|
| CD4+ T | 29169 | 0.37 |
| CD8+ T | 21595 | 0.27 |
| Classical monocyte | 11778 | 0.15 |
| NK | 5059 | 0.06 |
| B naive | 3932 | 0.05 |
| B memory | 3766 | 0.05 |
| Non−classical monocyte | 2668 | 0.03 |
| Dendritic | 886 | 0.01 |
| pDC | 326 | 0.00 |
| Unknown | 102 | 0.00 |

### Scale

| | | Seurat | | | | | | SingleR | | | | | |
|---|---|---|---|---|---|---|---|---|---|---|---|---|---|
| | | pbmcsca | | | pbmc3k | | | Mona | | | HPCA | | |
| Cluster | Cell count | Label | Score | % called | Label | Score | % called | Label | Score | % called | Label | Score | % called |
| 0 | 7538 | CD4+ T | 1.00 | 99.83 | CD4+ T | 0.73 | 100.00 | CD4+ T | 0.13 | 64.57 | T | 0.17 | 97.61 |
| 1 | 6923 | Classical monocyte | 1.00 | 98.27 | Classical monocyte | 0.87 | 88.75 | Monocyte | 0.00 | 29.89 | Granulocyte | 0.13 | 56.48 |
| 2 | 6626 | Classical monocyte | 1.00 | 99.98 | Classical monocyte | 0.89 | 98.93 | Monocyte | 0.00 | 31.39 | Granulocyte | 0.13 | 69.71 |
| 3 | 6546 | CD8+ T | 0.65 | 51.74 | NK | 0.77 | 69.66 | NK | 0.00 | 39.99 | NK | 0.22 | 93.43 |
| 4 | 6438 | B | 1.00 | 100.00 | B | 1.00 | 99.98 | B | 0.07 | 76.11 | B | 0.09 | 81.30 |
| 5 | 6185 | B | 1.00 | 99.95 | B | 1.00 | 99.92 | B | 0.07 | 68.21 | B | 0.11 | 80.02 |
| 6 | 6045 | CD4+ T | 0.99 | 99.88 | CD4+ T | 0.86 | 99.98 | CD4+ T | 0.11 | 61.99 | T | 0.12 | 95.15 |
| 7 | 5541 | CD8+ T | 0.79 | 91.12 | CD8+ T | 0.73 | 77.22 | NK | 0.00 | 34.69 | T | 0.09 | 53.13 |
| 8 | 5142 | CD4+ T | 0.96 | 98.17 | CD4+ T | 0.80 | 99.73 | CD4+ T | 0.10 | 60.81 | T | 0.14 | 92.34 |
| 9 | 4472 | CD4+ T | 1.00 | 99.96 | CD4+ T | 0.71 | 100.00 | CD8+ T | 0.06 | 61.23 | T | 0.14 | 96.38 |
| 10 | 4460 | Classical monocyte | 0.93 | 96.30 | Classical monocyte | 0.78 | 86.70 | Monocyte | 0.00 | 34.89 | Monocyte | 0.06 | 49.57 |
| 11 | 4420 | CD4+ T | 1.00 | 100.00 | CD4+ T | 0.70 | 100.00 | CD4+ T | 0.14 | 72.35 | T | 0.16 | 97.85 |
| 12 | 4394 | CD4+ T | 0.99 | 99.41 | CD4+ T | 0.75 | 100.00 | CD4+ T | 0.13 | 71.37 | T | 0.18 | 97.66 |
| 13 | 4282 | CD8+ T | 0.80 | 90.99 | CD8+ T | 0.71 | 79.10 | NK | 0.00 | 47.45 | NK | 0.11 | 66.14 |
| 14 | 3836 | CD4+ T | 0.74 | 80.27 | CD4+ T | 0.63 | 56.60 | NK | 0.00 | 27.42 | T | 0.11 | 80.27 |
| 15 | 3717 | CD4+ T | 0.92 | 97.96 | CD4+ T | 0.76 | 99.38 | CD4+ T | 0.07 | 50.07 | T | 0.13 | 89.86 |
| 16 | 3122 | CD4+ T | 0.81 | 77.10 | CD4+ T | 0.71 | 81.42 | CD8+ T | 0.00 | 30.27 | T | 0.11 | 83.79 |
| 17 | 2775 | CD4+ T | 0.92 | 98.13 | CD4+ T | 0.76 | 99.39 | CD8+ T | 0.02 | 48.32 | T | 0.12 | 90.85 |
| 18 | 2672 | Non−classical monocyte | 0.74 | 89.75 | Non−classical monocyte | 1.00 | 93.75 | Monocyte | 0.01 | 44.35 | Granulocyte | 0.10 | 41.62 |
| 19 | 2662 | CD4+ T | 1.00 | 99.89 | CD4+ T | 0.74 | 100.00 | CD8+ T | 0.06 | 54.51 | T | 0.13 | 95.00 |
| 20 | 2657 | CD4+ T | 0.94 | 98.08 | CD4+ T | 0.78 | 99.55 | CD4+ T | 0.09 | 62.85 | T | 0.13 | 92.89 |
| 21 | 2571 | CD4+ T | 0.96 | 98.95 | CD4+ T | 0.79 | 99.65 | CD4+ T | 0.08 | 52.94 | T | 0.11 | 94.83 |
| 22 | 1723 | CD4+ T | 0.99 | 99.59 | CD4+ T | 0.76 | 100.00 | CD4+ T | 0.11 | 68.49 | T | 0.14 | 94.78 |
| 23 | 1415 | Classical monocyte | 0.52 | 61.27 | Dendritic | 0.88 | 93.99 | Dendritic | 0.08 | 35.76 | Monocyte | 0.05 | 59.43 |
| 24 | 1060 | CD4+ T | 0.98 | 98.11 | CD4+ T | 0.71 | 97.36 | NK | 0.00 | 26.51 | T | 0.12 | 88.30 |
| 25 | 823 | Classical monocyte | 0.98 | 77.28 | Classical monocyte | 0.82 | 58.93 | Monocyte | 0.05 | 27.70 | Granulocyte | 0.14 | 48.85 |
| 26 | 621 | CD4+ T | 0.95 | 47.34 | CD4+ T | 0.84 | 49.76 | CD4+ T | 0.10 | 29.31 | T | 0.11 | 62.16 |
| 27 | 471 | CD8+ T | 0.52 | 42.25 | NK | 0.62 | 46.28 | NK | 0.00 | 32.27 | NK | 0.20 | 80.04 |
| 28 | 358 | pDC | 0.65 | 97.49 | Dendritic | 0.77 | 95.25 | Dendritic | 0.09 | 77.09 | Monocyte | 0.00 | 20.11 |
| 29 | 131 | CD4+ T | 0.99 | 99.24 | CD4+ T | 0.73 | 100.00 | CD4+ T | 0.08 | 61.83 | T | 0.13 | 96.18 |

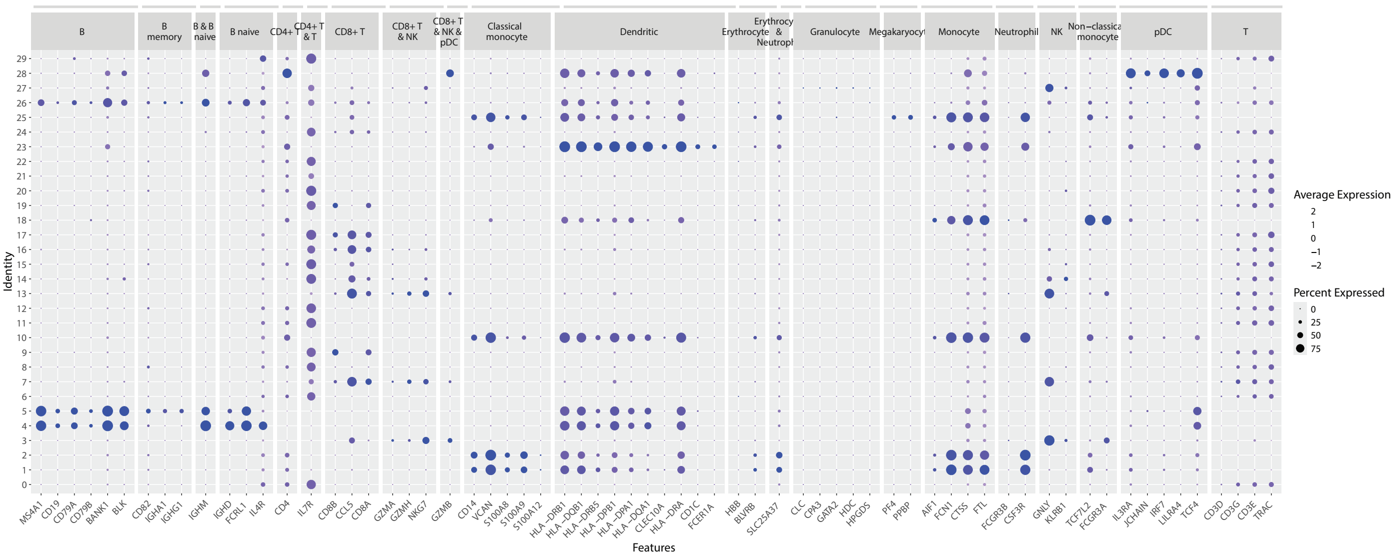

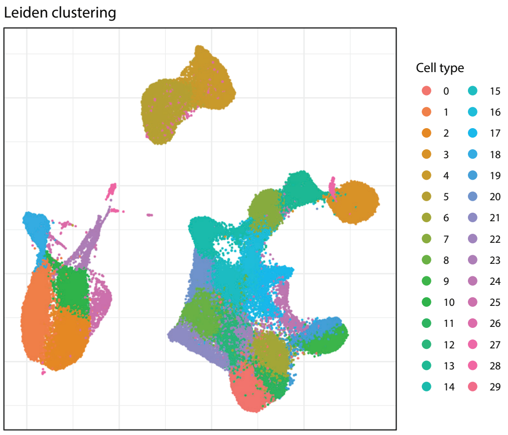

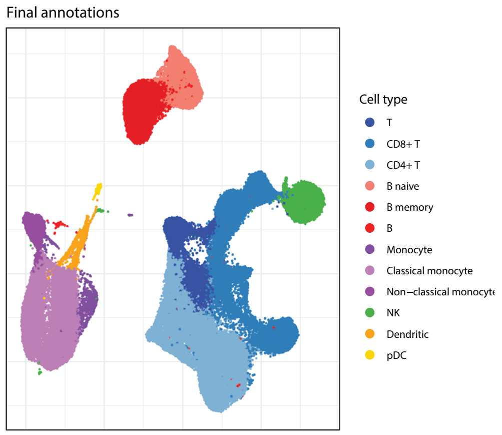

| Cell label | Count | Proportion |
|---|---|---|
| CD4+ T | 34621 | 0.32 |
| CD8+ T | 23914 | 0.22 |
| Classical monocyte | 18009 | 0.16 |
| T | 7553 | 0.07 |
| NK | 7017 | 0.06 |
| B naive | 6438 | 0.06 |
| B memory | 6185 | 0.06 |
| Non−classical monocyte | 2672 | 0.02 |
| Dendritic | 1415 | 0.01 |
| Monocyte | 823 | 0.01 |
| B | 621 | 0.01 |
| pDC | 358 | 0.00 |

### Supplemental figure 5a

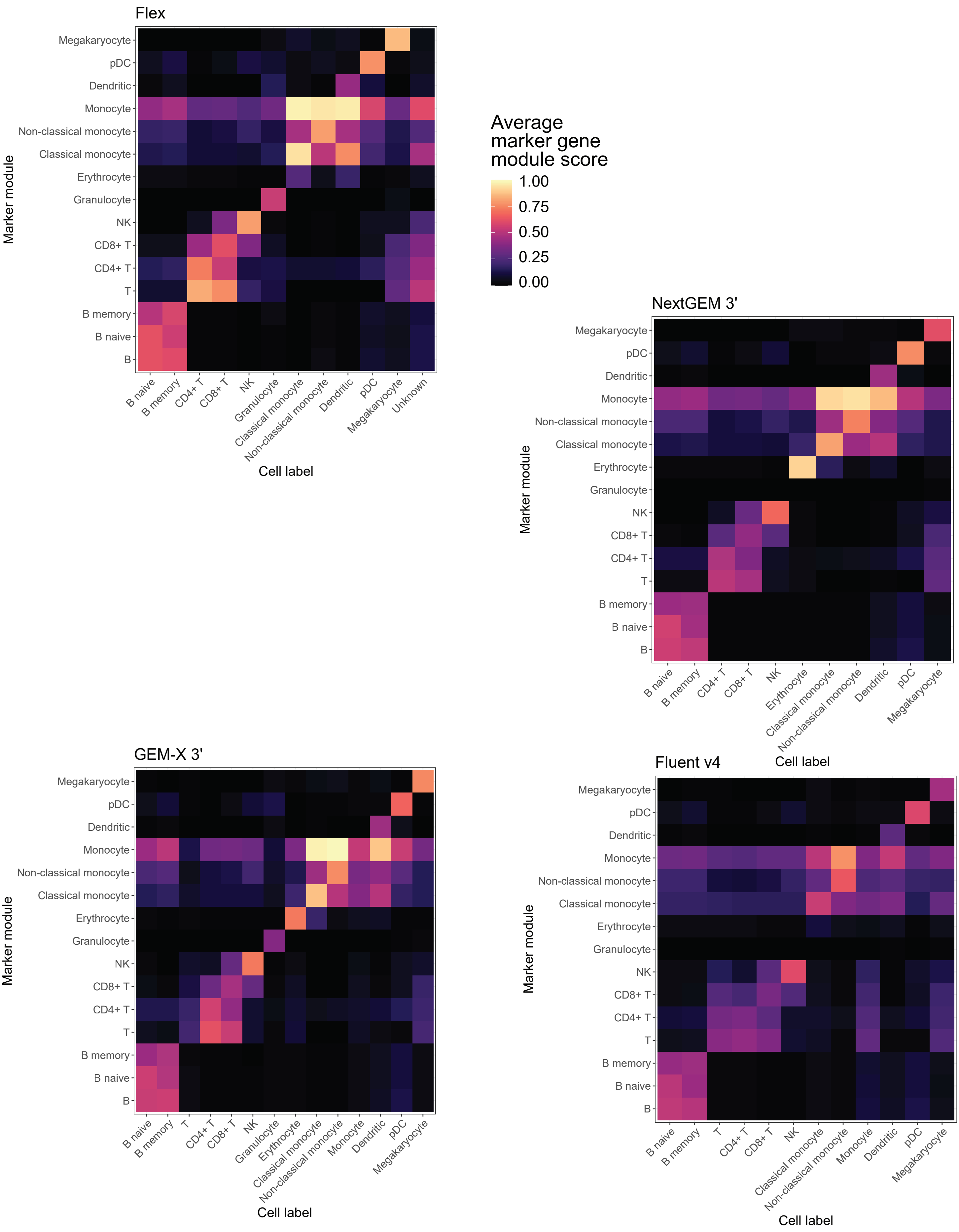

### Supplemental figure 5b

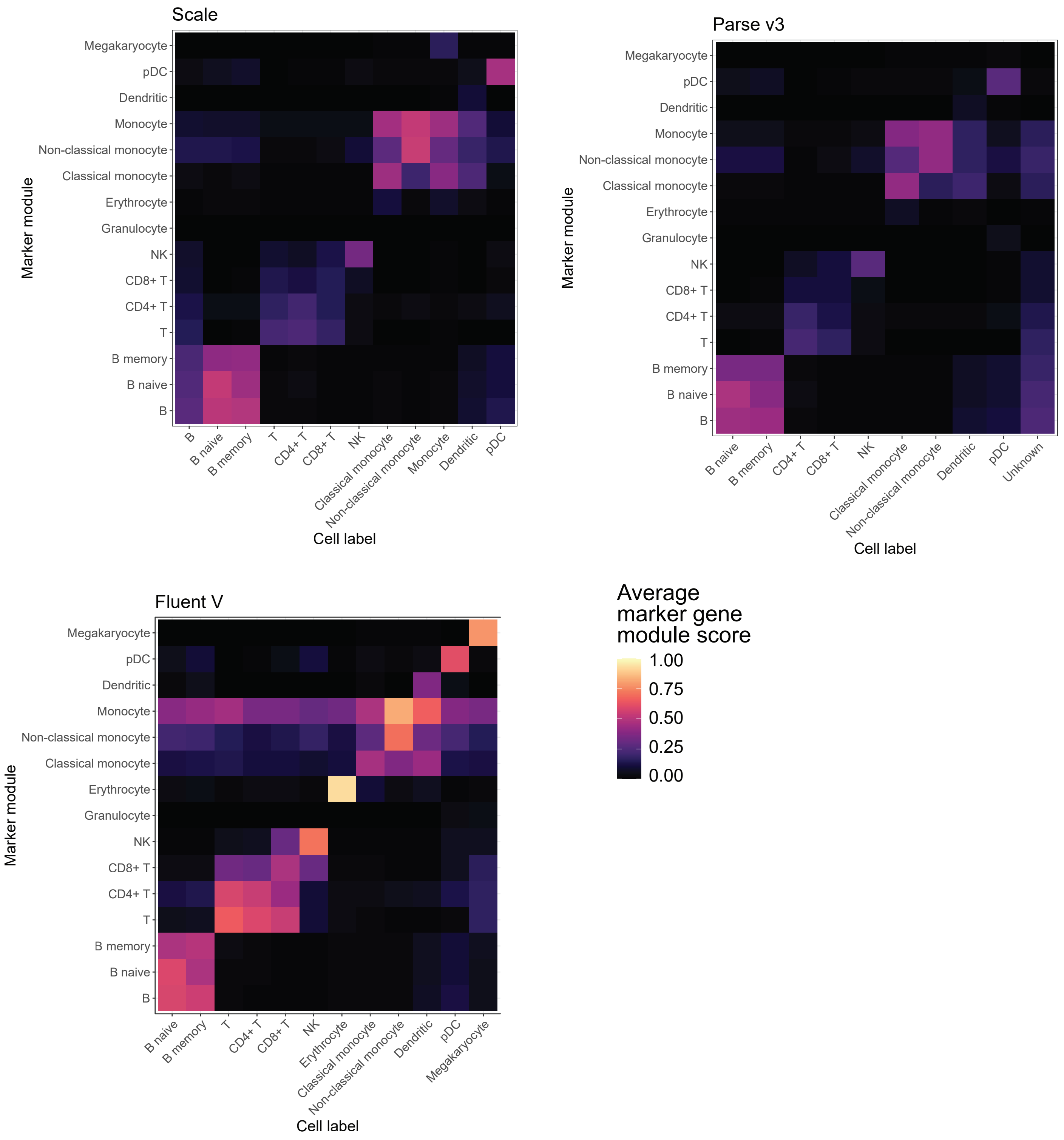

| | | Cell recovery | | | | | |
|---|---|---|---|---|---|---|---|
| | | Figure 2a/b | | | | | Figure 2c |
| | Metric | Recovered cell # | % recovery (of loaded cells) | High quality cell recovery % (of loaded cells) | % multiplets (of recovered cells) | % Low quality cells (of recovered cells) | Cost per recovered cell |
| | Unit | | | | | | |
| | kit level aggregation function | average ± standard error | average ± standard error | average ± standard error | average ± standard error | average ± standard error | |
| Benchmark 1 | Flex | 10195 ± 324 | 60.4% ± 1.9% | 53% ± 1.4% | 10% ± 0.5% | 2.3% ± 0.1% | $0.107 |
| | F1A | 10,024 | 59.4% | 52.8% | 9.0% | 2.1% | |
| | F1B | 10,952 | 64.9% | 56.2% | 11.2% | 2.2% | |
| | F5A | 10,397 | 61.6% | 53.5% | 10.7% | 2.5% | |
| | F5B | 9,408 | 55.8% | 49.3% | 9.3% | 2.3% | |
| | NextGEM | 14808 ± 905 | 62.6% ± 0.3% | 51.7% ± 0.4% | 10.3% ± 0.1% | 7.1% ± 0.3% | $0.230 |
| | F1A | 16,338 | 62.7% | 52.1% | 10.0% | 6.9% | |
| | F1B | 16,403 | 62.9% | 51.6% | 10.1% | 7.9% | |
| | F5A | 13,078 | 61.7% | 50.8% | 10.6% | 7.0% | |
| | F5B | 13,413 | 63.3% | 52.5% | 10.5% | 6.5% | |
| | GEM-X | 34761 ± 1763 | 83.8% ± 1.2% | 71.1% ± 1.5% | 11.6% ± 0.5% | 3.5% ± 0.2% | $0.083 |
| | F1A | 37,147 | 81.0% | 68.1% | 12.0% | 3.9% | |
| | F1B | 38,360 | 83.6% | 69.8% | 12.9% | 3.7% | |
| | F5A | 32,383 | 86.9% | 74.8% | 10.7% | 3.2% | |
| | F5B | 31,155 | 83.6% | 71.9% | 10.9% | 3.1% | |
| | Fluent v4 | 16831 ± 787 | 41.7% ± 1.6% | 37.6% ± 2% | 8.1% ± 0.5% | 1.8% ± 1.5% | $0.060 |
| | F1A | 18,953 | 45.7% | 42.3% | 7.3% | 0.2% | |
| | F1B | 16,574 | 41.4% | 38.1% | 7.8% | 0.3% | |
| | F5A | 16,646 | 41.6% | 37.4% | 9.6% | 0.6% | |
| | F5B | 15,151 | 37.9% | 32.6% | 7.7% | 6.3% | |
| | Fluent V | 18331 ± 318 | 45.9% ± 0.9% | 43.1% ± 1% | 3.6% ± 0.2% | 2.5% ± 0.7% | $0.052 |
| | F1A | 18,709 | 46.5% | 44.1% | 3.7% | 1.4% | |
| | F1B | 17,607 | 43.8% | 41.2% | 3.9% | 2.1% | |
| | F5A | 18,010 | 45.3% | 41.6% | 3.7% | 4.4% | |
| | F5B | 18,997 | 47.8% | 45.4% | 3.1% | 1.9% | |
| | Parse v3 | 22749 ± 1886 | 24.7% ± 3% | 21.5% ± 2.5% | 11.6% ± 0.5% | 1.3% ± 0.2% | $0.145 |
| | F1A | 26,679 | 32.7% | 28.1% | 12.9% | 1.2% | |
| | F1B | 18,961 | 18.9% | 16.4% | 11.8% | 1.7% | |
| | F5A | 25,214 | 25.7% | 22.6% | 10.8% | 1.0% | |
| | F5B | 20,142 | 21.7% | 19.1% | 10.9% | 1.1% | |
| | Scale | 29687 ± 5046 | 23.7% ± 4% | 21.9% ± 4.1% | 8.8% ± 2.9% | 0.2% ± 0.1% | $0.078 |
| | F1A | 15,782 | 12.6% | 10.4% | 17.4% | 0.3% | |
| | F1B | 30,025 | 24.0% | 22.4% | 6.2% | 0.6% | |
| | F5A | 33,298 | 26.6% | 25.2% | 5.4% | 0.0% | |
| | F5B | 39,642 | 31.7% | 29.7% | 6.3% | 0.0% | |
| Benchmark 2 | NextGEM 5' | 8411 ± 928 | 48.9% ± 5.6% | 42.1% ± 4.3% | 9.5% ± 1.9% | 4.1% ± 0.8% | $0.279 |
| | F1 | 7,098 | 41.0% | 36.0% | 6.9% | 5.3% | |
| | F2 | 9,723 | 56.8% | 48.2% | 12.1% | 3.0% | |
| | F4 | | | | | | |
| | F5 | | | | | | |
| | GEM-X 5' | 20406 ± 996 | 68.9% ± 3.5% | 59.1% ± 3.2% | 8.6% ± 0.3% | 5.6% ± 0.7% | $0.099 |
| | F1 | 18,103 | 59.5% | 50.5% | 8.0% | 7.1% | |
| | F2 | 22,708 | 75.7% | 65.5% | 9.2% | 4.1% | |
| | F4 | 21,211 | 72.6% | 61.3% | 9.0% | 6.5% | |
| | F5 | 19,601 | 67.8% | 59.1% | 8.1% | 4.8% | |
| | Parse v2 | 22825 ± 418 | 24.6% ± 0.8% | 23.2% ± 0.6% | 4.9% ± 0.6% | 0.7% ± 0.1% | $0.135 |
| | F1 | 23,921 | 26.9% | 25.0% | 6.1% | 1.0% | |
| | F2 | 23,022 | 23.3% | 22.0% | 5.6% | 0.4% | |
| | F4 | 22,285 | 24.0% | 22.9% | 3.9% | 0.7% | |
| | F5 | 22,072 | 24.2% | 23.1% | 4.1% | 0.7% | |

| | | | Sequencing | | | | | | |
|---|---|---|---|---|---|---|---|---|---|
| | | | Supplemental Figure 2 a-b | | Supplemental Figure 2 c-d | Figure 3a/c | | Figure 3b/d | |
| | Metric | | % of reads mapped to transcriptome | % of reads mapped and in called cells | % Ambient RNA | Maximum gene estimate | rd50 gene | Maximum UMI estimate | rd50 UMI |
| | Unit | | | | | | Reads per cell | | Reads per cell |
| | kit level aggregation function | | average ± standard error | average ± standard error | average ± standard error | | | | |
| Benchmark 1 | Flex | | 94.3% ± 0.1% | 88.9% ± 0% | 1.2% ± 0.1% | 2,993 | 4,853 | 6,229 | 8,971 |
| | | F1A | 94.47% | 88.90% | 1.20% | | | | |
| | | F1B | 94.11% | 88.95% | 1.30% | | | | |
| | | F5A | 94.27% | 88.90% | 1.40% | | | | |
| | | F5B | 94.49% | 88.97% | 1.00% | | | | |
| | NextGEM | | 72.5% ± 0.3% | 68.2% ± 0.3% | 1.1% ± 0.1% | 2,710 | 7,333 | 9,032 | 14,293 |
| | | F1A | 72.18% | 68.10% | 1.10% | | | | |
| | | F1B | 72.28% | 67.89% | 1.30% | | | | |
| | | F5A | 73.43% | 69.05% | 1.00% | | | | |
| | | F5B | 72.14% | 67.87% | 0.90% | | | | |
| | GEM-X | | 64.7% ± 0.5% | 59.8% ± 0.6% | 2.4% ± 0% | 3,923 | 10,895 | 17,834 | 32,415 |
| | | F1A | 66.12% | 61.35% | 2.30% | | | | |
| | | F1B | 63.97% | 59.39% | 2.50% | | | | |
| | | F5A | 64.67% | 59.54% | 2.40% | | | | |
| | | F5B | 63.86% | 58.76% | 2.30% | | | | |
| | Fluent v4 | | 77.4% ± 0.8% | 37.1% ± 0.7% | 8.3% ± 0.5% | 1,479 | 5,644 | 3,146 | 10,233 |
| | | F1A | 79.63% | 37.82% | 9.20% | | | | |
| | | F1B | 77.52% | 34.98% | 9.00% | | | | |
| | | F5A | 75.92% | 38.24% | 7.60% | | | | |
| | | F5B | 76.63% | 37.22% | 7.40% | | | | |
| | Fluent V | | 73.2% ± 2.9% | 58.5% ± 3.3% | 1.8% ± 0.2% | 1,721 | 5,679 | 4,882 | 9,114 |
| | | F1A | 66.76% | 51.42% | 2.10% | | | | |
| | | F1B | 71.30% | 55.18% | 1.90% | | | | |
| | | F5A | 80.48% | 66.11% | 1.30% | | | | |
| | | F5B | 74.41% | 61.48% | 1.70% | | | | |
| | Parse v3 | | 38.1% ± 1.1% | 33.8% ± 1% | 1.8% ± 0.1% | 2,563 | 10,732 | 6,857 | 24,452 |
| | | F1A | 35.84% | 31.98% | 1.90% | | | | |
| | | F1B | 37.40% | 32.53% | 1.80% | | | | |
| | | F5A | 40.88% | 36.63% | 1.70% | | | | |
| | | F5B | 38.31% | 34.04% | 1.60% | | | | |
| | Scale | | 63.3% ± 1.4% | 57.3% ± 2.7% | 1.9% ± 0.3% | 2,497 | 4,798 | 5,140 | 9,058 |
| | | F1A | 60.15% | 49.94% | 2.70% | | | | |
| | | F1B | 62.70% | 57.61% | 1.40% | | | | |
| | | F5A | 66.70% | 62.60% | 1.40% | | | | |
| | | F5B | 63.70% | 59.19% | 1.90% | | | | |
| Benchmark 2 | NextGEM 5' | | 70.8% ± 0% | 68.7% ± 0.2% | 0.5% ± 0% | 2,213 | 5,771 | 7,197 | 9,647 |
| | | F1 | 70.89% | 68.47% | 0.50% | | | | |
| | | F2 | 70.76% | 68.90% | 0.50% | | | | |
| | | F4 | | | | | | | |
| | | F5 | | | | | | | |
| | GEM-X 5' | | 62.8% ± 1.4% | 59.9% ± 1.6% | 0.6% ± 0.1% | 3,484 | 10,471 | 16,391 | 26,590 |
| | | F1 | 64.52% | 61.28% | 0.50% | | | | |
| | | F2 | 65.76% | 63.74% | 0.40% | | | | |
| | | F4 | 59.73% | 56.32% | 0.90% | | | | |
| | | F5 | 61.02% | 58.23% | 0.50% | | | | |
| | Parse v2 | | 54.3% ± 0.7% | 49.4% ± 0.9% | 1.8% ± 0.1% | 2,483 | 6,324 | 6,876 | 14,396 |
| | | F1 | 53.10% | 48.31% | 1.50% | | | | |
| | | F2 | 53.27% | 47.57% | 2.00% | | | | |
| | | F4 | 55.74% | 51.33% | 1.90% | | | | |
| | | F5 | 55.27% | 50.29% | 1.80% | | | | |

| | Metric | | Median genes per cell | Median UMI per cell | Median ribosomal read percentage per cell | Median mitochondrial read percentage per cell | Unique gene count | % of reads assigned to 'informative' genes (not ribosomal, mitochondrial, lncRNA, unidentified etc.) | Percentage of reads from unique genes |
|---|---|---|---|---|---|---|---|---|---|
| | | | Sensitivity | | | | | | |
| | | | Figure 4a | | | | Figure 4b | Figure 4c | |
| | Unit | | | | | | | | |
| | kit level aggregation function | | average ± standard error | average ± standard error | average ± standard error | average ± standard error | | average ± standard error | |
| Benchmark 1 | Flex | | 2397 ± 32 | 4282 ± 70 | 0% ± 0% | 1% ± 0% | 163 | 95.7% ± 0% | 2% ± 0% |
| | | F1A | 2342 | 4165 | 0.0% | 1.0% | | 95.6% | 2.0% |
| | | F1B | 2341 | 4157 | 0.0% | 1.1% | | 95.6% | 2.0% |
| | | F5A | 2457 | 4412 | 0.0% | 1.0% | | 95.7% | 1.9% |
| | | F5B | 2449 | 4395 | 0.0% | 0.9% | | 95.7% | 1.9% |
| | NextGEM | | 2080 ± 32 | 5628 ± 127 | 26.9% ± 0.5% | 5.9% ± 0.1% | 62 | 81.5% ± 0.1% | 4.4% ± 0% |
| | | F1A | 2163 | 5976 | 27.0% | 5.7% | | 81.6% | 4.4% |
| | | F1B | 2008 | 5470 | 28.2% | 5.8% | | 81.2% | 4.4% |
| | | F5A | 2085 | 5655 | 26.8% | 6.2% | | 81.6% | 4.4% |
| | | F5B | 2064 | 5412 | 25.7% | 6.1% | | 81.6% | 4.2% |
| | GEM-X | | 2704 ± 21 | 7560 ± 92 | 20.6% ± 0.1% | 3.2% ± 0% | 693 | 84.7% ± 0% | 3.4% ± 0% |
| | | F1A | 2730 | 7779 | 20.7% | 3.2% | | 84.7% | 3.5% |
| | | F1B | 2643 | 7357 | 20.8% | 3.2% | | 84.7% | 3.4% |
| | | F5A | 2731 | 7630 | 20.5% | 3.3% | | 84.8% | 3.4% |
| | | F5B | 2713 | 7472 | 20.4% | 3.3% | | 84.8% | 3.4% |
| | Fluent v4 | | 1159 ± 42 | 2087 ± 66 | 22.4% ± 0.9% | 4.4% ± 0.1% | 90 | 80.1% ± 0.3% | 4.6% ± 0% |
| | | F1A | 1188 | 2139 | 23.0% | 4.2% | | 80.1% | 4.6% |
| | | F1B | 1190 | 2120 | 22.1% | 4.2% | | 80.4% | 4.6% |
| | | F5A | 1223 | 2193 | 20.2% | 4.7% | | 80.7% | 4.6% |
| | | F5B | 1036 | 1894 | 24.4% | 4.4% | | 79.2% | 4.6% |
| | Fluent V | | 1433 ± 43 | 3616 ± 121 | 34% ± 0.8% | 5.6% ± 0.1% | 530 | 78.3% ± 0.3% | 4.7% ± 0.1% |
| | | F1A | 1359 | 3374 | 34.0% | 5.6% | | 78.1% | 4.9% |
| | | F1B | 1397 | 3482 | 34.4% | 5.7% | | 78.1% | 4.8% |
| | | F5A | 1418 | 3681 | 35.8% | 5.4% | | 77.8% | 4.5% |
| | | F5B | 1557 | 3925 | 31.8% | 5.6% | | 79.2% | 4.7% |
| | Parse v3 | | 1761 ± 41 | 3365 ± 136 | 0.7% ± 0% | 4.4% ± 0.2% | 2162 | 90.7% ± 0.1% | 4.8% ± 0.1% |
| | | F1A | 1753 | 3306 | 0.7% | 4.2% | | 90.8% | 4.7% |
| | | F1B | 1670 | 3067 | 0.7% | 4.1% | | 90.8% | 4.6% |
| | | F5A | 1754 | 3364 | 0.6% | 5.0% | | 90.6% | 4.8% |
| | | F5B | 1868 | 3724 | 0.6% | 4.4% | | 90.5% | 4.9% |
| | Scale | | 2037 ± 106 | 3585 ± 321 | 1% ± 0.1% | 0.6% ± 0% | 10946 | 87.4% ± 0.3% | 7.1% ± 0.1% |
| | | F1A | 1834 | 3000 | 1.1% | 0.5% | | 87.1% | 7.2% |
| | | F1B | 1931 | 3256 | 1.3% | 0.6% | | 86.9% | 7.2% |
| | | F5A | 2324 | 4469 | 0.8% | 0.7% | | 88.1% | 6.6% |
| | | F5B | 2058 | 3613 | 0.9% | 0.6% | | 87.3% | 7.1% |
| Benchmark 2 | NextGEM 5' | | 1784 ± 19 | 5072 ± 63 | 35.7% ± 1.4% | 3.2% ± 0.2% | | 81.1% ± 0.3% | 0.7% ± 0% |
| | | F1 | 1810 | 4982 | 33.7% | 3.5% | | 81.5% | 0.7% |
| | | F2 | 1757 | 5161 | 37.6% | 3.0% | | 80.7% | 0.8% |
| | | F4 | 668 | 1598 | 43.3% | 2.9% | | | |
| | | F5 | 850 | 2266 | 43.7% | 1.8% | | | |
| | GEM-X 5' | | 2478 ± 21 | 7942 ± 85 | 31.7% ± 0.8% | 4.1% ± 0.3% | | 82.7% ± 0.2% | 0.8% ± 0% |
| | | F1 | 2499 | 8106 | 31.1% | 4.3% | | 82.8% | 0.7% |
| | | F2 | 2419 | 8014 | 33.4% | 4.0% | | 82.3% | 0.8% |
| | | F4 | 2517 | 7707 | 29.9% | 4.6% | | 83.0% | 0.9% |
| | | F5 | 2477 | 7940 | 32.5% | 3.4% | | 82.9% | 0.8% |
| | Parse v2 | | 1943 ± 31 | 4176 ± 107 | 1.1% ± 0.1% | 3.8% ± 0.2% | | 90% ± 0.1% | 2.9% ± 0.1% |
| | | F1 | 1953 | 4167 | 1.0% | 3.6% | | 90.2% | 2.8% |
| | | F2 | 2022 | 4452 | 1.1% | 3.8% | | 90.0% | 2.9% |
| | | F4 | 1876 | 3930 | 1.2% | 3.6% | | 89.6% | 3.1% |
| | | F5 | 1920 | 4156 | 1.3% | 4.4% | | 90.2% | 2.8% |

| | | | Biological resolution | |
|---|---|---|---|---|
| | | Figure 6a | Figure 6b | Figure 6c |
| | Metric | Earth mover's distance | Percent of residual variance per gene | Successfully detected enriched pathways |
| | Unit | mean ± CI | median (IQR: Q1-Q3) | Number correctly reported of 6 expected |
| | kit level aggregation function | | | |
| Benchmark 1 | Flex | | 9 (3-26) % | 4/6 |
| | F1A | 0.06 ± 0.01 | | |
| | F1B | | | |
| | F5A | 0.07 ± 0.01 | | |
| | F5B | | | |
| | NextGEM | | 15 (5-38) % | 2/6 |
| | F1A | 0.28 ± 0.08 | | |
| | F1B | | | |
| | F5A | 0.14 ± 0.03 | | |
| | F5B | | | |
| | GEM-X | | 12 (4-33) % | 3/6 |
| | F1A | 0.21 ± 0.03 | | |
| | F1B | | | |
| | F5A | 0.12 ± 0.03 | | |
| | F5B | | | |
| | Fluent v4 | | 44 (17-81) % | 2/6 |
| | F1A | 0.34 ± 0.1 | | |
| | F1B | | | |
| | F5A | 1.38 ± 0.21 | | |
| | F5B | | | |
| | Fluent V | | 26 (9-60) % | 3/6 |
| | F1A | 0.17 ± 0.03 | | |
| | F1B | | | |
| | F5A | 0.35 ± 0.06 | | |
| | F5B | | | |
| | Parse v3 | | 24 (8-57) % | 1/6 |
| | F1A | 0.29 ± 0.05 | | |
| | F1B | | | |
| | F5A | 0.17 ± 0.04 | | |
| | F5B | | | |
| | Scale | | 39 (14-77) % | 1/6 |
| | F1A | 0.52 ± 0.1 | | |
| | F1B | | | |
| | F5A | 0.29 ± 0.04 | | |
| | F5B | | | |
| Benchmark 2 | NextGEM 5' | | | |
| | F1 | | | |
| | F2 | | | |
| | F4 | | | |
| | F5 | | | |
| | GEM-X 5' | | | |
| | F1 | | | |
| | F2 | | | |
| | F4 | | | |
| | F5 | | | |
| | Parse v2 | | | |
| | F1 | | | |
| | F2 | | | |
| | F4 | | | |
| | F5 | | | |

| | | TCR recovery | | | | | | |
|---|---|---|---|---|---|---|---|---|
| | | Figure 2d | | | | | Figure 2b | Figure 2f |
| | Metric | Cost per unique clonotype (1+TRA 1+TRB paired clonotypes only) | Cost per unique clonotype (including partial) | Cost per recovered clone (1+TRA 1+TRB paired clones only) | Cost per recovered clone (including partial) | Tcells recovered with clean TRA;TRB clones | Proportion of loaded cells recovered with clean TRA;TRB clones | Inverse Simpson index |
| | Unit | | | | | | | |
| | kit level aggregation function | | | | | average ± standard error | average ± standard error | |
| Benchmark 1 | Flex | | | | | | | |
| | F1A | | | | | | | |
| | F1B | | | | | | | |
| | F5A | | | | | | | |
| | F5B | | | | | | | |
| | NextGEM | | | | | | | |
| | F1A | | | | | | | |
| | F1B | | | | | | | |
| | F5A | | | | | | | |
| | F5B | | | | | | | |
| | GEM-X | | | | | | | |
| | F1A | | | | | | | |
| | F1B | | | | | | | |
| | F5A | | | | | | | |
| | F5B | | | | | | | |
| | Fluent v4 | | | | | | | |
| | F1A | | | | | | | |
| | F1B | | | | | | | |
| | F5A | | | | | | | |
| | F5B | | | | | | | |
| | Fluent V | | | | | | | |
| | F1A | | | | | | | |
| | F1B | | | | | | | |
| | F5A | | | | | | | |
| | F5B | | | | | | | |
| | Parse v3 | | | | | | | |
| | F1A | | | | | | | |
| | F1B | | | | | | | |
| | F5A | | | | | | | |
| | F5B | | | | | | | |
| | Scale | | | | | | | |
| | F1A | | | | | | | |
| | F1B | | | | | | | |
| | F5A | | | | | | | |
| | F5B | | | | | | | |
| Benchmark 2 | NextGEM 5' | $0.39 | $0.32 | $0.36 | $0.29 | 5765 ± 860 | 33.5% ± 5.1% | 2760 ± |
| | F1 | | | | | 4,548 | 26.28% | 1042 |
| | F2 | | | | | 6,981 | 40.80% | 4479 |
| | F4 | | | | | | | |
| | F5 | | | | | | | |
| | GEM-X 5' | $0.14 | $0.12 | $0.12 | $0.10 | 14847 ± 1239 | 50.1% ± 4.2% | |
| | F1 | | | | | 12,313 | 40.49% | 973 |
| | F2 | | | | | 18,251 | 60.81% | 7056 |
| | F4 | | | | | 14,565 | 49.86% | 383 |
| | F5 | | | | | 14,258 | 49.31% | 601 |
| | Parse v2 | $0.46 | $0.20 | $0.40 | $0.18 | 7804 ± 445 | 8.4% ± 0.3% | |
| | F1 | | | | | 6,672 | 7.49% | 837 |
| | F2 | | | | | 8,837 | 8.96% | 4884 |
| | F4 | | | | | 7,972 | 8.60% | 624 |
| | F5 | | | | | 7,734 | 8.49% | 471 |

#### Supplemental table 3: Modeled sequencing cost

Projected sequencing costs for each kit normalized for equivalent data recovery. Maximum UMI recovery and RD50 are calculated from a Michaelis-Menten model fit to the observed sequencing saturation data. Reads per cell required to recover a median of 2000 UMI per cell is calculated based on these values. Sequencing price is set at $1.5e-06 per read.

| Kit Abbreviation | Projected maximum UMI recovery | rd50 | Reads per cell required for 2000 UMI | Normalized sequencing cost (80k cells) | Expected cell recovery | Sequencing cost of whole kit |
|---|---|---|---|---|---|---|
| Flex | 6229 | 8971 | 4243 | $ 509 | 142956 | $ 910 |
| NextGEM3P | 9032 | 14293 | 4065 | $ 488 | 34154 | $ 208 |
| GEMX3P | 17834 | 32415 | 4094 | $ 491 | 82221 | $ 505 |
| Fluent_v4 | 3146 | 10233 | 17865 | $ 2,144 | 60201 | $ 1,613 |
| Fluent_V | 4882 | 9114 | 6325 | $ 759 | 68933 | $ 654 |
| Parse_v3 | 6857 | 24452 | 10069 | $ 1,208 | 74281 | $ 1,122 |
| Scale | 5140 | 9058 | 5769 | $ 692 | 109626 | $ 949 |
| NextGEM5P | 7197 | 9647 | 3712 | $ 445 | 28070 | $ 156 |
| GEMX5P | 16391 | 26590 | 3695 | $ 443 | 68486 | $ 380 |
| Parse_v2 | 6876 | 14396 | 5905 | $ 709 | 92754 | $ 822 |

#### Marker genes used for celltype modules and annotation

| gene_symbol | expression_level | celltype | relative_to | references |
|---|---|---|---|---|
| MS4A1 | Increased | B | Leukocyte | https://doi.org/10.3390/ijms25073828; https://doi.org/10.3389/fimmu.2023.1223471; https://doi.org/10.1038/s41587-020-0465-8 |
| CD19 | Increased | B | Leukocyte | https://doi.org/10.1002/eji.201747168; https://doi.org/10.1038/s41587-020-0465-8; https://doi.org/10.3389/fimmu.2023.1223471 |
| CD79A | Increased | B | Leukocyte | https://doi.org/10.3389/fimmu.2023.1223471; https://doi.org/10.1038/s41587-020-0465-8 |
| CD79B | Increased | B | Leukocyte | https://doi.org/10.3389/fimmu.2023.1223471; https://doi.org/10.1038/s41587-020-0465-8 |
| AFF3 | Increased | B | Leukocyte | https://doi.org/10.3389/fimmu.2023.1223471 |
| BANK1 | Increased | B | Leukocyte | https://doi.org/10.3389/fimmu.2023.1223471 |
| BLK | Increased | B | Leukocyte | https://doi.org/10.3389/fimmu.2023.1223471 |
| BTLA | Increased | B | Leukocyte | https://doi.org/10.3389/fimmu.2023.1223471 |
| IGHM | Increased | B | Leukocyte | https://doi.org/10.3389/fimmu.2023.1223471 |
| IGHG3 | Increased | B | Leukocyte | https://doi.org/10.3389/fimmu.2023.1223471 |
| IL7R | Increased | CD4+ T | Leukocyte | https://doi.org/10.1038/s41587-020-0465-8 |
| CD4 | Increased | CD4+ T | Leukocyte | https://doi.org/10.3390/ijms25073828 |
| CD27 | Increased | CD4+ T | Leukocyte | https://doi.org/10.1038/s41587-020-0465-8 |
| IL7R | Increased | CD4+ T | CD8+ T | cellxgene |
| CD8A | Increased | CD8+ T | Leukocyte | https://doi.org/10.3390/ijms25073828 |
| GZMA | Increased | CD8+ T | Leukocyte | https://www.sc-best-practices.org/cellular_structure/annotation.html |
| GZMB | Increased | CD8+ T | Leukocyte | https://www.sc-best-practices.org/cellular_structure/annotation.html |
| GZMH | Increased | CD8+ T | Leukocyte | https://www.sc-best-practices.org/cellular_structure/annotation.html |
| CD8A | Increased | CD8+ T | Leukocyte | https://doi.org/10.1038/s41587-020-0465-8 |
| CD8B | Increased | CD8+ T | Leukocyte | https://doi.org/10.1038/s41587-020-0465-8 |
| CCL5 | Increased | CD8+ T | Leukocyte | https://doi.org/10.1038/s41587-020-0465-8 |
| NKG7 | Increased | CD8+ T | Leukocyte | https://doi.org/10.1038/s41587-020-0465-8 |
| CD14 | Increased | Classical monocyte | Leukocyte | https://doi.org/10.3390/ijms25073828; https://doi.org/10.1038/s41587-020-0465-8 |
| VCAN | Increased | Classical monocyte | Leukocyte | https://doi.org/10.1038/s41587-020-0465-8 |
| S100A8 | Increased | Classical monocyte | Leukocyte | https://doi.org/10.1038/s41587-020-0465-8 |
| S100A9 | Increased | Classical monocyte | Leukocyte | https://doi.org/10.1038/s41587-020-0465-8 |
| CSF3R | Increased | Classical monocyte | Leukocyte | https://doi.org/10.1038/s41587-020-0465-8 |
| LYZ | Increased | Classical monocyte | Non-classical | https://doi.org/10.1038/s41587-020-0465-8 |
| S100A12 | Increased | Classical monocyte | Leukocyte | https://doi.org/10.1038/s41587-020-0465-8 |
| HLA-DRB5 | Increased | Classical monocyte | Myeloid | cellxgene |
| FCER1A | Increased | Dendritic | Leukocyte | https://scanpy.readthedocs.io/en/stable/tutorials/basics/clustering-2017.html#finding-marker-genes |
| CD1C | Increased | Dendritic | Leukocyte | https://doi.org/10.3389/fimmu.2023.1223471 |
| HLA-DRB1 | Increased | Dendritic | Leukocyte | https://doi.org/10.3389/fimmu.2023.1223471 |
| HLA-DQB1 | Increased | Dendritic | Leukocyte | https://doi.org/10.3389/fimmu.2023.1223471 |
| HLA-DRB5 | Increased | Dendritic | Leukocyte | https://doi.org/10.3389/fimmu.2023.1223471 |
| FCER1A | Increased | Dendritic | Leukocyte | https://doi.org/10.3389/fimmu.2023.1223471 |
| HLA-DPB1 | Increased | Dendritic | Leukocyte | https://doi.org/10.1038/s41587-020-0465-8; https://doi.org/10.3389/fimmu.2023.1223471 |
| HLA-DPA1 | Increased | Dendritic | Myeloid | https://doi.org/10.1038/s41587-020-0465-8; https://doi.org/10.3389/fimmu.2023.1223471 |
| HLA-DQA1 | Increased | Dendritic | Myeloid | https://doi.org/10.1038/s41587-020-0465-8; https://doi.org/10.3389/fimmu.2023.1223471 |
| CD1C | Increased | Dendritic | Leukocyte | https://doi.org/10.1038/s41587-020-0465-8 |
| FCER1A | Increased | Dendritic | Leukocyte | https://doi.org/10.1038/s41587-020-0465-8 |
| CLEC10A | Increased | Dendritic | Leukocyte | https://doi.org/10.1038/s41587-020-0465-8 |
| HLA-DRA | Increased | Dendritic | Myeloid | cellxgene |
| PF4 | Increased | Megakaryocyte | Leukocyte | https://doi.org/10.1038/s41587-020-0465-8 |
| PPBP | Increased | Megakaryocyte | Leukocyte | https://doi.org/10.1038/s41587-020-0465-8 |
| GP5 | Increased | Megakaryocyte | Leukocyte | https://doi.org/10.1038/s41587-020-0465-8 |
| NRGN | Increased | Megakaryocyte | Leukocyte | https://doi.org/10.1038/s41587-020-0465-8 |
| TUBB1 | Increased | Megakaryocyte | Leukocyte | https://doi.org/10.1038/s41587-020-0465-8 |
| SPARC | Increased | Megakaryocyte | Leukocyte | https://doi.org/10.1038/s41587-020-0465-8 |
| RGS18 | Increased | Megakaryocyte | Leukocyte | https://doi.org/10.1038/s41587-020-0465-8 |
| MYL9 | Increased | Megakaryocyte | Leukocyte | https://doi.org/10.1038/s41587-020-0465-8 |
| GNG11 | Increased | Megakaryocyte | Leukocyte | https://doi.org/10.1038/s41587-020-0465-8 |
| AIF1 | Increased | Monocyte | Leukocyte | https://doi.org/10.3389/fimmu.2023.1223471 |
| NKG7 | Increased | NK | Leukocyte | https://doi.org/10.3390/ijms25073828 |
| GNLY | Increased | NK | Leukocyte | https://doi.org/10.3389/fimmu.2023.1223471 |
| FGFBP2 | Increased | NK | Leukocyte | https://doi.org/10.3389/fimmu.2023.1223471 |
| KLRF1 | Increased | NK | Leukocyte | https://doi.org/10.3389/fimmu.2023.1223471 |
| GZMH | Increased | NK | Leukocyte | https://doi.org/10.3389/fimmu.2023.1223471 |
| PRF1 | Increased | NK | Leukocyte | https://doi.org/10.3389/fimmu.2023.1223471 |
| KLRD1 | Increased | NK | Leukocyte | https://doi.org/10.3389/fimmu.2023.1223471 |
| GZMB | Increased | NK | Leukocyte | https://doi.org/10.3389/fimmu.2023.1223471 |
| SPON2 | Increased | NK | Leukocyte | https://doi.org/10.3389/fimmu.2023.1223471 |
| CD247 | Increased | NK | Leukocyte | https://www.sc-best-practices.org/cellular_structure/annotation.html |
| KLRB1 | Increased | NK | Leukocyte | https://doi.org/10.1038/s41587-020-0465-8 |
| KLRD1 | Increased | NK | Leukocyte | https://doi.org/10.1038/s41587-020-0465-8 |
| KLRF1 | Increased | NK | Leukocyte | https://doi.org/10.1038/s41587-020-0465-8 |
| GZMA | Increased | NK | Leukocyte | cellxgene |
| CDKN1C | Increased | Non-classical monocyte | Leukocyte | https://doi.org/10.3389/fimmu.2023.1223471;https://doi.org/10.1038/s41587-020-0465-8 |
| LRRC25 | Increased | Non-classical monocyte | Leukocyte | https://doi.org/10.3389/fimmu.2023.1223471 |
| TCF7L2 | Increased | Non-classical monocyte | Leukocyte | https://doi.org/10.3389/fimmu.2023.1223471 |
| LYN | Increased | Non-classical monocyte | Leukocyte | https://doi.org/10.3389/fimmu.2023.1223471 |
| IF130 | Increased | Non-classical monocyte | Leukocyte | https://doi.org/10.3389/fimmu.2023.1223471 |
| FCGR3A | Increased | Non-classical monocyte | Classical mon | https://doi.org/10.3390/ijms25073828 |
| FCGR3A | Increased | Non-classical monocyte | Leukocyte | https://doi.org/10.1038/s41587-020-0465-8 |
| MS4A7 | Increased | Non-classical monocyte | Leukocyte | https://doi.org/10.1038/s41587-020-0465-8 |
| IL3RA | Increased | pDC | Leukocyte | https://doi.org/10.3389/fimmu.2023.1223471 |
| TCF4 | Increased | pDC | Leukocyte | https://www.sc-best-practices.org/cellular_structure/annotation.html |
| IL3RA | Increased | pDC | Leukocyte | https://doi.org/10.1038/s41587-020-0465-8 |
| GZMB | Increased | pDC | Leukocyte | https://doi.org/10.1038/s41587-020-0465-8 |
| JCHAIN | Increased | pDC | Leukocyte | https://doi.org/10.1038/s41587-020-0465-8 |
| IRF7 | Increased | pDC | Leukocyte | https://doi.org/10.1038/s41587-020-0465-8 |
| TCF4 | Increased | pDC | Leukocyte | https://doi.org/10.1038/s41587-020-0465-8 |
| LILRA4 | Increased | pDC | Leukocyte | https://doi.org/10.1038/s41587-020-0465-8 |

| | | | | |
|---|---|---|---|---|
| CLEC4C | Increased | pDC | Leukocyte | https://doi.org/10.1038/s41587-020-0465-8; https://doi.org/10.3390/ijms25073828 |
| TRBC2 | Increased | T | Leukocyte | cellxgene |
| CD3D | Increased | T | Leukocyte | https://doi.org/10.3389/fimmu.2023.1223471; https://doi.org/10.1038/s41587-020-0465-8 |
| CD3G | Increased | T | Leukocyte | https://doi.org/10.3389/fimmu.2023.1223471; https://doi.org/10.1038/s41587-020-0465-8 |
| CD3E | Increased | T | Leukocyte | https://doi.org/10.3389/fimmu.2023.1223471; https://doi.org/10.1038/s41587-020-0465-8 |
| IL7R | Increased | T | Leukocyte | https://doi.org/10.3389/fimmu.2023.1223471 |
| TRAC | Increased | T | Leukocyte | https://doi.org/10.3389/fimmu.2023.1223471; https://doi.org/10.1038/s41587-020-0465-8 |
| DFNB31 | Increased | T | Leukocyte | https://doi.org/10.3389/fimmu.2023.1223471 |
| EOMES | Increased | CD8+ T | Leukocyte | https://doi.org/10.3389/fimmu.2023.1223471 |
| TIM3 | Increased | T | Leukocyte | https://doi.org/10.3389/fimmu.2023.1223471 |
| TCF7 | Increased | T | Leukocyte | https://doi.org/10.1038/s41587-020-0465-8 |
| CD21 | Increased | B memory | B naive | https://pmc.ncbi.nlm.nih.gov/articles/PMC6851823/ |
| IGHD | Increased | B naive | Leukocyte | https://doi.org/10.1038/s41587-020-0465-8 |
| FCRL1 | Increased | B naive | Leukocyte | https://www.sc-best-practices.org/cellular_structure/annotation.html |
| IGHM | Increased | B naive | Leukocyte | https://doi.org/10.1038/s41587-020-0465-8 |
| IL4R | Increased | B naive | Leukocyte | https://www.sc-best-practices.org/cellular_structure/annotation.html |
| PAX5 | Increased | B | Leukocyte | https://www.sc-best-practices.org/cellular_structure/annotation.html |
| CD82 | Increased | B memory | B naive | https://doi.org/10.1002/jcla.20001 |
| IGHA1 | Increased | B memory | B naive | https://doi.org/10.1002/jcla.20001 |
| IGHG1 | Increased | B memory | B naive | https://doi.org/10.1002/jcla.20001 |
| SARAF | Increased | CD4+ T | Leukocyte | |
| LTB | Increased | CD4+ T | CD8+ T | |
| LDHB | Increased | CD4+ T | CD8+ T | |
| MAL | Increased | CD4+ T | CD8+ T | |
| HES4 | Increased | Non-classical monocyte | Leukocyte | |
| FCN1 | Increased | Monocyte | Leukocyte | |
| HBB | Increased | Erythrocyte | Leukocyte | cellxgene |
| SLC25A37 | Increased | Erythrocyte | Leukocyte | cellxgene |
| BLVRB | Increased | Erythrocyte | Leukocyte | cellxgene |
| CTSS | Increased | Monocyte | Leukocyte | https://doi.org/10.3389/fimmu.2023.1223471 |
| FTL | Increased | Monocyte | Leukocyte | cellxgene |
| CLC | Increased | Granulocyte | Leukocyte | cellxgene |
| CPA3 | Increased | Granulocyte | Leukocyte | cellxgene |
| GATA2 | Increased | Granulocyte | Leukocyte | cellxgene |
| HDC | Increased | Granulocyte | Leukocyte | cellxgene |
| HPGDS | Increased | Granulocyte | Leukocyte | cellxgene |
